# Recapitulation of HIV-1 Env-Antibody Coevolution in Macaques Leading to Neutralization Breadth

**DOI:** 10.1101/2020.08.05.237693

**Authors:** Ryan S. Roark, Hui Li, Wilton B. Williams, Hema Chug, Rosemarie D. Mason, Jason Gorman, Shuyi Wang, Fang-Hua Lee, Juliette Rando, Mattia Bonsignori, Kwan-Ki Hwang, Kevin O. Saunders, Kevin Wiehe, M. Anthony Moody, Peter T. Hraber, Kshitij Wagh, Elena E. Giorgi, Ronnie M. Russell, Frederic Bibollet-Ruche, Weimin Liu, Jesse Connell, Andrew G. Smith, Julia DeVoto, Alexander I. Murphy, Jessica Smith, Wenge Ding, Chengyan Zhao, Neha Chohan, Maho Okumura, Christina Rosario, Yu Ding, Emily Lindemuth, Anya M. Bauer, Katharine J. Bar, David Ambrozak, Cara W. Chao, Gwo-Yu Chuang, Hui Geng, Bob C. Lin, Mark K. Louder, Richard Nguyen, Baoshan Zhang, Mark G. Lewis, Donald Raymond, Nicole A. Doria-Rose, Chaim A. Schramm, Daniel C. Douek, Mario Roederer, Thomas B. Kepler, Garnett Kelsoe, John R. Mascola, Peter D. Kwong, Bette T. Korber, Stephen C. Harrison, Barton F. Haynes, Beatrice H. Hahn, George M. Shaw

## Abstract

Neutralizing antibodies elicited by HIV-1 coevolve with viral Envs in distinctive patterns, in some cases acquiring substantial breadth. Here we show that primary HIV-1 Envs, when expressed by simian-human immunodeficiency viruses in rhesus macaques, elicited patterns of Env-antibody coevolution strikingly similar to those in humans. This included conserved immunogenetic, structural and chemical solutions to epitope recognition and precise Env-amino acid substitutions, insertions and deletions leading to virus persistence. The structure of one rhesus antibody, capable of neutralizing 49% of a 208-strain panel, revealed a V2-apex mode of recognition like that of human bNAbs PGT145/PCT64-35M. Another rhesus antibody bound the CD4-binding site by CD4 mimicry mirroring human bNAbs 8ANC131/CH235/VRC01. Virus-antibody coevolution in macaques can thus recapitulate developmental features of human bNAbs, thereby guiding HIV-1 immunogen design.

**One sentence summary:** Virus-antibody coevolution in rhesus macaques recapitulates developmental features of human antibodies.

## Introduction

A major roadblock to rational HIV-1 vaccine design is the lack of a suitable primate model in which broadly neutralizing antibodies (bNAbs) can be commonly induced and the molecular, biological and immunological mechanisms responsible for such responses studied in a reproducible and iterative fashion. Since most HIV-1 bNAbs identified to date have come from humans chronically infected by HIV-1, we hypothesized that one means to elicit such antibodies in primates might be by infecting Indian rhesus macaques (RMs) with simian-human immunodeficiency virus (SHIV) strains that bear primary or transmitted/founder (T/F) HIV-1 Envs that induced bNAbs in humans (*1–7*). SHIV infected RMs could then be employed to assess the potential of particular HIV-1 Envs to elicit bNAbs and to characterize the coevolutionary pathways of bNAb lineages and the cognate Env intermediates that elicited them (*2–8*), thereby serving as a molecular guide for rational vaccine design. Recent innovations in SHIV design (*9*) now make this experimental strategy testable.

HIV-1 bNAbs target one of several conserved regions on the native Env trimer, including the CD4-binding site (CD4bs), V2 apex, V3 high mannose patch, gp120/gp41 interface, fusion peptide, glycosylated silent face, and membrane-proximal external region (MPER) (*10, 11*). Such antibodies generally share certain features such as exceptional HCDR3 length, extensive somatic hypermutation, autoreactivity or unusual mechanisms for binding glycans or peptidoglycans (*10–13*). Long HCDR3s result from initial germline immunoglobulin gene rearrangements whereas the characteristic high frequencies of V(D)J mutations that lead to affinity maturation and neutralizing breadth result from persistent virus replication, epitope variation, and continued Env-Ab coevolution. In some subjects, cooperating antibody lineages that target the same epitope have been identified, and together they contribute to Env diversification and bNAb development (*4, 6*). A critical question in the HIV-1 vaccine field is if the relatively rare examples of high titer, broadly neutralizing antibodies that have been identified in a subset of chronically infected humans represent stochastic accidents of nature, not likely to be replicated by a vaccine, or if there are special properties of particular HIV-1 Envs that along with deterministic features of Env-Ab coevolution make HIV-1 vaccine development more plausible.

The current study is based on the premise that while Env diversity within HIV-1 group M is extraordinarily high, primary HIV-1 Envs [i.e., native Env trimers on infectious virions that allow for persistent replication and clinical transmission in humans (*1*)] are nonetheless constrained in their immediate evolutionary potential (*14–16*). This paradox – extreme Env diversity globally but constraints on immediate or near-term evolution in an individual – can be explained by competing evolutionary forces: positive selection imposed by immune evasion balanced by purifying (negative) selection acting to retain viral fitness. A single zoonotic transmission event ∼100 years ago gave rise to HIV-1 group M (*17, 18*), and virus has since replicated essentially continuously with a short virus generation time of ∼2 days (*19*). This rate equates to tens of thousands of error-prone viral replication cycles over a period of a century, amplified by complex phylogenies that arose in tens of millions of infected humans each with unique adaptive immune responses driving viral antigenic variation. This amplification has resulted in the extraordinary global HIV-1 diversity observed today (www.hiv.lanl.gov). At the same time, variation in amino acid sequence and secondary, tertiary and quaternary protein structure has been constrained by strict requirements for antibody evasion and retention of essential biological functions including efficient Env expression, processing and assembly, receptor and coreceptor binding, and membrane fusion. Other sequence requirements that relate to overlapping reading frames, RNA splicing and Rev binding have exerted additional constraints on *env* (*14*). Antibody avoidance is mediated by at least four interrelated Env properties – trimer exclusion (*20*), glycan shielding (*21*), epitope variation (*22*) and conformational masking (*23*) – that act in concert. Altogether, these features limit each primary Env’s immediate and short-term evolutionary trajectory. When introduced into a human as HIV-1 infection or into RMs as SHIV infections, each Env will therefore be uniquely restricted in its exposure of antigenic sites that are accessible to neutralizing antibodies and will favor distinct molecular pathways of escape from such antibodies, with each pathway exacting a different fitness cost from the virus. These considerations led us to hypothesize that SHIV infection of RMs will recapitulate HIV-1 infection of humans with respect to molecular patterns of Env-Ab coevolution, within bounds set by any particular Env sequence and the respective rhesus and human immunoglobulin gene repertoires that respond to it (*24*).

## Results

Some of the best studied examples of HIV-1 Env-Ab co-evolution have come from the analysis of HIV-1 infected human subjects CH505, CH848 and CAP256, who developed bNAb responses targeting the CD4bs, V3 high mannose patch and V2 apex of HIV-1 Env, respectively (*2–7*). CH505 and CH848 were each productively infected by a single transmitted/founder (T/F) virus, whereas CAP256 was infected by a single virus (CAP256PI, primary infection) and then superinfected by a second genetically-divergent virus (CAP256SU, superinfection) (*25*). The CAP256SU variant was thought to trigger the V2 bNAb lineage in this individual (*5*), so we constructed and analyzed a SHIV containing this Env (**Fig. S1**). SHIVs containing CH505, CH848 and CAP256SU Envs were modified at gp120 residue 375 to enhance binding and entry into rhesus CD4 T cells. The Env375 substitutions resulted in no discernible change in antigenicity or sensitivity to anti-HIV-1 NAbs compared with wild-type virus, but they were essential for replication in primary RM CD4+ T cells [**Fig. S1** and (*9*)]. We inoculated 22 RMs (**Table S1**) with SHIV.CH505 (n=10), SHIV.CH848 (n=6) or SHIV.CAP256SU (n=6) by the intravenous route and followed them for as long as 180 weeks for viral replication kinetics and persistence, the development of strain-specific (autologous) and broadly reactive (heterologous) NAbs, and patterns of Env sequence evolution. Eight of the animals were treated with anti-CD8 mAb at or near the time of SHIV inoculation in order to transiently deplete CD8^+^ cells, increase peak and set point viremia titers, and potentially enhance the induction of NAbs. One animal (RM5695) was repurposed from an earlier study involving HIV-1 gp120 protein immunization (*26*). This animal was inoculated with acute phase plasma from three SHIV.CH505 infected animals (RMs 6069, 6070, 6072) to increase early virus diversity.

### SHIV Replication *In Vivo* and Elicitation of Neutralizing Antibodies

**Figure S2** depicts the kinetics of virus replication and the development of autologous, strain-specific tier 2 NAbs as measured by plasma vRNA and inhibition of virus entry into TZM-bl cells (*21*). Acute virus replication kinetics were consistent for all SHIVs in all animals with peak viremia occurring about 14 days post-inoculation and reaching titers in the 10^5^-10^8^ vRNA/ml range (geometric mean = 4.2×10^6^). Peak vRNA titers were higher in anti-CD8 treated animals than in untreated animals (geometric mean titers of 2.6×10^7^ vs 1.5×10^6^, respectively; p<0.0001). Plasma virus load set points at 24 weeks post-infection varied widely between undetectable and 988,110 vRNA/ml with a mean for anti-CD8 treated animals of 248,313 ± 136,660 vRNA/ml compared with 8,313 ± 3509 vRNA/ml for untreated animals (p=0.002). Set point vRNA levels in human subjects infected with the corresponding HIV-1 CH505, CH848 or CAP256 strains were 81,968, 132,481 and 750,000 vRNA/ml, respectively. Compared with two human cohort studies of acute and early HIV-1 infection (*27, 28*), peak and setpoint vRNA titers in SHIV infected animals were comparable, although several macaques developed undetectable viral loads, which is rare in humans. Six macaques developed AIDS-defining events and were euthanized 36, 40, 60, 65, 89 or 136 weeks post-SHIV infection; four of these animals had been treated with anti-CD8 mAb (**Table S1**).

Autologous tier 2 NAb responses to the three SHIVs were detected as early as 10 weeks post-infection and peaked between 25 and 80 weeks with 50% inhibitory dilutions (ID_50_) between 0.05 (1:20 dilution) and 0.000125 (1:8,000 dilution) (**Fig. S2**). The kinetics of appearance and titers of autologous tier 2 Nabs that developed in response to SHIV infections were generally comparable to those observed in humans infected by viruses bearing the homologous Envs (*2, 7, 25*). Among all animals, there was a significant association between higher setpoint virus titers and higher autologous tier 2 NAb titers (Spearman correlation rank correlation coefficient r_s_=0.74, p<0.0001). Heterologous plasma neutralizing activity was assessed against a diverse 22-member global panel of tier 1a (n=3) or tier 2 (n=19) HIV-1 strains (*29–32*) (**Table S2**). All animals developed potent neutralizing responses to the three tier 1a viruses (geometric mean titer ID_50_ = 0.0004). Tier 1a viruses have “open” Env trimers that spontaneously expose linear V3 and conformational CD4 induced (CD4i) bridging sheet epitopes, thus explaining their extreme sensitivity to what are otherwise non-neutralizing antibodies. Such “non-neutralizing” antibodies that target linear V3 and CD4i epitopes are elicited in virtually all HIV-1 infected humans, thereby selecting for Envs with an open-closed equilibrium that strongly favors the closed configuration (*23, 33–35*). Tier 2 viruses typify primary or T/F viruses that are generally resistant to neutralization by heterologous plasma antibodies except for those that target one of the canonical bNAb epitope clusters (*1, 10, 11, 29*). Seven of 22 RMs in this study developed antibody responses that neutralized between 6 and 17 of the 18 heterologous HIV-1 tier 2 viruses in our test panel at titers between 0.05 and 0.0001, in addition to robust responses to the autologous virus strains (**Fig 1A**; **Table S2**). Two of these animals (RM5695 and RM6070) were infected by SHIV.CH505, two (RM6163 and RM6167) by SHIV.CH848, and three (RM40591, RM42056 and RM6727) by SHIV.CAP256SU. Neutralization breadth was first detected as early as 16 weeks post-SHIV infection in two animals and as late as 88 weeks post-SHIV infection in others. The remaining 15 animals showed either no or very limited, low titer neutralization breadth. These findings show that SHIVs bearing primary T/F Envs reproduce in RMs key features of early virus replication dynamics, virus persistence and NAb elicitation that are characteristic of HIV-1 infection in humans, including the potential to elicit bNAbs.

**Fig. 1.**
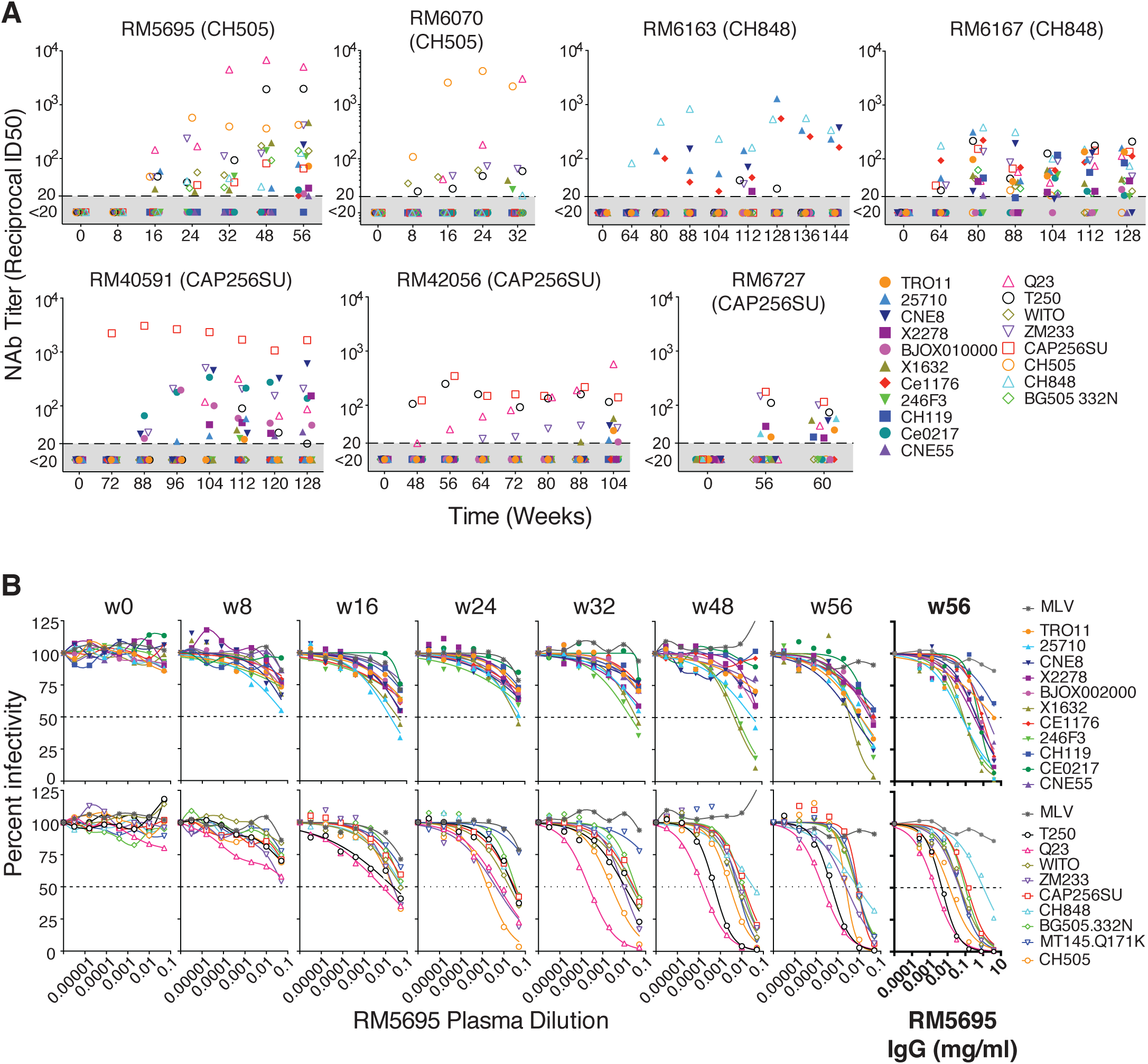
Broadly neutralizing antibodies in seven rhesus macaques. (**A**) RMs 5695 and 6070 were infected by SHIV.CH505, RMs 6163 and 6167 by SHIV.CH848, and RMs 40591, 42056 and 6727 by SHIV.CAP256SU. Neutralizing antibody titers (ID_50_, 50% inhibitory dilution) from longitudinal plasma specimens against 19 global HIV-1 strains (*29, 30, 86*) are depicted. (**B**) Neutralization curves for longitudinal plasma specimens and purified plasma IgG (highlighted bold) from RM5695 show the development of broad neutralization. MT145K.Q171K is a chimpanzee-derived SIVcpz strain that shares antigenic cross reactivity with HIV-1 in the V2 apex (*36, 134*). Dashed lines indicate 50% reduction in virus infectivity. MLV, murine leukemia virus.

**Figure 1B** highlights the kinetics, potency and breadth of neutralization in the plasma from the SHIV.CH505 infected animal RM5695 and identifies immunoglobulin G (IgG) as the active component. By 16 weeks post-infection, an autologous NAb response to SHIV.CH505 was detectable at an ID_50_ titer of 0.02 along with heterologous responses to viruses bearing HIV-1 25710, X1632, Q23, ZM233, T250 and WITO Envs at titers of 0.05 – 0.01. NAbs increased rapidly in breadth and titer thereafter. By week 48, bNAbs targeting Q23 and T250 jumped in titer to 0.0002 – 0.0005 and against X1632, 246F3, ZM233 and WITO to titers of 0.005. By week 56, bNAbs in the plasma of RM5695 neutralized 17 of 18 viruses in the heterologous HIV-1 test panel at titers ranging from 0.05 – 0.0002, along with a divergent tier 2 simian immunodeficiency virus strain (SIVcpz.MT145.Q171K) that shares selective antigenic cross-reactivity with HIV-1 in the V2 apex bNAb epitope cluster (*36*), at a titer of 0.006. IgG was purified by *Staphylococcal* Protein A/G column chromatography from RM5695 week 56 plasma and was assayed for neutralizing activity against the same 19 heterologous viruses: IgG concentrations between 0.002 mg/ml (corresponding to a ∼1:10,000 dilution of rhesus plasma) and 4 mg/ml neutralized 18 of the 19 viruses in a rank order similar to the polyclonal plasma (**Figure 1B**, rightmost panels). Neither plasma nor purified IgG neutralized control viruses pseudotyped with the murine leukemia virus (MLV) Env. Thus, anti-HIV-1 specific IgG accounted for all of the autologous and heterologous neutralizing activity in the RM5695 plasma. Of note, RM5695 IgG from week 56 plasma reached ID_90_ or ID_95_ thresholds against most viruses and exhibited steep inflections at the ID_50_ midpoint, indicating potent neutralization. Neutralization breadth detected as early as 16 weeks post-infection is unusual in HIV-1 infection but not unprecedented (*37*) and occurs most often with V2 apex bNAbs, likely because their activity depends more on long HCDR3s than on extensive somatic hypermutation. We show below that the bNAb activity in RM5695 plasma and its isolated IgG fraction as well as monoclonal bNAbs derived from RM5695, all targeted a bNAb epitope in the V2 apex that included the conserved lysine rich C-strand and N160 glycan. The kinetics of appearance, breadth, titers and potency of bNAbs elicited in the six other SHIV-infected RMs, including three animals (RM6070, RM40591 and RM42056) whose bNAbs also targeted the V2 apex C-strand, are summarized in **Figure 1A** and **Table S2**.

### Env Evolution in SHIV Infected Macaques and HIV-1 Infected Humans

To explore if SHIV Env evolution in RMs recapitulates that of HIV-1 in humans in a strain-specific manner, we analyzed Env sequences in the 22 SHIV infected animals over one to three years of follow-up and compared them with the evolution of the homologous Envs in humans infected by HIV-1. We used single genome sequencing (SGS) for this analysis, since it allows for retention of genetic linkage across intact viral genes and enables precise molecular tracking of sequence diversification from unambiguous T/F genomes (*1, 38–40*). We analyzed cDNA sequences derived from plasma virion RNA, since the half-life of circulating plasma virus is <1 hour (*19*) and that of cells producing most of this virus is <1 day (*19*). This short composite half-life (<1 day) reflects the lifespan of >99.9% of circulating plasma virus in individuals not receiving antiretroviral therapy (*41, 42*), making the genetic composition of the plasma virus quasispecies an exquisitely sensitive real time indicator of *in vivo* selection pressures acting on virus and virus-producing cells, including that exerted by NAbs, cytotoxic T cells (CTLs) or antiretroviral drugs (*21, 38, 40, 42, 43*). **Figure 2A** illustrates Env amino acid substitutions in virus from sequential plasma specimens from human subject CH505 and from six SHIV.CH505 infected RMs. By inspection and the algorithm-based statistical methods previously described [(*38, 39*) and *Supplementary Materials*], amino acid substitutions in Env were found to be nonrandomly distributed in patterns that evolved over time beginning as early as 4 weeks post-infection. Using LASSIE [Longitudinal Antigenic Sequences and Sites from Intra-Host Evolution (*44*)], a computational tool that systematically identifies amino acid differences in a set of evolved sequences that reach a predetermined frequency threshold compared to a founder sequence, we identified in subject CH505 fifteen Env residues where mutations altered the encoded amino acid in at least 75% of sequences at one or more subsequent time points. Thirteen of these 15 variable sites were shared in one or more monkeys and 10 sites were shared by at least half of the animals (**Fig. 2B**). Surprisingly, the substituted amino acids at many of these variable sites were identical in Env sequences from human and rhesus, including mutations that created potential N-linked glycans (PNGs) that filled “glycan holes” (**Figs. 2C, S3**) (*45*). A remarkably parallel CH505 Env evolutionary trajectory was observed between the human host and monkey RM6072 (**Fig. 2**). The conserved patterns of amino acid substitutions in CH505 Envs were very different from mutational patterns of Env sequences observed in the human subject CH848 and in RMs infected by SHIVs bearing the CH848 Env (**Fig. S4**). Twenty-nine variable sites were identified in Envs from the human subject CH848 and 16 of these were shared in SHIV.CH848 infected RMs, often with identical amino acid substitutions including some that altered the distribution of PNGs and glycan holes (**Fig. S5**). As a control, we compared patterns of Env evolution in CH848 infected RMs against selected sites in the CH505 infected human, and vice versa. Many fewer sites of shared evolution were observed in either discordant Env pairing (p < 0.01 for both comparisons) (**Fig. S6**). The Env strain-specific mutational patterns observed in humans and rhesus animals infected by CH505 and CH848 viruses were different still from the CAP256SU infected RMs in which 45 selected sites were identified, including 25 that were shared in more than one animal (**Figs. S7A,B and S8**). A comparable Env-wide analysis in the human subject CAP256 could not be performed because this individual was infected and then superinfected by two widely divergent viruses whose progeny underwent extensive recombination (*25*) (**Fig. S7C**). Still, we could identify common sites of positive selection in human and rhesus sequences, including a key mutation at residue 169 that led to virus escape from V2 apex C-strand targeted bNAbs (**Fig. S7D**). Overall, we identified 89 selected Env residues in CH505, CH848 or CAP256SU T/F viruses and 50 of these sites were shared among humans and RMs that were infected by viruses bearing the same Env strain. This result was markedly different for individuals infected by different virus strains where no more than 6 of these 89 variable sites were shared by any CH505, CH848 or CAP256SU pairing (p<0.0001) and only 14 sites were shared overall (p<0.0001). These findings thus show striking examples of strain-specific evolution of HIV-1 Env sequences in humans and RMs.

**Fig. 2.**
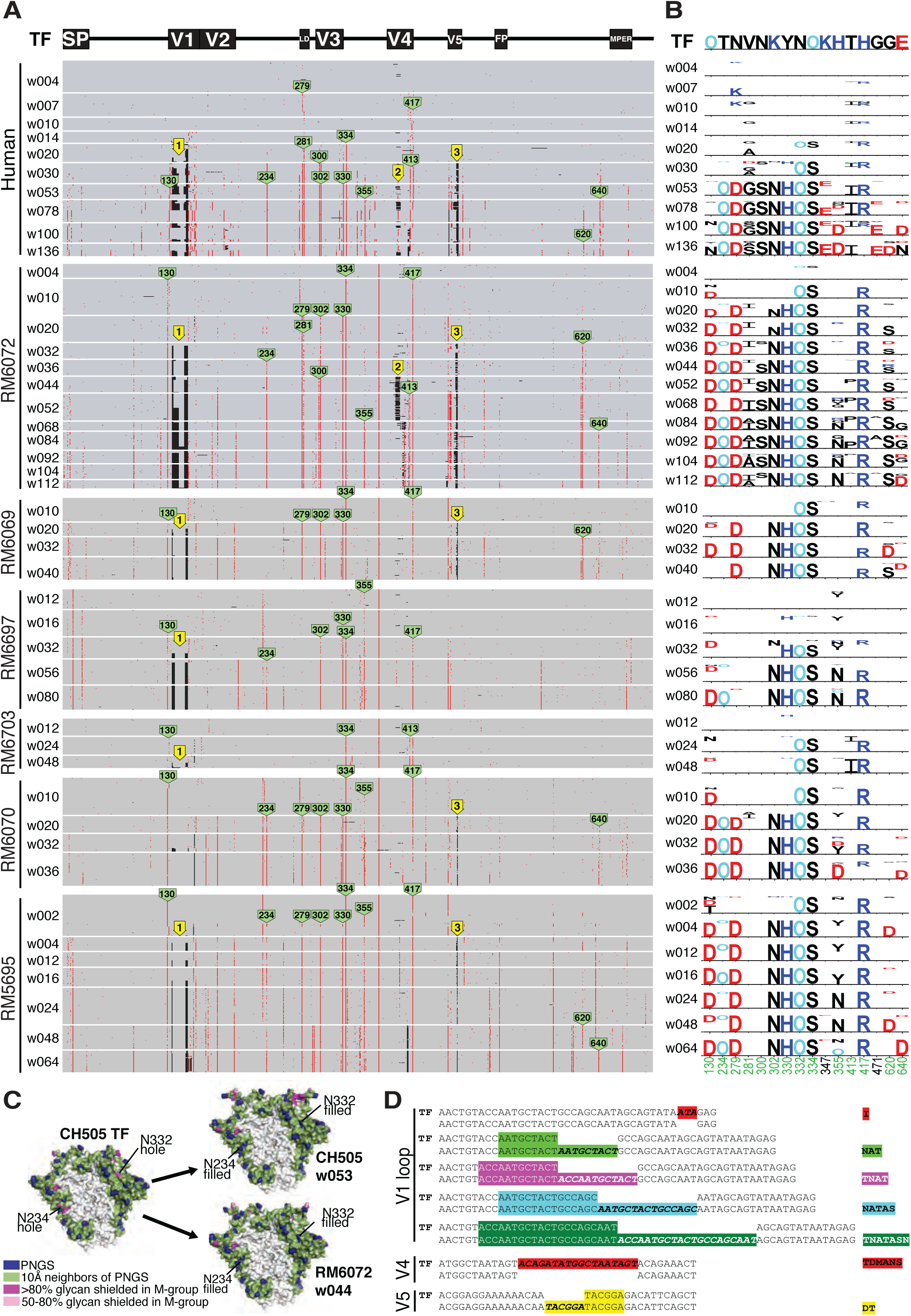
Env evolution in six SHIV.CH505 infected rhesus macaques recapitulates HIV-1 Env evolution in the human subject CH505. (**A**) A PIXEL plot (https://www.hiv.lanl.gov/content/sequence/pixel/pixel.html) (https://www.hiv.lanl.gov/content/sequence/HIV/HIVTools.html) of amino acid alignments of longitudinal HIV-1 Env sequences obtained by single genome sequencing of plasma virion RNA. Between 15-60 individual sequences are grouped by time point and amino acids are colored red to highlight mutations relative to the infecting CH505 transmitted/founder (T/F) strain. The image is highly compressed such that each row within a time block represents a single sequence, and each column represents an amino acid position in the alignment; thus each pixel represents a single amino acid that is colored grey if it matches the T/F sequence, red if it is a point mutation, and black if it is an insertion or deletion (INDEL) relative to the T/F virus. All SHIV sequences differ from the T/F at position 375, reflecting the SHIV design strategy that enables replication in rhesus macaques (*9*). Green tags indicate amino positions (HXB2 numbering) that are mutated in both human and rhesus. Yellow tags indicate three sites of identical INDELS observed in both human and rhesus. (**B**) LASSIE analysis (*44*) of the same longitudinal Env sequences was used to characterize mutations under positive selection. If a T/F amino acid was retained in <25% of the sequences in the human CH505 infection, the site was considered to be under selective pressure and tracked in all hosts. The height of each amino acid mutation is proportional to its frequency at the respective time point. Red, dark blue and black indicate acidic, basic and neutral residues. “O” indicates asparagine (N) embedded in an N-linked glycosylation motif. Numbers indicate residue positions (HXB2 numbering). Green numbers indicate mutations that reached 75% frequency in the human and at least one animal. (**C**) The area that is typically glycan shielded on the surface of the Env protein is shown in green (*45*). Magenta and pink highlight areas that are shielded in most Envs but are exposed in the CH505 T/F and referred to as “glycan holes”. Two glycan holes in the CH505 T/F were filled by the addition of NXS/T motifs at positions 234 and 332 over time in all of the monkeys (see **Fig. S3**). (**D**) INDELs in hypervariable loops arose in the first year of infection in the human host and were repeated in one or more rhesus macaques (**Fig. S9**). Insertions in HIV hypervariable regions were usually a direct sequence repeat, as shown immediately beneath the T/F sequences.

Another type of Env variation common in HIV-1 infection that can affect immune recognition results from insertions and deletions (Indels). Indels occur as a consequence of polymerase slippage and template switching during reverse transcription of the retroviral genome (*46, 47*). Indels are most commonly observed in the surface-exposed variable loops 1, 2, 4 and 5, which can better accommodate changes in sequence length than structural or enzymatic elements of the viral proteome, this was the case for the human subject CH505 and macaques infected by SHIV.CH505 (**Fig. 2A**). Remarkably, we found multiple examples of identical Indels between human and rhesus CH505 sequences and among CH505 sequences from different rhesus animals (**Figs. 2D and S9**). A total of six distinct V1 Indels, three V4 Indels and one V5 Indel were identified in sequences from human subject CH505 that were replicated exactly in sequences from one or more monkey. Two of these Indels were found in the human and in all six monkeys. Additional identically replicated Indels in V1 and V5 were identified only in monkeys. Indels were also observed in sequences from the human subject CH848 (**Figs. S4,S10**). An exact replica of all but one of these Indels was present in one or more macaque. Additional Indels were replicated in multiple animals (**Fig. S10**). None of the Indels in CH505 sequences were found in CH848, and neither set was found in CAP256SU sequences. In both human and rhesus, Indels generally lengthened or shortened variable loops in a direction that approached the median lengths of globally circulating viruses (**Figs. S9,S10**). We performed statistical analyses to determine the likelihood that the conserved sets of Env specific Indels that we observed in different individuals could occur by chance and that likelihood was estimated to be vanishingly small (see Extended Discussion in Supplement).

### NAbs select for mutations in Env

#### CH505

To examine if NAb-mediated selection was a primary driving force in the Env-specific evolutionary patterns that we observed, early strain-specific NAb responses and later bNAb responses were mapped using site-directed mutagenesis to introduce observed mutations into neutralization sensitive autologous and heterologous Envs. We then tested these mutant Envs, compared with their wild-type counterparts, for sensitivity to polyclonal plasma and mAbs isolated from the infected RMs or human subjects. Variable sites in the CH505 Env (**Figs. 2A,B**) were interrogated alone or in combination for their effects on Env sensitivity to autologous antibodies. Aside from variable residues in the leader sequence of gp160 that generally represent CTL escape mutations (*38, 43*), CTL epitope reversions to global consensus at or near residue 417 (*4*), and mutations at residues 300, 620 and 640 that occurred inconsistently and late, most of the variable residues were found to represent escape mutations from autologous NAbs (**Fig. S11A**). These included residues 234 and 334 where mutations restored PNGs that filled T/F glycan holes (**Fig. S3**), loop D residues 279 and 281 involved in CD4 binding, CD4 contact residues in V5 (460/ΔV5), and residue 130. The temporal appearance of these NAbs coincided with the appearance of phenotypically demonstrable NAb escape mutations in Env (**Fig. 2A,B**). These results were corroborated by neutralization patterns of V3 targeted mAbs DH647 and DH648 and CD4bs targeted mAbs DH650.UCA and DH650 (all isolated from the SHIV.CH505 infected RM6072) and the human CD4bs targeted mAbs CH235.UCA, CH235.IA3 and CH235.9 isolated from human subject CH505 (*6*) (**Figs. S11B,C**). Human subject CH505 developed two cooperative lineages of CD4bs antibodies that targeted epitopes that included loop D residues 279-281 and residues in V5, eventually leading to neutralization breadth by both antibody lineages (*4, 6*). In RM6072, a mAb lineage, termed DH650, was isolated that targeted these same epitopes but it never developed neutralization breadth, a finding for which we found a structural explanation (see below).

In addition to the strain-specific NAb responses that we identified in SHIV.CH505 infected RMs, we found in two CH505 animals (RM5695 and RM6070) neutralizing antibodies that targeted heterologous tier 2 viruses (**Fig. 1**, **Table S2**). In both animals, these bNAbs were first detected at 16 weeks post-infection and their development was temporally associated with the appearance of mutations in the Env V2 apex, including residues 166 or 169 (**Figs. 3A**) that are common contact residues of human V2 apex bNAbs (*5, 8, 48*). A minor fraction of sequences lost PNGs at 156 and 160. By 24 weeks post-infection in RM5695 and 32 weeks post-infection in RM6070, most circulating virus contained mutations at residues 166 or 169; by 36-48 weeks post-infection, all sequences were mutant at one or the other of these positions. We corroborated this single genome sequence evidence of strong virus selection in the central cavity of the V2 apex of RM5695 by next generation sequencing, which revealed that 99.3% of 10,000 sequences sampled between weeks 48 and 64 post-infection contained mutations at residues 166 or 169. When these residues were mutated in the wild type versions of heterologous primary virus strains T250, Q23, MT145K, 246F or BG505 that were otherwise neutralized by RM5695 and RM6070 plasma, neutralization was abrogated (**Figs. 3C,S12**). Neutralization of these heterologous strains was variably dependent on the glycan at N160 (**Fig. S13**), similar to what has been reported for V2 apex bNAbs in subject CAP256SU (*49*). These findings indicated the presence of potent V2 apex C-strand targeted bNAbs in RM5695 and RM6070. In summary, the mapping of autologous and heterologous Nab responses in RMs infected by SHIV.CH505 indicated that most mutations in Env that could not be ascribed to CTL selection were the result of NAb selection.

#### CH848

We also examined Env mutations shared between SHIV.CH848 infected RMs and the HIV-1 infected CH848 human subject as potential sites targeted by NAbs. We focused first on deletions in V1 that were similar, and in some cases identical, in viruses from human and rhesus (**Figs. S4,S9**). In the human subject CH848, an early strain-specific NAb response targeted V1, as demonstrated previously by mapping both polyclonal NAb responses and a mAb, DH475, isolated from subject CH848 (*7*). The circulating virus quasispecies targeted by this early V1 NAb response in subject CH848 escaped by deleting ∼10 amino acids in V1, which enabled the development of the V3 high mannose patch targeted bNAb lineage DH270 (*7, 50*). Remarkably, the same sequence of events occurred in SHIV.CH848 infected animals RM6163 and RM6167: V1 was first targeted by an early autologous NAb that we identified in plasma of RM6167 and in a six-member lineage of CH848 strain-specific mAbs isolated from RM6163 (**Fig. S14**). These polyclonal and monoclonal antibodies exhibited V1 specificity similar to that of the human DH475 mAb (*7*). Subsequently, deletions occurred in V1 sequences from RMs 6163 and 6167 (**Figs. S4,S10**), followed in both animals by the development of bNAbs (**Fig. 1A**, **Table S2**). We mapped the target epitopes of these polyclonal bNAb responses to the glycan at residue N332 and the Gly-Asp-Ile-Arg (GDIR) motif at residues 324-327 by introducing mutations into

heterologous Envs that were otherwise neutralized by plasma from RMs 6163 and 6167 (**Fig. 4B**). Coincident with the development of V3 glycan targeted bNAbs, we observed escape mutations at residues 332-334 and GDIR residues 324-327 in the evolving virus quasispecies of both RMs (**Fig. 4A**). Like the prototypic human V3 glycan bNAbs DH270, PGT121 and PGT128, the bNAbs that we identified in RMs 6163 and 6167 were strictly N332 dependent, and mutations in spatially-associated surface residues V295, N301, N138 and N133 had similar effects on neutralization potency of both rhesus and human bNAbs (**Figs. 4B,C**).

**Fig. 4.**
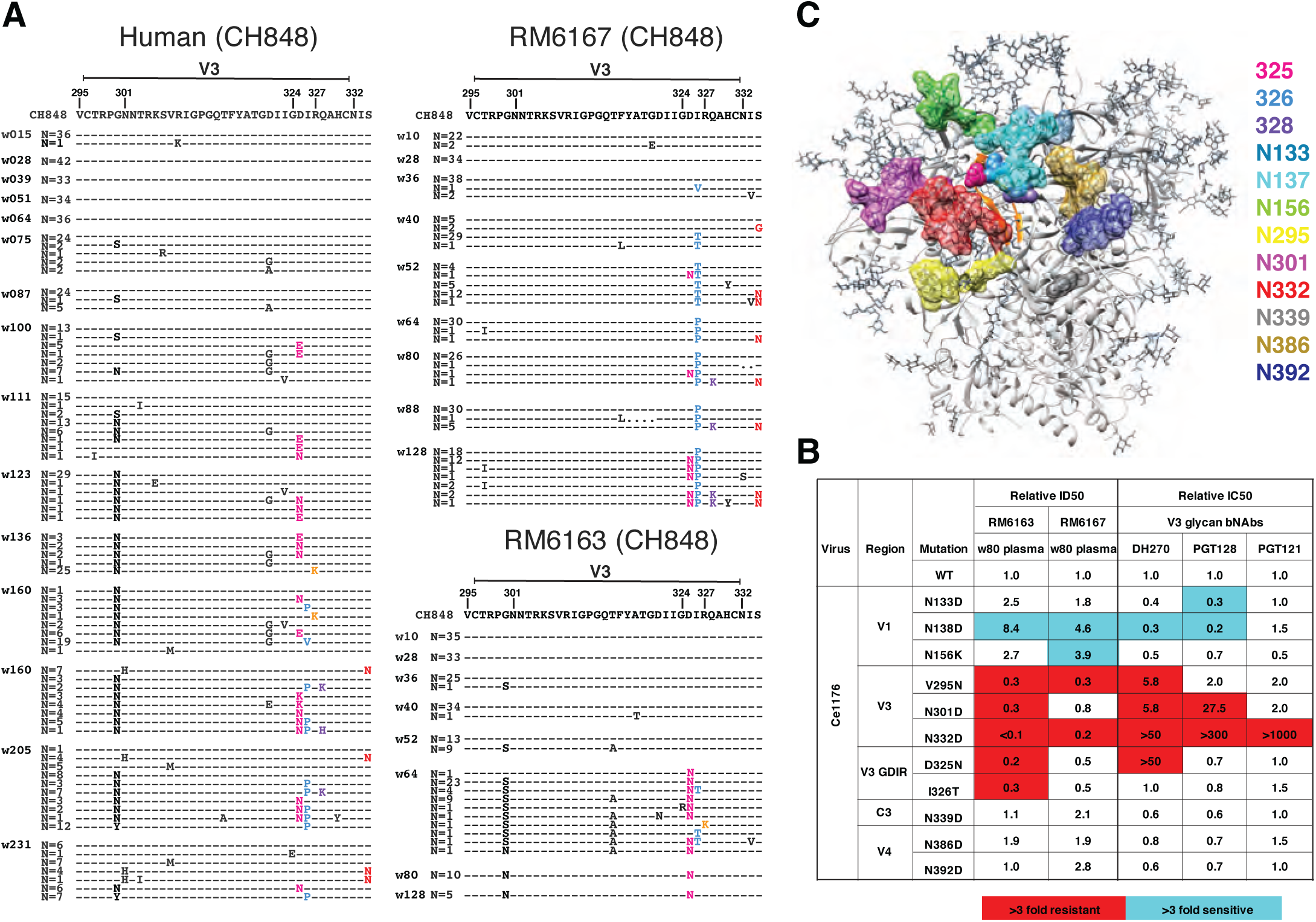
bNAb responses in two rhesus macaques map to the V3 glycan high mannose patch. (**A**) Sequential Env sequences from human subject CH848 and SHIV.CH848 infected RMs 6167 and 6163 showed selection and fixation of mutations in the GDIR motif, with fewer mutations eliminating the PNG site at 332 in RM6167. Similar mutations occurred in the human subject CH848. (**B**) Heterologous neutralization of Ce1176 Env-pseudotyped virus by RMs 6163 and 6167 plasma is reduced (expressed as fold-change from wildtype) by mutations at N332D and V295N, consistent with some human V3 glycan bNAbs. Neutralization of GDIR mutants is reduced by 2-5-fold. Elimination of a glycan at position N138 enhances neutralization in RMs 6163 and 6167 and for some human V3 glycan bNAbs. (**C**) V3 glycan bNAb escape mutations are displayed on the 5FYL structure of BG505.N332 SOSIP highlighting their close spatial proximity.

#### CAP256

In the human subject CAP256, recombination between PI and SU lineage sequences (**Fig. S7C**) precluded a gp160-wide Env analysis. We focused instead on mutations in and near the V2 C strand, since the human subject and two of six RMs (40591 and 42056) developed V2 apex targeted bNAbs (**Fig. 1A and Table S2**). In both human and RMs, very similar patterns of Env V2 sequence evolution occurred (**Fig. 3B**). These mutations included substitutions at the canonical residues 160, 166, 169 and 171 shown to be contact residues for other prototypical human V2 apex bNAbs (*5, 8, 51-53*). When introduced into heterologous neutralization-sensitive Envs, mutations at 166, 169 and 171 abrogated neutralization by plasma from RMs 40591 and 42056 as it did for control human bNAbs PGT145 and VRC26.25, the latter having been isolated from subject CAP256 (*49, 53*) (**Fig 3C**). In RMs 40591 and 42056, we observed that V2 apex targeted antibodies were variably dependent on binding to the glycan at Env residue 160 for neutralizing activity, a pattern that was similar to antibodies from animals RM5695 and RM6070 and the human subject CAP256 (*49*). This variable dependence on N160 for bNAb activity is further shown in **Fig. S13** and is discussed in Supplementary Materials. Overall, the patterns of Env sequence evolution reported here for SHIV-infected macaques, and previously in humans (*21, 38-40, 43*), highlight the exquisite sensitivity of dynamic measurements of localized sequence variation as an indicator of epitope-specific adaptive immune responses.

**Fig. 3.**
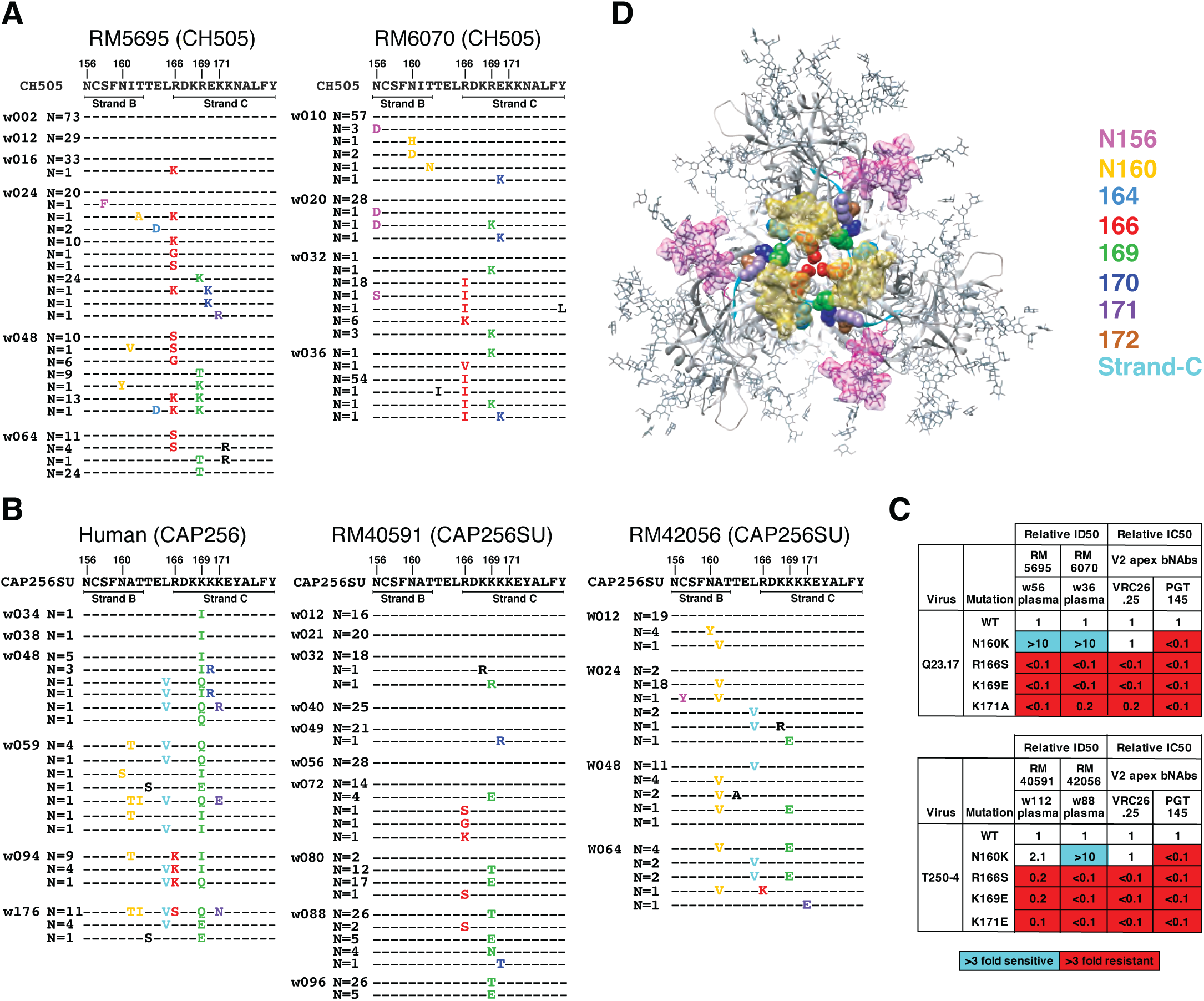
bNAb responses in four rhesus macaques map to the V2 apex. (**A**) Single genome sequencing (N = number of sequences; w = weeks post-SHIV infection) in SHIV.CH505 infected animals RM5695 and RM6070 reveals selection and fixation of mutations in strand C and additional mutations that eliminate PNG sites at 156 and 160. (**B**) Sequential Env sequences from human subject CAP256 and SHIV.CAP256SU infected RMs 40591 and 42056 showed selection and fixation of mutations in strand C, with fewer mutations eliminating the PNG site at 160. (**C**) Heterologous neutralization of Q23.17 and T250-4 Env-pseudotyped viruses by rhesus plasma is drastically reduced (expressed as fold-change from wildtype) by mutations at residues 166, 169 and 171 similar to human V2 apex bNAbs. Enhanced neutralization against the N160K mutants is illustrated in **Fig. S13** and further described in the Supplement. (**D**) Escape mutations from rhesus V2 apex bNAbs are displayed on the 5FYL structure of BG505.N332 SOSIP.

### Homologous B Cell Responses in Humans and Macaques

The observation that homologous Envs evolved in similar molecular patterns in humans and macaques could be explained by limited numbers of antigenic sites accessible for antibody binding and restricted pathways of virus escape. In addition, homologous human and rhesus germline B cell receptors could favor binding to common HIV-1 Env epitopes and follow similar patterns of Ab-Env coevolution. We explored the latter possibility by isolating and characterizing neutralizing mAbs from two RMs that were infected by SHIV.CH505. We selected these animals for study because both exhibited favorable virus replication kinetics, comparable early NAb responses, and an overall pattern of Env evolution similar to that in the human subject CH505; nonetheless, one animal (RM5695) developed bNAbs and the other (RM6072) did not. We asked what might be the genetic and structural similarities and dissimilarities between the NAbs elicited in these animals.

#### CD4bs-directed antibodies

Two broadly neutralizing mAb lineages (CH235 and CH103) targeting the HIV-1 CD4bs were previously isolated from the human subject CH505 and their evolutionary pathways from germline receptors to mature antibodies determined (*2, 4, 6*). In animal RM6072, serological analysis of early plasma samples suggested the presence of CD4bs targeted Abs based on selective Ab binding to CH505 T/F gp120 and resurfaced Env core but not to their isogenic Δ371I mutants, which do not bind CD4 (**Fig. S15A**). Additional evidence of CD4bs targeted Abs in the plasma of this animal included competitive blocking of soluble CD4 (sCD4) and CH235 and CH106 mAbs (**Fig. S15B**). We sorted memory B cells from RM6072 from longitudinal timepoints 20, 24 and 32 weeks post-SHIV infection and selected cells that reacted with CH505 T/F gp120 but not with the Δ371I mutant. We isolated a 15-member B cell clonal lineage designated DH650 (**Figs. S15C,D**) that selectively bound the autologous CH505 T/F Env gp120 but not the Δ371I mutant (**Figs. S15E**). Some of these mAbs competed with sCD4 and CH235 and CH106 mAbs for binding CH505 T/F gp120 (**Figs. S15F**). Mature DH650 lineage mAbs, but not the inferred germline UCA, bound CH505 T/F gp140 SOSIPv4.1 trimer (**Fig. S16A**). Most members of the DH650 lineage neutralized a glycan-deficient mutant of CH505 Env (gly4) and two-thirds of them neutralized the wildtype CH505 T/F strain (**Fig. S16B**). The DH650 UCA neutralized neither. None of the DH650 lineage mAbs neutralized heterologous viruses (**Fig. S16C and Table S3**).

The immunogenetic features of the DH650 lineage mAbs suggest how they recognize HIV-1 Env. The lineage comes from V(D)J recombination of the macaque V_H_1-h gene (**Fig. S15D**), which is 91% similar to an orthologous human gene V_H_1-46 used by the CD4bs bNAbs CH235 and 8ANC131 (*6, 54*). DH650 antibodies share key VH residues with CH235 (**Fig. S17A**), which were shown previously to be contact sites with gp120 Env (*6, 55*). They included residue 57, which in both the CH235 UCA and DH650 UCA underwent affinity maturation to R57, which is important for CH235 bNAb activity and is shared among CD4bs bnAbs using V_H_1-46 (*6, 55*). We found this N57R substitution in DH650 to be essential for binding to CH505 T/F Env (**Fig. S17B**). Other DH650 V_H_1-h gene residues that we found to be important for Env binding included N35, Q62 and R72 (**Figs. 5, S17C,D**). A distinguishing feature of DH650 lineage antibodies was the IGV*k*2 light chain, which has an exceptionally long LCDR1 of 17 amino acids (**Fig. S17E**) that we explored by structural studies.

The crystal structure of DH650 bound to the gp120 Env core of the CH505 T/F virus showed that its interactions with the gp120 CD4bs closely resembled those of the human CD4bs mAbs CH235, 8ANC131 and VRC01 (**Fig. 5A-D, Table S4**). This similarity included conserved HCDR2-mediated CD4-mimicry and coordination of Env Asp368 by Arg72. An important difference between the rhesus and human antibody lineages was in the light chains (DH650, macaque IGV*k*2; CH235, human IGVλ3), in which the LCDR1 of the DH650 light chain was six residues longer than its CH235 counterpart (**Fig. S17E)**. The structure showed that in the Env-Ab complex, the CH505 gp120 loop D had undergone a conformational change to accommodate the longer DH650 LCDR1. As we show elsewhere (Chug et al., to be submitted), this shift could occur only because of the absence of a commonly found glycan at gp120 position 234 in the CH505 TF virus. Moreover, addition of that glycan, which occurred in both the human donor and RM6072 by 30-36 weeks post infection, conferred resistance to DH650 (**Fig. S11C**) and likely eliminated selective pressure in the monkey to enforce deletions in LCDR1. Thus, infection of RM6072 with SHIV.CH505 expanded a B-cell clone bearing an antigen receptor encoded by the RM VH1-h gene segment that is orthologous to the human VH1-46 gene. This B-cell lineage underwent affinity maturation, including selection for a critical R57 VH mutation that is also found in the human CH235 bNAb lineage. Zhou et al. have shown that maturation of VRC01-class CD4bs bnAbs generally includes deletions in LCDR1 or mutations to glycine that confer flexibility (*56*). Evolution of the DH605 lineage in RM6072 failed to include deletions or flexibility in LCDR1 and hence neutralization breadth did not develop.

**Fig. 5.**
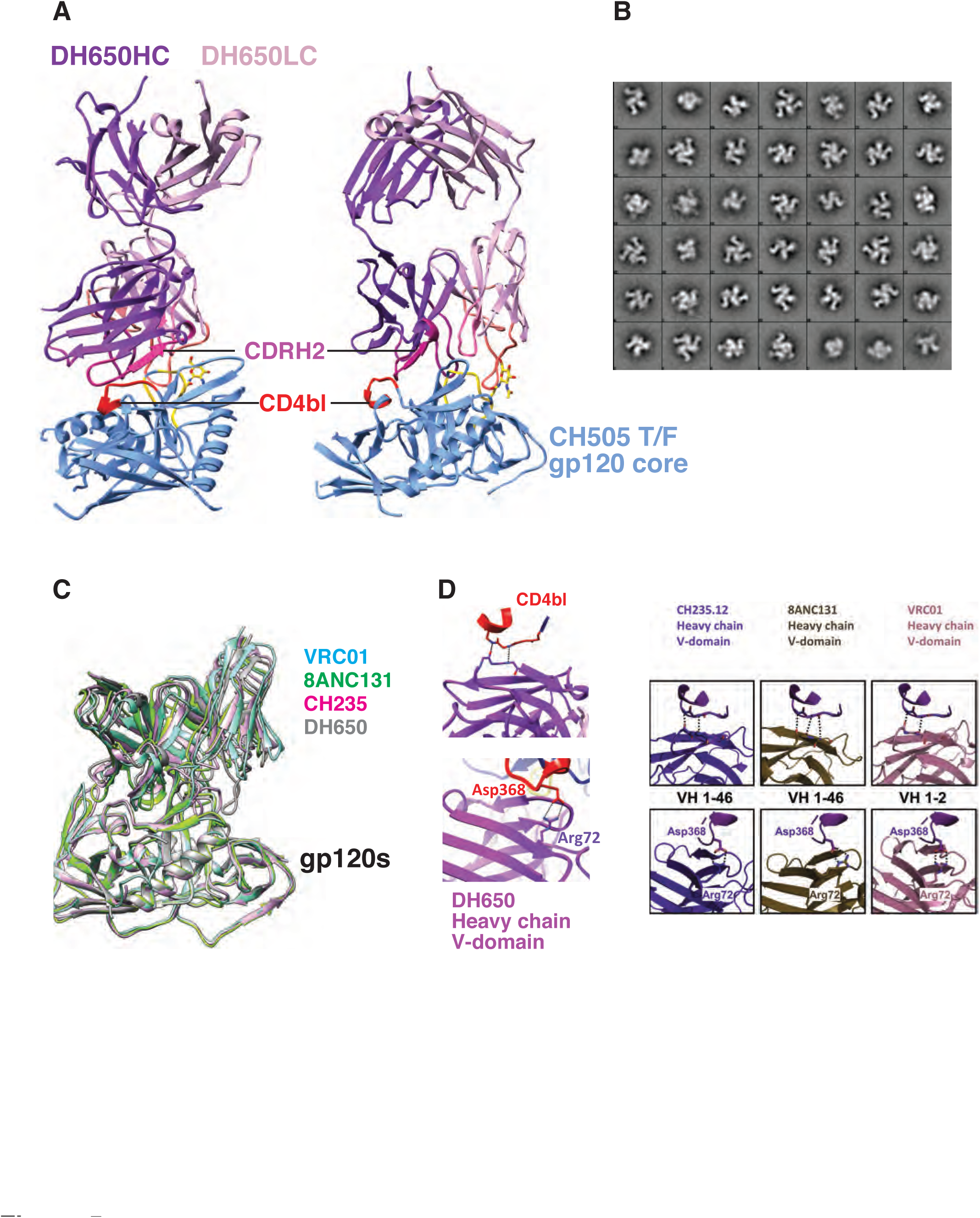
Crystal structure of Fab DH650 bound to CH505 gp120 core mimics human CD4bs antibodies. (**A**) Ribbon representation of Fab DH650 heavy chain (purple), light chain (pink), CHF505 gp120 core (blue), CDRH2 (magenta) and CD4 binding loop, bl (red). (**B**) Representative 2D class averages from EM images of negatively stained DH650-CH505 DS-SOSIP trimer. (**C**) Superposition of Fab DH650-CH505 gp120 complex with other CD4 binding site Fab-gp120 complexes, aligned on the gp120 core. Complex of DH650 in grey, VRC01 (PDB-3NGB) in cyan, CH235.12 (PDB-5F96) in magenta and 8ANC131 (PDB-4RWY) in green. Constant regions are omitted. (**D**) Recognition of CD4 binding loop of CH505 gp120 by heavy-chain variable domain of DH650 (top left) and coordination of Asp368 by Arg72 of DH650 (bottom left). This pattern of CD4 binding-loop recognition is similar to that of three other prototypic human CD4bs bNAbs (bottom right) (*6*).

#### V2 apex-directed antibodies

RM5695, infected with SHIV.CH505, developed broadly neutralizing antibodies which, based on Env escape patterns *in vivo* and neutralization phenotypes of site-directed Env mutants (**Fig. 3**), targeted the V2 apex. We used an unbiased FACS strategy to isolate 20,000 individual memory B cells from blood mononuclear cells 65 weeks post-SHIV infection and expanded these cells in 384-well plates. Culture supernatants were screened for neutralizing activity against two heterologous virus strains (T250-4 and Q23.17). Five wells scored positive and paired heavy and light chain immunoglobulin genes were successfully amplified from cellular mRNA from four of them (**Figs. 6,S18**). All four rhesus mAbs belonged to a single lineage that we designated RHA1.V2 (<u>R</u>hesus <u>H</u>IV <u>A</u>ntibody <u>1</u> targeting a <u>V2</u> epitope, with lineage members RHA1.V2.01-04). The IGVH and IGVL genes of the four RHA1.V2.01-04 mAbs were closely related with maximum nucleotide diversities of 5.3% and 3.9%, respectively. We employed NextGen sequencing to characterize immunoglobulin transcripts expressed by naïve IgD+IgM+IgG-B cells from RM5695 and used IgDiscover (*57*) and a recent database of RM Ig alleles (*24*) to identify a personalized immunoglobulin gene repertoire (**Table S5**). From this analysis, we determined the germline origins of the mature RHA1 mAbs to include a novel IGVH4 allele, IGHV4-ABB-S*01_S8200 (**Fig. S18A**), and IGλV1-ACN*02 (**Fig. S18B**). Somatic hypermutation within the lineage was modest, with nucleotide divergence from germline of 7.1 - 8.5% for V_H_ and 6.2 - 6.6% for V_L_ (**Fig. 6A**). These values are comparable to some human V2 apex bNAbs, which typically have lower frequencies of VH mutations than members of other human bNAb classes. The mature rhesus bNAb heavy chains contained a two amino acid insertion within HCDR1 (**Figs. S18A,19A**) and a 24 residue long HCDR3 (IMGT numbering) that was derived from rearrangement of the V_H_4-ABB-S*01_S8200, D_H_3-9 and J_H_2-P genes plus six non-templated amino acids (**Fig. S18A,19B**). This HCDR3 was rich in aromatic and negatively charged residues, like HCDR3s of human V2 apex bNAbs (**Fig. S19C**). Despite this long HCDR3, the mature rhesus bNAb RHA1.V2.01 was not auto- or polyreactive (**Fig. S20**). The HCDR3 of the RHA1 lineage antibodies was similar in length to the human V2 apex bNAb PCT64-35S (24 vs 25 amino acids, respectively), and the two broadly neutralizing mAbs contained conserved motifs within their respective HCDR3s including a negatively charged “DDY” segment (**Fig. S19C**), which in PCT64 was shown to be tyrosine-sulfated and to interact with positively charged V2 apex-hole residues of Env (*8, 58*). All four rhesus mAbs were tested for neutralization against the 19-member global panel of tier 2 viruses and showed similar patterns of reactivity, neutralizing 15 - 17 strains (**Fig. S21**). One antibody (RHA1.V2.01) was tested for neutralization against a 208 member global virus panel and was found to neutralize 102 heterologous virus strains, or 49%, at a maximum concentration threshold of 50 µg/ml (**Figs. 6B,S22**). Neutralization of heterologous virus strains depended on Env residues N160, R/K166 and R/K169, with partial dependence on K171 (**Figs. 6C,S23**). This precise pattern of neutralization sensitivity to N160, R/K166 and C-strand residue R/K169 and K171 mutations was shared by the human V2 apex bNAbs PCT64-35S and PGT145 but was different from that of PG9, VRC26.25 and CH01. CH505 Envs that evolved in RM5695 *in vivo* coincident with the development and maturation of RHA1 lineage antibodies showed evidence of strong selection at residues 166 and 169 (**Fig. 3A**). Introduction of these mutated residues into the CH505 T/F Env resulted in loss of neutralization sensitivity to RHA1 mAbs (**Fig 6D**).

**Fig. 6.**
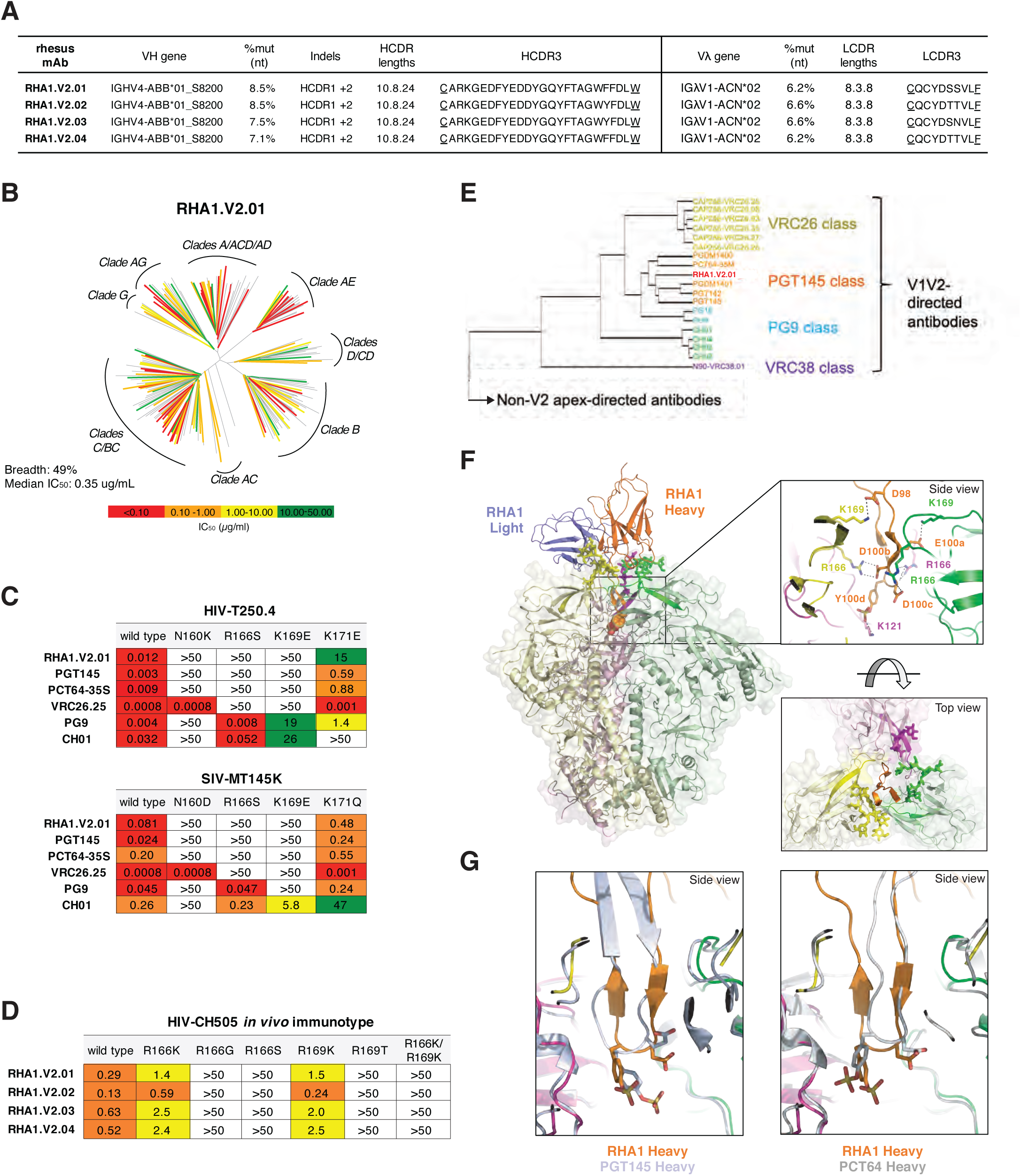
Rhesus bNAb lineage RHA1 targets the V2 apex and a sulfated tyrosine shows precise chemical mimicry to human bNAbs PCT64 and PGT145. (**A**) Immunogenetic characteristics of four RHA1 lineage broadly neutralizing monoclonal antibodies. A key feature is the 24 amino acid long CDRH3 that contains an acidic EDDY core motif. (**B**) Neutralization breadth and potency of RHA1.V2.01 against a 208 strain global virus panel. The dendrogram depicts phylogenetic relatedness of the HIV-1 Envs tested. (**C**) Neutralization expressed as IC_50_ (µg/ml) of wildtype heterologous viruses and their V2 apex mutants by RHA1.V2.01 and by prototypic human V2 apex bNAbs. Like most human V2 apex bNAbs, RHA1.V2.01 is strictly dependent on N160 and positively charged residues at 166 and 169. (**D**) Neutralization of CH505 T/F (wildtype) virus and C-strand variants, or “immunotypes,” that evolved *in vivo* in RM5695. Predominant mutations at 24 wks post-SHIV infection in RM5695 were R166K or R169K; at 48 wks R166S, R166G, R169T, and R166K plus R169K were prevalent; at 64 wks R166S or R169T became fixed (see Fig. 3A). Panel **D** shows progressive loss in neutralization sensitivity to RHA1 bNAbs by the evolving CH505 Envs, beginning with CH505 T/F (most sensitive), CH505.R166K or R169K (intermediately sensitive), and ending with CH505.R166G, R166S, R169T or R166K+R169K (all resistant). Results are expressed as IC_50_ (µg/ml). (**E**) Neutralization fingerprint for RHA1.V2.01 shows it to cluster within the PGT145 class. (**F**) Cryo-EM structure of RHA1.V2.01 in complex with BG505 DS-SOSIP at 4-Å resolution. Inset (top) highlights electrostatic contacts of the HCDR3 with Env protomers, including interactions of the tyrosine-sulfated 100d residue with Env K121. Inset (bottom) shows the trimer apex cavity highlighting glycans at N160 and the C-strands. (**G**) Alignment of gp120 from the complex trimer structure with RHA1.V2.01 Fab to trimer complexes with human Fabs PGT145 (PDB-5V8L) and PCT64 (modeled with PDB-6CA6 fit to EMD-7865) reveals alignment of tyrosine sulfated residues within the respective HCDR3 tips, which insert into the hole at the V2 apex of the Env trimer.

#### Bioinformatical comparisons of human and RM V2-apex bnAbs

We carried out a more thorough bioinformatics analysis of similarities and differences between rhesus mAb RHA1.V2.01 and prototypic human bNAbs. Neutralization “fingerprints” (*59*) compare the potency of individual bNAbs against a large set of HIV-1 strains (**Fig. S22**). We observed clustering of RHA1.V2.01 within the general grouping of human V2 apex bNAbs, and more specifically, within the PGT145 class that includes PCT64 and PGDM1400 (**Figs. 6E,S19D,E**). We also performed a hierarchical clustering analysis of the neutralization profiles of the mAb RHA1.V2.01 and other prototypic human V2 apex bNAbs measured against the 208 virus panel (**Fig. S24A**). RHA1.V2.01 clustered most closely with PCT64-35M, a similarity that was strongly supported by the overlap of viruses that were sensitive or resistant to those antibodies [Fisher’s exact test p=2×10^-16^; odd’s ratio = 13.14; accuracy ratio (all concordant = 1, none = 0) = 0.78] (**Fig. S24B**) and by the correlation of IC_50_ titers of RHA1.V2.01 versus PCT64-35M (R^2^=0.4467; p=1.4×10^-10^) and other V2 apex bNAbs (**Fig. S24C**). We next examined neutralization profiles across different HIV-1 group M subtypes (**Fig. S25**). We found that RHA1.V2.01 was more subtype-independent than any of the human V2 apex bNAbs but was otherwise most similar to PCT64-35M and CAP256.VRC26. Finally, we performed a neutralization signature analysis using GenSig (https://www.hiv.lanl.gov/content/sequence/GENETICSIGNATURES/gs.html) (**Fig. S26**).

Neutralization “signatures” identify individual Env residues that contribute directly or indirectly to antibody binding, including those potentially involved in selecting for affinity maturation (*60*). The signature analysis was performed using Fisher’s test with a binary IC_50_ threshold greater or less than 50 µg/ml and a Wilcoxon test that measures the difference in IC_50_ distributions with and without a given amino acid residue. High-confidence signature sites were defined as those meeting at least two of three criteria: i) contact sites; ii) at least one phylogenetically corrected signature at the site; or iii) at least one signature at the site that had a false discovery rate q < 0.1. For the rhesus mAb RHA1.V2.01, statistically robust signatures were identified at residues 130, 160, 166, 167 and 169. Among all of the V2 apex bNAbs analyzed, only PCT64 shared all five of these signature sites.

#### Structure of V2-apex bnAb RHA1.V2.01

The structure of mAb RHA1.V2.01 in complex with the BG505 DS-SOSIP Env trimer, determined by cryo-EM at 3.9Å resolution, showed striking similarity to PGT145 and PCT64-35S (**Figs. 6F,G** and **Table S6**). These antibodies bind Env with a 1:1 stoichiometry near the trimer 3-fold axis and are surrounded by the three N160 glycans. Their respective CDRH3s adopt a needle-like antiparallel β-hairpin conformation that extends from the combining surface of the Fab and inserts into a cationic hole at the trimer apex. The C-terminal ends of each of the three C-strands abut the apex hole and are oriented perpendicular to the inserting HCDR3. Like PCT64 and PGT145, the acidic EDDY motif of RHA1.V2.01 was tyrosine-sulfated and made key contacts with Env residues 121, 166 and 169. (**Fig. 6F insert**). When the Env-bound structures of RHA1.V2.01, PCT64-35S and PGT145 were overlaid, the respective EDDY motifs aligned at the tips of their respective HCDR3 loops around the β-hairpin turn (**Fig. 6G**). Otherwise, the overall Fab orientations differed, indicating the HCDR3 tip structural mimicry to be the main source of the neutralization similarity among these antibodies. The HCDR1 of RHA1.V2.01, which contained a non-templated two amino acid insertion in addition to other strongly selected mutations, was sandwiched between the Env N160 glycans of two protomers and proximal to the C-strand of one, with buried surface area of 52 Å^2^ and 49 Å^2^ for the two glycans and a key electrostatic interaction between D29 and K171 (**Figs. 6F,S19A,S27F**). Thus, the V2 apex bNAb lineage in RM5695 exhibits genetic, chemical and structural solutions to epitope recognition that are shared with human V2 apex targeted bNAbs, especially PCT64 and PGT145.

## DISCUSSION

A principal finding of this study is that SHIVs bearing primary T/F HIV-1 Envs elicit strain-specific and heterologous NAbs in RMs that can replicate, to a striking degree, responses to HIV-1 in humans. This mimicry includes the frequency, kinetics, titers, immunogenetics, structures and target epitopes of elicited antibodies; structural and chemical features of epitope recognition; and coevolutionary pathways of antibody maturation and Env escape. All are key features to be considered in vaccine design. Our findings add substantially to earlier reports of sporadic or low-titer neutralization of heterologous tier 2 viruses elicited in RMs by SHIVs bearing lab-adapted or animal passaged HIV-1 Envs (*61–64*) or Env immunogens (*65–72*). Unlike previous reports, the current results show how closely neutralizing antibody responses in RMs can mirror responses in humans and indicate the extent to which protective responses elicited by reverse engineered or lineage-based vaccines in RMs might be expected to predict human responses to candidate vaccines (*10, 11, 73, 74*).

A surprising observation was the extent to which Env evolution in multiple rhesus animals recapitulated evolution of homologous Envs in human infections. Similarities included site-specific and amino acid-specific mutations and identities or near identities of insertions and deletions. These similarities likely resulted from: (i) the highly evolved and functionally constrained nature of primary T/F Env trimers; (ii) limited sites of antibody accessibility and variable fitness costs of escape mutations; and (iii) homologous germline B cell responses in different animals and humans to conserved Env epitopes. Equally surprising were the genetic and structural similarities between rhesus and human antibodies that targeted CD4bs or V2 apex epitopes and their conserved mechanisms of epitope recognition. This included the HCDR2-mediated CD4 mimicry of the rhesus antibody DH650 and the tyrosine-sulfated, 24 residue long HCDR3 of the rhesus antibody RHA1.V2.01, which bound N160 glycans and positively charged Env residues at positions 121, 166, 169 and 171. Together, the conserved patterns of Env-specific sequence variation and the homologous and orthologous B cell responses in humans and rhesus represent remarkable examples of convergent evolution (*14*) that may aid in the design and testing of novel HIV-1 vaccines.

Our findings suggest that HIV-1 Envs are not equal in their propensity for eliciting epitope specific bNAb responses. For example, we found that CAP256SU Env, which elicited V2 apex bNAbs in the human subject CAP256 (*3, 5, 25*), induced bNAbs of the same specificity in 2 of 6 SHIV infected RMs. CH848 Env, which elicited V3 glycan targeted bNAbs in a human subject (*7*), did the same in 2 of 6 SHIV infected RMs. And CH505 Env, which elicited CD4bs targeted bNAbs in a human subject (*2, 4, 6*), induced homologous strain-specific CD4bs targeted NAbs in RM6072. CH505 Env also elicited V2 apex targeted NAbs with variable breadth in vaccinated CH03 heavy chain Ig knock-in mice and an immunized RM (*69*). In the present study, it did so as well in 2 of 10 SHIV infected RMs. In other work (G.M.S., unpublished), we have identified fusion peptide targeted bNAbs in a SHIV.Ce1176 infected RM. These findings highlight the potential for RMs to develop antibody responses targeting different canonical bNAb epitope clusters and suggest a tendency for some Envs to elicit bNAb responses targeting particular epitopes.

SHIV replication in RMs is the only model system other than naturally infected humans where the immunogen (Env) coevolves with antibodies. The SHIV infected macaque can therefore be particularly informative for vaccine design by enabling the identification, and then rapid testing, of Env intermediates that guide the evolution of germline bNAb precursor B cells through stages of affinity maturation to acquire breadth and potency. The high mutability and dynamic replication of HIV-1 and SHIV result in a constantly evolving virus quasispecies, which means that Envs with binding affinities sufficient to drive bNAb lineage affinity maturation are constantly being generated and can be identified precisely. For example, in the CAP256 (*5*) and PCT64 (*8*) infected human subjects, viral sequences showed very similar patterns of Env evolution at residues 166 and 169, which in turn were similar to the pattern in the SHIV.CH505 infected RM5695. Deep sequencing of RM5695 plasma vRNA covering the V1V2 region revealed selection focused primarily on residues 166 and 169 with mutations at these two sites rapidly and completely replacing the infecting virus strain. The infecting SHIV.CH505 virus had an Arg at these two positions, which evolved progressively to R166K or R169K and then to R166G, R166S or R166T (**Fig. 3**). The earliest mutants, R166K or R169K, were approximately 5-fold more resistant to the mature rhesus bNAb mAbs than the infecting virus, whereas the subsequent R166G, R166S or R166T mutants were >100-fold more resistant (**Fig. 6D**). Thus, sequential Envs that varied at residues 166 and 169 in animal RM5695 showed progressive phenotypic escape from V2 apex bNAb antibodies, closely resembling the viral Env-bNAb coevolution observed in humans CAP256 and PCT64. Cryo-EM analysis of the RHA1.V2.01 mAb provided a structural explanation for this loss of antibody recognition by showing that Env residues 166 and 169 were primary electrostatic contacts with the antibody. Mutations in these two residues in the V2 apex appear to be largely or solely responsible for driving affinity maturation of diverse antibody lineages to breadth in multiple rhesus animals and humans within a relatively short time (<1 yr), suggesting that Env intermediates or “immunotypes” (*5*) required for V2 apex bNAb elicitation may be few and simple. This hypothesis has important implications for V2 apex targeted HIV-1 vaccine design, which can be tested rapidly in Ig knock-in mice and outbred macaques using immunogens designed from V2 apex variants of CH505 and other primary Envs. The goal of such research would be to learn the “rules” governing consistent bNAb induction in RMs and then to translate these findings to human studies using SOSIP, mRNA or other non-SHIV based vaccine platforms.

V3 high mannose patch peptidoglycans are also commonly targeted by bNAbs in HIV-1 infected humans (*75*) and are of high interest for HIV-1 vaccine development (*50, 76, 77*). Site-directed mutagenesis coupled with antibody neutralization showed that polyclonal bNAb responses in SHIV.CH848 infected RMs 6167 and 6163 targeted canonical N332 and ^324^GDIR^327^ motifs, similar to human V3 glycan bNAbs (*78, 79*). Deletions in V1 of SHIV.CH848 sequences preceded the development of V3 glycan bNAbs in both monkeys and in the human subject CH848. Long, glycosylated (N133, N138) V1 segments obstruct access of V3 glycan bNAbs (*7, 80*), and germline-targeted Env immunogens with shortened V1 segments depleted of glycans enhance Ab access to V3 high mannose patch epitopes (*50, 76*). Because Env-Ab coevolution leading to V3 glycan bNAbs generally requires more extensive somatic hypermutation compared with V2 apex bNAbs (*10, 11*), a rhesus model in which vaccinations with germline-targeted Envs (*50, 76, 77*) is followed by infection with SHIVs whose Envs are similarly targeted, offers a novel strategy for identifying the “finishing” or “polishing” immunogens necessary for bNAb affinity maturation.

It is generally believed that the development of an effective neutralizing antibody-based HIV-1 vaccine will require consistent activation of multiple germline precursor B cells that express immunoglobulin receptors specific for one or more of the canonical bNAb epitope clusters, followed by efficient antigen-driven selection for antibody affinity maturation (*10, 11, 50, 73, 74, 76, 77, 81*). The present study demonstrates that the SHIV-infected rhesus model can inform both of these critical steps in bNAb elicitation. The fact that only a minority of SHIV infected animals in the current study developed bNAbs is a faithful reflection of the natural prevalence of bNAb responses in HIV-1 infected humans (*75, 82*) and further argues for the relevance of the rhesus model. It should be possible to combine established immunization platforms such as Env trimers, outer domain scaffolds, virus-like particles, or DNA/RNA expression with SHIV infections to identify optimized priming and boosting immunogens that elicit broad neutralization in macaques as a guide for HIV-1 vaccine design in humans.

## SUPPLEMENT

### Extended Discussion

#### Convergence in the Indel mutations in the hypervariable V1, V4 and V5 loops of CH505 and CH848 infected humans and rhesus macaques

In functional sequences, insertions and deletions (Indels) occur in multiples of three nucleotides so as to retain the integrity of the Env open reading frame. Insertions in the hypervariable regions primarily occur by duplications, which result in perfect or imperfect direct repeats (*47*). In the CH505 infected human and rhesus animals, there was a striking recurrence of precise Indel events (**Fig. S9**). In V1, for example, *env* sequences from the human and all six animals contained an identical 3 nucleotide deletion. The human and three RMs contained an identical 12 nucleotide perfect direct repeat insertion. Five additional distinct insertions of between 6 and 24 nucleotides were shared among between the human sequences and a subset of rhesus macaque sequences. The most striking set of shared Indel patterns overall was observed in subject CH505 and RM6069, where the two shared five unique insertions and one deletion. The other five animals shared between two and three Indels identically with the human subject (**Fig. S9**). To estimate the probability that these shared Indels could have occurred by chance, we subjected sequences from the human subject CH505 and from three animals (RMs 6069, 6070, 6072) to rigorous statistical analysis (see below for statistical methods). First, we considered the natural distribution of V1 loop lengths found in the LANL database (www.hiv.lanl.gov) and used a comparable window in time from the longitudinal sampling (∼1 year from infection) to study the evolution of the hypervariable loops in subject CH505 and the RMs. We estimated the probability of 5 precisely shared Indels in both the human CH505 and RM6069 at p<3.3×10^-9^ by chance alone, and the probability of all three monkeys sharing as many precise Indels with CH505 as p<3.0×10^-13^. Another approach to estimating the likelihood of seeing identical Indels is to focus on the 3 base deletion in V1 shaded in red (**Fig. S9**) that was found in the human CH505 and in all 6 RMs infected with SHIV CH505. The probability of an identical deletion occurring in all 7 hosts by chance was estimated to be p<6.2 x 10^-7^. Seven distinct insertions of 3 to 9 nucleotides were identified in the hypervariable region of V5 in CH505 infected human or macaque sequences. Each of these insertions represented perfect direct repeats. One of these Indels was shared between the human and RM6069, and five others were shared among different animals. Two direct repeats were shared among three animals. The likelihood of this happening by chance was estimated to be p < 1.5 x 10^-5^. Additional Indels in the V4 region of CH505 Envs were shared between human and rhesus (**Fig. S9**), and still other indels in V1, V4 and V5 were shared between the CH848 infected human and rhesus animals (**Fig. S10**). The statistical likelihood that these reiterations occurred by chance is vanishingly small. Instead, the repetition of Indels in human and rhesus variable loop sequences attests to the fitness advantage associated with specific shared patterns of convergent or parallel evolution.

##### Statistical methods to analyze repeated Indels

We first collected all viable V1 loops, Cys to Cys, from the LANL database (www.hiv.lanl.gov) of 2546 full HIV-1 genome DNA alignments to get a baseline assessment of what is possible and common for the virus in terms of V1 loop lengths. We assumed that this distribution reflects how readily HIV-1 tolerates variation in V1 length and how likely V1 is to take on a particular size. We used this distribution as an *a priori* baseline rather than assuming all sizes are equally probable. We also assumed that the strand switching that gives rise to the indels is not impacted so much by length of the indel but rather by what sequences are observed and by how well the length of the V1 is tolerated after the primary indel-mutation event. CH505 T/F Env has a relatively short V1 length of 69 nucleotides. One of the V1 insertions was a perfect direct repeat of 15 nucleotides (AATGCTACTGCCAGCAATGCTACTGCCAGC), which gave rise to a V1 length of 69+15=84 nucleotides. In the HIV-1 database, the frequency of 84 base V1 segments is 195/2546=0.0766, so we made the assumption that random Indels would create a length of 84 bases 0.077% of the time. The hypervariable part of V1 is 51 bases long (this is the stretch within V1 that is unalignable in the database) and we assume an Indel that retains the correct reading frame can happen anywhere in this 51 base

long stretch, with the probability of this particular length insertion at the exact location estimated at 0.0766/51=0.00150. We then derive:

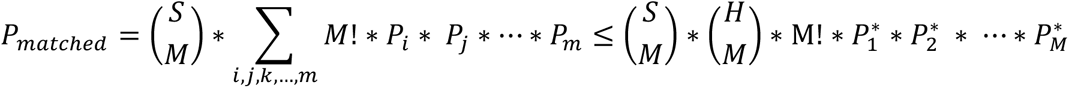

where

*H* = number of human Indels

*M* = number of matched Indels found in monkeys and human

*S* = number of monkey Indels

*P_i_* = probability of seeing Indel *i (length*location)*

Let *P_1_^*^,P_2_^*^,…,P_M_^*^* be the M largest values of *P_i_*

There are many different ways (*H* choose *M*) of matching *M* out of *H* Indels, but the probability is different for each choice of *M*, and depends on the probabilities *P_i_, P_j_, P_k_, P_l_, P_m_* associated with the particular *i,j,k,l,m* in each choice. Since order doesn’t matter, there is an extra factor of *M!* in the probability. The separate computation for each combination of M out of H would give the probability for that particular combination; we add those probabilities to get an overall probability of matching M out of H. To simplify this, we can place an upper-bound on the probability that a set of M Indels would exactly be matched in a monkey by recognizing that there are (H choose M) terms, and that all the terms are less than or equal to the maximum term. The S choose M prefactor takes into account that there were more Indels in the monkey than matched the human. This maximum is computed by taking the product of the M largest probabilities; in this study the upperbound was extremely small so we didn’t take this further.

The probability of the number of 5 shared Indels between CH505 and RM6069 happening by chance alone is <3.3×10^-9^

10 choose 5 = 252, 5! = 120

10 chose 1 = 10

10 choose 2 = 45, 2! = 1

7 choose 5 = 21

P_RM6069_ = 21*252*120*(0.0015*0.0015*0.0015*0.0013*0.0012) = 3.3×10^-9^

P_RM6070_ = 3*10*0.0015 = 0.045

P_RM6072_ = 1*45*2*(0.0015*0.0015) = 0.0002

The probability of all three monkeys having the shared Indels that were observed occurring by chance alone is <3.0×10^-13^.

We can also ask how likely it would be for all four hosts (one human and three monkeys) to share the same Indel; we ask this without specifying which Indel is shared. The question is: what is the chance that any Indel is shared among all four? Let P_1_, P_2_, …, P_K_ correspond to the K possible Indels. Let A1, A2, A3, A4 be the number of Indels observed in the four animals (for in our study, A1=10, A2=7, A3=3, A4=2); then A_i_P_k_ is the probability of observing the k’th Indel in the i’th animal. And A1A2A3A4Pk^4^ is the probability of observing a specific Indel k in all animals. Thus we can write

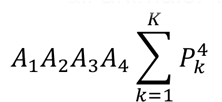

as the probability of observing an unspecified Indel in all four animals. This probability is: 10*7*3*2*(1.47975 x 10^-9^) = 6.2 x 10^-7^, as 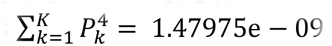.

The length of the hypervariable region of V5 in CH505 T/F is 30 bases, which is short compared to global group M sequences with a median of 39. Selective pressures in the human subject CH505 and in 4 of 6 monkeys drove V5 to get longer throughout the course of infection. Given that identical insertions showed up in multiple individuals, we asked how likely it would be for three individuals to share the same Indel. Again, we asked this without specifying which Indel is shared. Let P_1_, P_2_, …, P_K_ correspond to the K possible Indels. Let A1, A2, A3 be the number of Indels observed in the 3 animals (for in our study, A1=5, A2=4, A3=5,); then A_i_P_k_ is the probability of observing the k’th Indel in the i’th animal. And A1A2A3Pk^3^ is the probability of observing a specific Indel k in all animals. Thus we can write

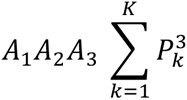

as the probability of observing an identical Indel in all three animals. This probability is 5*4*5*(3.89133e-05)= 0.0039 for a single event. The likelihood that two unique Indels would occur in three animals, as was the case for RMs 6072, 6069 and 5695, is 0.00389^2 = 1.5 x 10^-5^.

#### Enhanced N160K responses

In **Figs. 1**, **3** and **S12**, we showed evidence of V2 apex, C-strand targeted bNAb activity in the plasma of RMs 5695, 6070, 42056 and 40591. These data included broad neutralizing activity against multiple heterologous HIV-1 strains; loss of neutralizing activity when key residues 166 and 169 were mutated to eliminate positively charged arginines or lysines; and detection of bNAb escape mutations at these same 166 and 169 residues in vRNA sequences from serially collected plasma specimens. In animal RM5695, we isolated four bNAb mAbs (RHA1.V2.01-04) that exhibited these properties and accounted for most of the neutralization breadth observed in the animal’s plasma. We then showed by Cryo-EM that the mAb RHA1.V2.01 binds to the trimer apex-hole residues with contacts to C-strand residues and N160 glycan of the BG505 DS-SOSIP trimer (**Figs. 6, S19,27**). A key feature of the RHA1.V2.01 mAb was its strict dependence on N160 for binding and neutralization. This is an expected property since nearly all reported human V2 apex targeted bNAbs (e.g., PG9, PGT145, CH01, PCT64) depend on interactions with N160 for their binding and neutralization activity. The exceptions are some members of the VRC26 lineage of V2 apex bNAbs that do not require N160 for their activity (*3, 49*). A surprising finding was that the plasma from RMs 5695, 6070, 42056 and 40591 variably neutralized heterologous viruses strains with N160K substitutions, depending on the Env background (**Figs. 3**, **S13,S23**). This included animal RM5695 whose RHA1 broadly neutralizing mAbs were strictly N160 dependent. Plasma from RM5695 showed enhanced neutralization of Q23.17.N160K, no difference in neutralization of T250 wt compared with T250.N160K, and failed to neutralize BG505.N160K (**Fig. S13A**). Plasma from animals RM6070 (**Fig. S13B**) and RM42056 (**Fig. S13D**) exhibited plasma neutralization titers against heterologous viruses with N160K substitutions that were extraordinarily high (ID_50_ >1:100,000). This potent neutralization was directed entirely to the V2 apex, since N160K.K169E double mutants eliminated or drastically reduced the N160K enhanced neutralization (**Figs S13B,D**). We considered two possible explanations for these findings: (i) In RMs 5695, 6070 and 42056, strain-specific 2909-like antibodies (*83, 84*) were elicited in addition to prototypical N160 dependent bNAbs; and (ii) C-strand targeted V2 apex bNAbs were elicited that do not require binding to N160 for their activity but instead are enhanced in potency when the protective shielding afforded by N160 is lost. Recently, we isolated rhesus V2 apex C-strand targeted broadly neutralizing mAbs from a SHIV.CH505 infected RM that was not part of the current study that exhibits the latter features (R. Roark and G. Shaw, unpublished).

### Materials and Methods

#### Nonhuman Primates

Indian RMs were housed at Bioqual, Inc., Rockville, MD, according to guidelines of the Association for Assessment and Accreditation of Laboratory Animal Care standards. Experiments were approved by the University of Pennsylvania and Bioqual Institutional Animal Care and Use Committees. RMs were sedated for blood draws or excisional biopsies of lymph nodes, anti-CD8 mAb infusions, and SHIV inoculations. Selected animals received an intravenous infusion of either 25-50 mg/kg of anti-CD8alpha mAb (MT807R1) or anti-CD8beta mAb (CD8beta255R1) one week prior to or at the time of SHIV inoculation. SHIV inoculations were intravenous and consisted of an equal mixture of six Env375 allelic variants, 50 ng p27Ag of 293T-derived virus of each, in 1 ml of DMEM culture medium plus 10% heat-inactivated fetal bovine serum (FBS). Blood was collected in sterile 10 ml yellow-top vacutainers containing acid citrate dextrose A as an anticoagulant and sent same-day or overnight to the University of Pennsylvania for processing.

#### Processing and storage of clinical specimens

40 ml of ACD-A anticoagulated blood was combined in a sterile 50 mL polypropylene conical tube, centrifuged at 2100 rpm (1000xg) for 10 min at 20°C, and the plasma collected in a fresh 50 mL conical tube without disturbing the buffy coat WBC layer and large red cell pellet. The plasma was centrifuged again at 2500 rpm (∼1500g) for 15 minutes at 20°C in order to remove all platelets and cells. Plasma was collected and aliquoted into 1 ml cryovials and stored at −80°C. The RBC/WBC pellet was resuspended in an equal volume of Hanks balanced salt solution (HBSS) without Ca^++^ or Mg^++^ and containing 2mM EDTA and then divided into four 50 ml conical tubes. Additional HBSS-EDTA (2mM) buffer was added to bring the volume of the RBC/WBC mixture to 30 ml in each tube. The cell suspension was then carefully underlayered with 14 ml 96% Ficoll-Paque and centrifuged at 1800 rpm (725xg) for 20 min at 20°C in a swinging bucket tabletop centrifuge with slow acceleration and braking so as not to disrupt the ficoll-cell interface. Mononuclear cells at the ficoll interface were collected and transferred to a new 50ml centrifuge tube containing HBSS-EDTA (w/o Ca^++^ or Mg^++^) and centrifuged at 1000 rpm (∼200 g) for 15 min at 20°C. This pellets PBMCs and leaves most of the platelets in the supernatant. The supernatant was removed and the cell pellet was resuspended in 40 ml HBSS (with Mg^++^/Ca^++^ and without EDTA) + 1% FBS. To remove additional contaminating platelets, the cell suspension was centrifuged again at 1000 rpm (∼200 g) for 15 minutes at 20°C and the supernatant discarded. The cell pellet was tap-resuspended in the residual 0.1-0.3 ml of media and then brought to a volume of 10 ml HBSS (with Mg^++^/Ca^++^) +1%FBS. Cells were counted and viability assessed by trypan blue exclusion. Cells were centrifuged again at 1200rpm (300xg) for 10 min at 20°C, the supernatant discarded, and the cells resuspended at a concentration of 5-10×10^6^ cells/ml in CryoStor cell cryopreservation media (Sigma Cat. C2999) and aliquoted into 1ml cryovials (CryoClear cryovials; Globe Scientific Inc., Cat. 3010). Cells were stored in a Mr. Frosty at −80°C overnight and then transferred to vapor phase liquid N_2_ for long-term storage. Mononuclear cells collected from lymph nodes (LN) and spleen were processed similar to blood mononuclear cells. LN nodes and spleen were excised and placed immediately into RPMI1640 medium on wet ice. LNs were diced with a sterile scalpel and spleen was homogenized and the material passed through a sterile mesh grid. Cells were collected from the pass-through and subjected to Ficoll density gradient purification as described above.

#### SHIV construction and characterization

The experimental design for constructing SHIVs bearing primary or transmitted/founder Envs with allelic variation at gp120 residue 375 was previously described, including SHIV.CH505 and SHIV.CH848 (*9*). For the construction of SHIV.CAP256SU, we synthesized (GenScript) sequence KF996583.1 from GenBank (GenBank: KF996583.1) and cloned it into the pCRXL-TOPO-SIVmac766 backbone (*9*) by recombinant PCR. The QuikChange II XL Site-Directed Mutagenesis kit (Agilent Technologies) was used to create allelic variants (M, Y, F, W, or H) of the wild type Env375S codon. Wild type and mutant plasmids were transformed into MAX Efficiency Stbl2 Competent Cells (Invitrogen) for maxi-DNA preparations. Each 10-kb viral genome was sequenced in its entirety to authenticate its identity. Infectious SHIV stocks were generated in 293T cells as previously described (*9*). On day 0, five million 293T cells were plated in 100-mm tissue culture dishes in 10 mL of complete MEM growth media with 10% FBS. On day 1, 6 ug of SHIV plasmid DNA was combined with 18 uL of FuGENE 6 (Promega) in 500 µL of DMEM was added dropwise to tissue culture dishes. Media containing virus was harvested on day 3 and aliquoted for storage at −80C. Virus concentration was estimated by p27 antigen (p27Ag) ELISA (Zeptometrix) and infectious particle concentration was determined by entry into TZM-bl cells in the presence of DEAE–Dextran, as previously described (*21*). Typically, 293T-derived SHIV stocks contained >1,000 ng/ml p27Ag and >1,000,000 IU/ml on TZM-bl cells. The replication kinetics of each of the SHIV.CAP256SU Env375 variants in primary, activated human and rhesus CD4 T cells were determined as previously described (*9*). 293T supernatants containing 300 ng p27Ag of each variant, were added to 2 x 10^6^ purified human or rhesus CD4 T cells in complete RPMI growth medium (RPMI1640 with 15% FBS (Hyclone), 100 U/mL penicillin–streptomycin (Gibco), 30 U/mL IL-2 (aldesleukin, Prometheus Laboratories) and 30 µg/ml DEAE-Dextran. 300 ng p27Ag is equal to ∼3 x 10^9^ virions, ∼3 x 10^5^ IU on TZM cells, or ∼3 x 10^4^ IU on primary CD4 T-cells, so the estimated MOI of this titration was estimated to be ≤ 0.1. The cell and virus mixtures were incubated for 2 hours under constant rotation at 37C to facilitate infection, washed three times with RPMI1640, and resuspended in complete RPMI1640 medium lacking DEAE-Dextran. Cells were plated into 24-well plates at 2 million cells in 1 ml and cultured for 11 days, with sampling of 0.2ml supernatant and media replacement every 2-3 days for 11 days. Supernatants were assayed for p27Ag concentration by ELISA (Zeptometrix).

#### Plasma vRNA quantification

Plasma viral load measurements were performed by the NIH/NIAID-sponsored Nonhuman Primate Virology Core Laboratory at the Duke Human Vaccine Institute. This core facility is CLIA certified and operates a highly standardized, quality-controlled Applied Biosystems Real-Time SIV and HIV vRNA PCR assays. QIAsymphony SP and QIAgility automated platforms (QIAGEN) are used for high throughput sample processing and PCR setup. Viral RNA is extracted and purified from plasma, annealed to a target specific primer and reverse transcribed into cDNA. The cDNA is treated with RNase and added to a custom real-time PCR master mix containing target specific primers and a fluorescently labeled hydrolysis probe. Thermal cycling is performed on a QuantStudio3 (ThermoFisher Scientific) real-time quantitative PCR (qPCR) instrument. Viral RNA cp/reaction is interpolated using quantification cycle data. Raw data is quality-controlled, positive and negative controls are checked, and the mean viral RNA cp/mL is calculated. The current limit of quantification using 0.5 ml of NHP plasma is set at 62 RNA cp/mL with a 94% detection and a Clopper-Pearson binomial confidence interval of 84.8 and 98.3. However, this limit of quantification was previously as high as 250 RNA cp/ml, so for this study we adopted this more conservative limit.

#### Viral sequencing, pixel plots and LASSIE analysis

Single genome sequencing of SHIV 3’ half genomes was performed as previously described (*1, 9*). Geneious R7 was used for alignments and sequence analysis and sequences were visualized using the LANL Highlighter and Pixel tools https://www.hiv.lanl.gov/content/sequence/pixel/pixel.html https://www.hiv.lanl.gov/content/sequence/HIV/HIVTools.html (*44*). The specific implementation of this software for this project is described in the figure legends.

#### IgG isolation from plasma

Total polyclonal IgG was isolated from rhesus plasma using the Protein A/Protein G GraviTrap kit (GE Healthcare). Plasma was heat-inactivated (1 hour at 57C), clarified by centrifugation at 21,000g for 4 min, and applied to the Protein A/G column. The sample was washed and eluted per the manufacturer’s instructions, and then buffer-exchanged with phosphate buffered saline (PBS). The concentration of purified IgG sample was quantified using the Pierce BCA Protein Assay Kit (ThermoScientific).

#### Neutralizing antibody assay

Assays for neutralizing antibodies were performed using a single-round TZM-bl indicator cell line, as previously described (*9, 21*). This assay is essentially identical to that employed by Montefiori, Seaman and colleagues (*85*) https://www.hiv.lanl.gov/content/nab-reference-strains/html/home.htm, the differences being that in our assay we plate virus and test plasma onto adherent TZM-bl cells and we hold the concentration of test plasma constant (generally 5% vol/vol) constant across all wells. This is in addition to 10% heat-inactivated fetal bovine serum in the complete RPMI1640 culture medium. Target viruses express HIV-1 Envs reported elsewhere (*29, 30, 86*).

#### Binding antibody assays

HIV-1 Env binding by recombinant mAbs, and plasma or sera, were tested in ELISA as previously described (*50, 87*). In brief, recombinant Envs were coated directly to Nunc-absorb (ThermoFisher) plates overnight at 4°C or captured using AbC-mAb (AVIDITY, Colorado, USA) that was directly coated to Nunc-absorb plates overnight at 4°C. Antibody binding was detected with goat anti-human or goat anti-rhesus HRP-labeled anti-IgG Fc antibodies (Jackson ImmunoResearch Laboratories), and HRP detection was subsequently quantified with 3,3′,5,5′-tetramethylbenzidine (TMB). Competitive ELISA to assess cross-blocking of recombinant mAbs or plasma antibodies were previously described (*7, 88*). We biotinylated the antibodies using the following product: BIOTIN-X-NHS, Cayman Chemicals, CAT# 13316. Competitive inhibition of biotiniylated-mAbs was measured as a percent of binding in the presence of a competing non-biotinylated mAb relative to binding in the absence of this competing mAb. MAbs were also tested for binding HIV-1 Envs using Biolayer interferometry (BLI) as described (*89*). Here, antibody binding was measured using mAb-captured sensors that were placed into solutions of CH505 gp120 or SOSIP trimers at 50 µg/ml for 1000s. MAbs were captured using anti-human IgG Fc sensors, and non-specific or background binding was subtracted using binding levels by anti-influenza HA mAb (CH65).

#### Rhesus B cell staining and sorting of strain-specific mAbs

CH505 gp120 T/F CD4bs-specific antibodies were isolated from memory B cells in PBMCs, lymph node or bone marrow collected at weeks 20, 24, 32 and 52 using two approaches: direct single-cell sorting into PCR plates (weeks 20, 32 and 52) (*26, 90*), and memory B cell cultures (week 24) (*91, 92*). For direct sorting, we performed single-cell isolation of memory B cells decorated with AlexaFluor® 647 (AF647) or Brilliant Violet 421 (BV421)-tagged HIV-1 CH505 TF gp120 using a BD FACSAria™ or a BD FACSAria™ II (BD Biosciences, San Jose, CA), as previously described (*26*). The flow cytometry data were analyzed using FlowJo (Treestar, Ashland, OR) (*90, 93, 94*). For our sort strategy, we isolated antigen-specific IgD-negative, CD27-All memory B cells that bound BV421-tagged CH505 T/F gp120, but not AF647-tagged CH505 T/F gp120 Δ371 mutant protein; antibodies isolated from these B cells were referred to as CH505 differential binders or CD4BS antibodies (*26*). For memory B cell cultures, from 8 million PBMCs, we sorted 30,792 CH505 T/F gp120-specific B cells, defined as CD3-negative, CD14-negative, CD16-negative, IgD-negative, CD27-All, CD20-positive, AF647-tagged CH505 T/F gp120-positive and BV421-tagged CH505 T/F gp120-positive. As previously described (*92*), cells were flow sorted in bulk into wells containing 5,000 MS40L feeder cells, RPMI-1640 supplemented with 15% FBS, 1 mM sodium pyruvate, 1% non-essential amino acids, 25 mM HEPES buffer, 2.5 μg/ML ODN2006 (Invivogen, TLRL-2006-5), 5 μM CHK2-inhibitor (Calbiochem, 220486), 100 ng/mL recombinant human interleukin (IL)-21 (Peprotech, Cat. no. 2001-21), 10 ng/mL recombinant Human BAFF (Peprotech, Cat. no. 310-13), 200 U/ml IL-2 (from the myeloma IL-2 producing cell line IL2-t6, kindly provided by Dr. Antonio Lanzavecchia, IRB, Bellinzona, Switzerland), and 100 μL supernatant of the Herpesvirus papio (HVP)-infected Baboon cell line S594 (NHP Reagent Resource). The concentration of each supplement was previously determined to achieve optimal *in vitro* stimulation. Following overnight incubation at 37 °C in 5% CO_2_, memory B cells were transferred at limiting dilution into 96-well round bottom tissue culture plates containing 5,000 MS40L feeder cells. Culture medium was refreshed 7 days after plating and harvested 2 weeks after plating to test for binding to CH505 T/F gp120, CH505 T/F gp120 Δ371, RSC3 (*95*) and RSC3 Δ371 P363N, as well as for neutralization of pseudotyped the CH505 T/F HIV-1 strain in the TZM-bl-based neutralization assay using a single dilution of supernatant (*91, 96*). CD4bs DH650 lineage autologous tier 2 NAbs were isolated from CH505 differential-binding memory B cells in PBMCs from week 20, 24 and 32 post SHIV CH505 infection of RM6072. In capturing maximum numbers of B cells bearing candidate CD4bs antibodies, we used a less stringent gating strategy for capturing CH505 T/F gp120 (+) and CH505 T/F gp120 Δ371 (-) B cells. In so doing, we also captured nonCD4-binding site autologous tier 2 NAbs, including DH647 and DH648 that were isolated from memory B cells in PBMCs at week 20 post SHIV infection of RM6072.

#### Rhesus B cell staining, culture and microneutralization screening for V2 apex bNAbs

Cryopreserved PBMCs from RM5695 at week 65 post-SHIV infection were thawed and stained with LIVE/DEAD Fixable Aqua Dead Cell Stain (Life Technologies), as previously described (*97, 98*). Cells were washed and stained with an antibody cocktail of CD3 (clone SP34-2, BD Biosciences), CD4 (clone OKT4, BioLegend), CD8 (clone RPA-T8, BioLegend), CD14 (clone M5E2, BioLegend), CD20 (clone 2H7, BioLegend), IgG (clone G18-145, BD Biosciences), IgD (polyclonal, Dako) and IgM (clone G20-127, BD Biosciences) at room temperature in the dark for 20 mins. The stained cells were washed 3 times with PBS, re-suspended in 1 ml of PBS and passed through a 70 μm cell mesh (BD Biosciences). Total memory B cells (CD3-CD4-CD8-CD14-CD20+IgD-IgM-IgG+) were sorted with a modified 3-laser FACSAria cell sorter using the FACSDiva software (BD Biosciences) and flow cytometric data was subsequently analyzed using FlowJo (v9.7.5). B cells were sorted at 1 cell per well of a 384-well plate containing B cell culture media based on a human B cell culture protocol (*99*) that was optimized for the expansion of rhesus B cells. Briefly, sorted B cells were expanded for 14 days in B cell culture medium consisting of Iscove’s modified Dulbecco’s medium (IMDM) with GlutaMAX™ supplemented with 10% heat-inactivated fetal bovine serum (FBS), 1X MycoZap Plus-PR, 100 U/ml IL-2, 0.05 ug/ml IL-4, 0.05 ug/ml IL-21, 0.05 ug/ml BAFF, 2 ug/ml CpG ODN2006, and 3T3-msCD40L feeder cells at a density of 5,000 cells per well. Supernatants from ∼14,000 individual wells were evaluated for neutralization of HIV-1 Q23.17 and T250-4 Env-pseudotyped viruses using the high throughput NVITAL automated microneutralization assay, as previously described (*100*). Wells were selected for RT-PCR based on 50% reduction in infectivity of at least one virus. Out of nearly 14,000 wells tested, five met these neutralization criteria. Paired heavy and light Ig chains were successfully amplified from four of these wells. These amplicons were sequenced, cloned and expressed as IgG1 (RHA1.V2.01-04), and all four mAbs exhibited broad and potent neutralization.

#### Rhesus B cell cloning and expression

Heavy (IGHV) and light (IGKV, IGLV) chain genes were isolated via single cell PCR approaches (*90, 101*), and the gene sequences were computationally analyzed using rhesus Cloanalyst program (*102–104*). Antibody immunogenetics were reported as gene families and segments, mutation frequencies, and CDR3 lengths using the rhesus Cloanalyst database of reference genes (*24*). Using the rhesus cloanalyst program, we identified B cell clonal lineages for antibodies with the same inferred IGHV VDJ rearrangement and CDR3 length, and paired with the same light chain (Ig VJ segments). For DH650 lineage, the unmutated common ancestor (UCA) and intermediate (IA) genes were inferred computationally using the Cloanalyst program. The automated inference of antibody clonality was followed up by visual inspection of the DNA sequence alignments for confirmation. The heavy and light chain gene sequences from the sorted B cells or inferred UCA and IAs were commercially generated, and used to express purified recombinant mAbs as described (*90*). Heavy and light chain immunoglobulin repertoire next generation sequencing of monkey RM6072 was performed with the Illumina MiSeq platform utilizing primers targeting the VH1 and V*k*2 families to identify DH650 clonal members using a previously described protocol (*26, 105*). For mAb isolation from RM5695, bulk cDNA was synthesized from the five neutralization-positive B cell culture wells using random hexamers as previously described (*106*). Subsequently, immunoglobulin heavy chain (IgG) and light chain (IgK and IgL) variable regions were separately amplified by nested PCR cycles using pools of rhesus macaque primers as previously described (*107*). Sequences were analyzed using Cloanalyst (Kepler et al, Front Imunol 2014) to infer putative variable region germline genes and identify clonal lineages. The heavy and lambda chain variable regions of the RM5695 lineage were codon optimized, synthesized with a murine immunoglobulin signal peptide (tripeptide sequence VHS) immediately preceding the 5’ end of FRW1, and cloned into rhesus IgG1 (RhCMV-H) and IgL (RhCMV-L) expression vectors directly upstream of the respective constant regions using AgeI/NheI and AgeI/ScaI restriction sites, respectively (*107*). Recombinant mAbs were expressed by cotransfection of paired heavy and lambda chain plasmids into 293Freestyle cells as previously described (*95*), purified from cell supernatant using the Protein A/Protein G GraviTrap kit (GE Healthcare), and buffer-exchanged into PBS.

#### Next generation sequencing of naïve B cells and IgDiscover analysis

PBMCs isolated from RM5695 plasma obtained at week 48 post-infection were stained with LIVE/DEAD Aqua, CD3-PerCP-Cy55, CD4-BV785, CD8-BV711, CD14-PE-Cy7, CD20-BV605, IgD-FITC, IgG-Ax680, and IgM-BV650. Approximately 300,000 naive B cells (CD20+, IgG-, IgD+, IgM+) were bulk sorted into RPMI with 10% FBS and 1% Pen-Strep using a BD FACSAria II. Total RNA was extracted using RNAzol RT per the manufacturer’s guidelines (Molecular Research Center, Inc). Reverse transcription of mRNA transcripts, IgM and IgL variable region library preparation, and next generation sequencing were performed as previously described (*108*), as these methods can be efficiently used for both humans and rhesus macaques. Both heavy and lambda immunogloblulin libraries were sequenced on Illumina Miseq with 2×300 bp runs using the MiSeq Reagent V3 kit (600-cycle). Filtered, quality-controled IgM and IgL sequences were analyzed using IgDiscover (*57*) to curate a personalized immunogloblin repertoire library for RM5695. A naïve IgK variable region library was not prepared since the RM5695 lineage utilizes a lambda chain. IgDiscover can identify functional VH, VL, VK, JH, JL and JK genes, and denotes any novel genes and/or alleles with respect to the provided reference database. For this analysis, a recently published rhesus macaque database was used as the template (*24*). The resulting personalized database was then used in SONAR (*109*) to accurately assign germline immunoglobulin genes and precisely determine somatic hypermutation frequencies for the RHA1.V2 lineage.

#### Auto/Polyreactivity analysis

RHA1.V2.01 mAb reactivity to nine autoantigens was measured using the AtheNA Multi-Lyte ANA kit (Zeus scientific, Inc, #A21101). Antibodies were 2-fold serially diluted starting at 50 µg/mL and kit methods were followed per manufacturer’s instructions. Samples were analyzed using AtheNA software. Indirect immunofluorescence binding of RHA1.V2.01 mAb to human epithelial (HEp-2) cells (Zeus Scientific, Somerville, NJ) was also performed per manufacturer’s instructions. Briefly, 20µL of diluted antibody (50µg/ml) was added to antinuclear antibody (ANA) test slides. Slides were incubated for 20 minutes at room temperature in a humid chamber, and then washed with 1X PBS. Goat-anti-rhesus Ig-FITC (Southern Biotech, Birmingam, AL) secondary antibody was added at a concentration of 30µg/ml to each well. The slides were incubated for 20 minutes, washed twice, dried, fixed with 33% glycerol and cover-slipped. Slides were imaged using an Olympus AX70 microscope with a SpotFlex FX1520 camera. Images were acquired on a 40X objective using the FITC fluorescence channel. Positivity was determined by comparison with positive and negative control non-human primate mAbs DH1037 and DH570.30, respectively. Staining patterns were identified using the Zeus Scientific pattern guide.

#### Hierarchical clustering of RHA1.V2.01 and other V2 apex bNAb profiles

We used the Heatmap webtool at the Los Alamos HIV database https://www.hiv.lanl.gov/content/sequence/HEATMAP/heatmap.html (*95, 108, 109*). Input data were Log10 transformed IC_50_ titers for RHA1.V2.01 and other V2 apex bNAbs for the 208 virus panel. For clustering, Euclidean distances and the Ward algorithm (*110*) were used. Bootstraps were calculated using 1000 iterations.

#### Neutralization Signature Analyses

These analyses were performed using the webtool GenSig (https://www.hiv.lanl.gov/content/sequence/GENETICSIGNATURES/gs.html) (*59*) as described (*60*). Briefly, IC_50_ titers for RM5695 RHA1.V2.01 and other V2 apex bNAbs, together with Env sequences from the 208 strain virus panel, were used as inputs. Two statistical strategies were used: i) Fisher’s test with a binary phenotype, IC_50_ titer above or below the threshold (resistant and sensitive, respectively), and ii) Wilcoxon test that compares distribution of IC_50_ titers with and without a feature. The latter analyses were conducted two ways, including and excluding resistant viruses. In both strategies, every amino acid and glycan at each site were tested for associations, with and without a phylogenetic correction. A relaxed multiple test false discovery rate (FDR) (q-value < 0.2) was used to first filter all possible hypothetical associations. At this high threshold, several phylogenetically uncorrected signatures were found, raising the suspicion that phylogenetic artifacts could be at play. However, some of these signatures could be relevant and underlie the clade-specific sensitivity/resistance of the bNAb in question. Thus, to down-select the most statistically robust and relevant signature sites, each selected site was required to meet at least two of the following three criteria: i) contact site (as defined below), ii) at least one phylogenetically corrected signature, and iii) at least one strong signature at q < 0.1. All associations with q < 0.2 at selected sites were used to identify sequence features associated with sensitivity or resistance to the bNAb. The Wilcoxon signatures did not identify any new sites or associations. Env contact sites were defined using the cryo-EM structure of PGT145 in complex with Env trimer (PDB:5V8L, Lee2017 PMID:28423342) and a cutoff of 8.5Å from any antibody heavy atom (HXB2 positions: 120, 121, 122, 123, 124, 125, 126, 127, 128, 129, 130, 156, 157, 158, 159, 160, 161, 162, 163, 164, 165, 166, 167, 168, 169, 170, 171, 172, 173, 174, 175, 184, 191, 192, 200, 201, 305, 306, 307, 308, 309, 312, 313, 314, 315, 321).

#### Soluble HIV-1 envelope trimer generation

HIV Env trimer BG505 DS-SOSIP.664 was produced in transiently transfected 293F cells as previously described (*111, 112*). Briefly, the plasmid encoding the SOSIP trimer and a plasmid encoding furin were mixed at 4:1 ratio and transfected into 293F cells at 0.75 mg plasmid / 1 L cells (1.5 x 10E6 /ml) using 293Fectin (Thermo Scientific) or Turbo293 transfection reagent (Speed BioSystems). Cells were then incubated in a shaker at 120 rpm, 37 °C, 9% CO2. The following day, 80 ml HyClone SFM4HEK293 medium and 20 ml FreeStyle™ 293 Expression Medium were added to each liter of cells. Env trimer protein was purified from the day-7 supernatant with VRC01 affinity chromatography, followed by gel filtration on a Sephadex200 16/60HL column and V3-exposed trimers were removed with negative selection on a 447-52D affinity column. The antigenicity of the trimers was confirmed with a panel of antibodies in the Meso Scale Discovery (MSD) platform.

#### Antibody Fab preparation

Variable regions of the RHA1.V2.01 antibody heavy and light chain genes were synthesized (Genscript) and subcloned into the pVRC8400 vector, in which a HRV3C cleavage site was inserted in the heavy-chain hinge region. The heavy and light chain pair was co-transfected in Expi293F cells (Thermo Fisher) using Turbo293 transfection reagent (Speed BioSystems) as described previously (*113*). The culture supernatant was harvested 6 days post transfection and loaded on a protein A column, the column was washed with PBS, and IgG proteins were eluted with a low pH buffer. The eluted IgG proteins were cleaved by HRV3C, and the cleavage mixture was passed through a protein A column.

#### Cryo-EM data collection and processing

Antibody Fab fragments of RHA1.V2.01 were incubated with BG505 DS-SOSIP with the Fab in molar excess. 2.3 μl of the complex at 1 mg/ml concentration was deposited on a C-flat grid (protochip.com). An FEI Vitrobot Mark IV was used to vitrify the grid with a wait time of 30 seconds, blot time of 3 seconds and blot force of 1. Automated data collection was performed with Leginon (*114*) on a Titan Krios electron microscope equipped with a Gatan K2 Summit direct detection device. Exposures were taken in movie mode for 10 s with the total dose of 71.06 e–/Å^2^ fractionated over 50 raw frames. Images were pre-processed through Appion (*115, 116*); MotionCor2 (*117*) was used for frame alignment and dose-weighting. The CTF was determined using CTFFind4 (*118, 119*). Initial particle picking was done with DoG Picker (*116, 120*). RELION (*121*) was then used for particle extraction. CryoSPARC 2.12 (*122*) was used for 2D classifications, ab initio 3D reconstruction, homogeneous refinement, and nonuniform 3D refinement. Initial 3D reconstruction was performed using C1 symmetry, confirming 1 Fab per trimer and C1 symmetry was applied for the final reconstruction and refinement. Coordinates from PDB ID 6NNF (*123*) and 6CA6 (*58*) were used for initial fit to the reconstructed map. Simulated annealing was performed on the first refinement followed by iterative manual fitting and real space refinement in Phenix (*124*) and Coot (*125*). Geometry and map fitting were evaluated throughout the process using Molprobity (*126*) and EMRinger (*127*). PyMOL (www.pymol.org) was used to generate figures.

#### Expression and crystallization of CH505 gp120:DH650 Fab complex

CH505 gp120 core (residues 44-492, ΔV1-V2 and ΔV3) (*128*) was expressed in HEK293S GnTI^-^ cells and purified by affinity chromatography on Galanthus nivelis lectin (Vector Laboratories) followed by gel filtration on Superdex 200 column (GE). Deglycosylation was carried out with endoglycosidase H (New England biolabs) in deglycosylation buffer (50 mM sodium acetate, pH 6, 5 mM EDTA, 500 mM NaCl, 10uL endoH, 1 μg/μl leupeptin, 1 μg/μl aprotonin) at 37° overnight, followed by buffer exchange to 40 mM Tris HCl, pH 7.4, 1 M NaCl, 2 mM MnCl_2_, 2 mM CaCl_2_ and passage through a concanavalin A column (Sigma) to remove any gp120 that had not been fully deglycoylated by the endoH treatment. The eluate was buffer exchanged by passage over a Superdex200 column (GE) into 2.5mM Tris-HCl pH7.5, 350mM NaCl, followed by concentration to 4mg/ml. The Fab fragment of mAb DH650 was expressed in HEK293T cells, as described (*128*). DH650-gp120 core complex was formed by incubating gp120 core with DH650 in 1:1.3 molar ratio followed by gel filtration on Superdex 200 (GE) column in buffer 2.5mM Tris-HCl pH 7.5, 350mM NaCl. The complex was concentrated to 8.5 mg/ml and crystallized by hanging-drop vapor diffusion in 20% PEG 8K, 100mM Tris pH 8, 500mM NaCl.

#### Determination of CH505 gp120:DH650 Fab crystal structure

X-ray diffraction data to 2.8 Å resolution were collected at the Advanced Photon Source (Argonne National Laboratories) on NE-CAT beamline 24-ID-C. Intensity data were integrated, scaled and merged with XDS and XSCALE (*129*) (see **Table S4**). An initial model, obtained by molecular replacement with Phaser (*130*) using the ZM176.6 gp120 core (PDB ID:4LST), was rebuilt and refined with COOT (*131*) and Buster (Global Phasing, Ltd., Cambridge, UK), respectively. The final model (PDB ID: 6XCJ) includes residues 52-128, 207-299, 327-396 and 409-488 of CH505 gp120, and heavy chain residues 1-223 and light chain residues 1-219 of Fab DH650.

#### Negative-stain electron microscopy of CH505 SOSIP:DH650 Fab complex

CH505 DS-SOSIP, prepared as described (*7*), was mixed with a DH650 Fab in a 1:4 molar ratio. After incubation for 1 hour, the complex was loaded on a Superose 6 column (GE). The SOSIP-Fab complex was diluted to 0.1mg/ml and applied to freshly glow-discharged, carbon-coated EM grids and negatively stained with 1% uranyl formate. Images were recorded on an FEI Tecnai T12 electron microscope, operated at 120 kV and equipped with a ccd detector. Particles were selected and class averages computed with EMAN2 (*132*).

#### Neutralization fingerprint

The neutralization fingerprint of a monoclonal antibody is defined as the potency pattern with which the antibody neutralizes a set of diverse viral strains. The neutralization fingerprints of V2 targeting broadly neutralizing antibodies, including VRC26, PGT145, PG9, and VRC38 classes, as well as a set of other HIV-1 broadly neutralizing antibodies were compared and clustered according to fingerprint similarity, as described previously (*59*), using a panel of 208 HIV-1 viral strains.

#### Statistical analyses

Statistical tests were calculated by using GraphPad Prism 7 software. The Mann-Whitney test was used to determine whether the viral loads (peak and setpoint) of anti-CD8 treated animals were significantly different from untreated animals. We chose a nonparametric rank-based test because both peak and setpoint viral loads of the untreated group failed the D’Agostino & Pearson normality test (P-values < 0.05). The Spearman’s rank correlation test was used to determine if there is a significant correlation between setpoint viral loads and autologous tier 2 NAb titers. The geometric means were calculated using the Column statistics function of GraphPad Prism. The Chi-squared test was used on 2×2 contingency tables of shared and non-shared mutations within or between groups of humans and animals infected by viruses bearing the same (homologous) or different (heterologous) Envs to determine if Env residue mutations demonstrated significant strain-specificity. Separate tests were run to analyze total shared and non-shared mutations for the groups overall and for any discordant CH505, CH848 or CAP256SU group pairing.

#### Data and software availability

GenBank accession numbers for all HIV-1 *env* sequences analyzed in this study are as follows: MN471655-MN472016, MT484339-MT487491, MT580365-MT580445, MT509359, GQ999989, HQ625604, KC863461-KC863464, KF241776, KF996577-KF996601, KF996604, KF996606, KF996610-KF996630, KF996632-KF996662, KF996664-KF996678, KF996680, KF996682-KF996683, KF996685-KF996716, KT698223-KT698227, MF572809-MF572829, EF203980-EF203981, MK205498-MK205507. HIV-1 Env sequences from the human subject CAP256 were previously published (*3, 5, 133*). GenBank accession numbers for immunoglobulin genes are: MT581213-MT581268, XXXX-YYYY. Cryo-EM maps and fitted coordinates have been deposited with database codes EMDB-xxxx and PDB ID xxxx, respectively. DH650 bound to the gp120 core was deposited with database code PDB ID xxxx.

## Supporting information

Supplement to Roark et al

## ACKNOWLEDGMENTS

We thank A. Abuahmad, H. Chen, A. Eaton, A. Foulger, M. Gladden, T. Gurley, G. Hernandez, A. Huang, C. Jones, J. Kim, J. Meyer, A. Monroe, R. Parks, J. Rathmann, P. Rawls, A. Sanzone, J. Sprenz, K. Tilahun, R. Verardi, T. Von Holle, A. Wang, R. Zhang and the flow cytometry core facilities at the Duke Human Vaccine Institute and the Vaccine Research Center (NIAID/NIH) for technical assistance. We thank T. Denny, T. Demarco and N. DeNaeyer and members of the Nonhuman Primate Virology Core Laboratory at the Duke Human Vaccine Institute for SHIV plasma vRNA measurements. We thank J. Baalwa, D. Ellenberger, F. Gao, K. Hong, F. McCutchan, D. Montefiori, L. Morris, J. Overbaugh, E. Sanders-Buell, R. Swanstrom, M. Thomson, S. Tovanabutra, C. Williamson and L. Zhang for contributing HIV-1 Env plasmids and K. McKee, C. Moore, S. O’Dell, G. Padilla, S.D. Schmidt, C. Whittaker, A.B. McDermott and M. Seaman for assistance with neutralization assessments on the 208-strain and 118-strain virus panels. We thank D. Fera for helpful advice and discussion and we thank the staff at Bioqual for exceptional care and assistance with nonhuman primates. Cryo-EM was performed at the Simons Electron Microscopy Center and National Resource for Automated Molecular Microscopy located at the New York Structural Biology Center and was supported by grants from the Simons Foundation (SF349247), NYSTAR, and the NIH National Institute of General Medical Sciences GM103310. X-ray diffraction data were collected on the Northeastern Collaborative Access Team beamline 24 ID-C (AdvancedPhoton Source), funded by NIH Grant P41 GM103403. This work was funded by the Bill & Melinda Gates Foundation (OPP1145046, OPP1206647/INV-007939); by the Intramural Research Program of the Vaccine Research Center, National Institute of Allergy and Infectious Diseases, NIH; and by the Division of AIDS (NIAID/NIH) through support of the Center for HIV/AIDS Vaccine Immunology-Immunogen Discovery (UM1 AI100645), the Consortium for HIV/AIDS Vaccine Development (UM1 AI144371), the Penn Center for AIDS Research (P30 AI045008), and grants AI131251, AI131331, AI128832, AI050529 and AI150590. R. Roark was supported by an NIH training grant in HIV Pathogenesis (T32-AI007632).

## Notes

### Competing Interest Statement

The authors have declared no competing interest.

