## Supplement to Roark et al for "Recapitulation of HIV-1 Env-Antibody Coevolution in Macaques Leading to Neutralization Breadth": SHAW EnvAb Coevoluton.BioRxivFINAL supplement check red.pdf

Short title: Env-Ab coevolution in human and macaque

Ryan S. Roark[1]*, Hui Li[1]*, Wilton B. Williams[2,3]*, Hema Chug[4]*, Rosemarie D. Mason[5]*, Jason Gorman[5]*, Shuyi Wang[1], Fang-Hua Lee[1], Juliette Rando[1], Mattia Bonsignori[2,3], Kwan-Ki Hwang[2], Kevin O. Saunders[2,6], Kevin Wiehe[2,3], M. Anthony Moody[2,7], Peter T. Hraber[8], Kshitij Wagh[8], Elena E. Giorgi[8], Ronnie M. Russell[1], Frederic Bibollet-Ruche[1], Weimin Liu[1], Jesse Connell[1], Andrew G. Smith[1], Julia DeVoto[1], Alexander I. Murphy[1], Jessica Smith[1], Wenge Ding[1], Chengyan Zhao[1], Neha Chohan[1], Maho Okumura[1], Christina Rosario[1], Yu Ding[1], Emily Lindemuth[1], Anya M. Bauer[1], Katharine J. Bar[1], David Ambrozak[5], Cara W. Chao[5], Gwo-Yu Chuang[5], Hui Geng[5], Bob C. Lin[5], Mark K. Louder[5], Richard Nguyen[5], Baoshan Zhang[5], Mark G. Lewis[9], Donald Raymond[4], Nicole A. Doria-Rose[5], Chaim A. Schramm[5], Daniel C. Douek[5], Mario Roederer[5], Thomas B. Kepler[10], Garnett Kelsoe[2,6], John R. Mascola[5], Peter D. Kwong[5], Bette T. Korber[8], Stephen C. Harrison[4,11], Barton F. Haynes[2,3], Beatrice H. Hahn[1], George M. Shaw[1†]

[1]Departments of Medicine and Microbiology, Perelman School of Medicine, University of Pennsylvania, Philadelphia, PA 19104, USA.
[2]Duke Human Vaccine Institute, Duke University School of Medicine, Durham, NC 27710, USA.
[3]Department of Medicine, Duke University School of Medicine, Durham, NC 27710, USA.
[4]Laboratory of Molecular Medicine, Boston Children's Hospital and Harvard Medical School, Boston, MA 02115, USA.
[5]Vaccine Research Center, National Institute of Allergy and Infectious Diseases, National Institutes of Health, Bethesda, MD 20892, USA.
[6]Departments of Immunology and Surgery, Duke University School of Medicine, Durham, NC, 27710, USA.
[7]Departments of Pediatrics and Immunology, Duke University School of Medicine, Durham, NC, 27710, USA.
[8]Theoretical Biology and Biophysics, Los Alamos National Laboratory, Los Alamos, NM 87545, USA.
[9]Bioqual, Inc., Rockville, MD 20850 USA.
[10]Departments of Microbiology, Boston University School of Medicine, and Mathematics & Statistics, Boston University, Boston, MA 02118, USA.
[11]Howard Hughes Medical Institute, Harvard Medical School, Boston, MA 02115, USA.

*These authors contributed equally to this work

#### Supplementary Figure Legends

**Fig. S1. Design and characterization of SHIV.CAP256SU.** (**A**) SHIV.CAP256SU was constructed using a design scheme previously reported (*9*) and represents a chimera of HIV-1 CAP256SU (blue), HIV-1 191859 (red) and SIVmac766 (grey) sequences (GenBank accession #MT509359). Env375-Ser (S) is the wildtype allele found in HIV-1 CAP256SU and in most M group viruses. This residue was replaced with each of five aliphatic, basic or aromatic residues (M, Y, H, W, F). (**B**) SHIVs expressing each of the six Env375 alleles were tested for neutralization sensitivity to a panel of prototypic human mAbs that targeted canonical bNAb epitopes, a linear V3 epitope (447-52D) and a CD4-induced bridging sheet epitope (17b). All six Env375 alleles retained native antigenicity and tier 2 neutralization profiles. (**C**) Replication of SHIV.CAP256SU Env375 variants in primary, activated rhesus and human CD4 T cells is depicted. All six allelic variants replicated efficiently in human cells but only the W, F, Y and H variants replicated in rhesus cells. This result highlights the ability of a single amino acid substitution at position 375 in Env to confer dramatic differences in replication efficiency in primary rhesus CD4 T cells.

**Fig. S2. Virus load and autologous neutralizing antibody kinetics in the plasma of rhesus macaques and humans.** Human and rhesus infections by CH505 (left panels), CH848 (middle panels) and CAP256SU (right panels) viruses are depicted. Animals pretreated with anti-CD8 mAbs are shown in red, untreated animals in blue, and humans in open black boxes. Crosses indicate animals that developed AIDS-defining events requiring euthanasia. Dashed lines demarcate the limits of quantification of the plasma virus load assay (250 vRNA molecules/ml).

**Fig. S3. Glycan shield evolution in CH505 human and SHIV.CH505 infected RMs.** (**A**) Longitudinal glycan frequencies over time for selected glycans that show similar patterns across human (red) and RMs (different shades of blue and green). N332 is gained in all hosts with similar kinetics. N234 is gained in human and most RMs. Both glycans fill rare glycan holes in CH505 T/F. Glycan 130 is lost over time in human and all RMs except RM5695 in which week 2 sequences lack N130 and this glycan is sporadically gained or lost. N130 is not predicted to have an effect on glycan shielding but is typically associated with resistance to V2 apex bNAbs (see **Fig. S26**). (**B**) Longitudinal glycan shield evolution over time. Blue patches indicate the location of time point consensus PNGs and green their predicted glycan shields. Magenta and pink regions indicate glycan holes at regions that are glycan shielded in >80% or 50-80% of M-group sequences, respectively (*45*).

**Fig. S4. Conserved Env evolutionary patterns between the HIV-1 infected human subject CH848 and SHIV.CH848 infected rhesus macaques.** A complete description of the PIXEL plots (left panels) and the LASSIE analysis (right panels) is provided in the legend to **Fig. 2**. Sixteen of 29 residues that mutated to >75% fixation in the human subject CH848 also mutated in one or more monkey and these are highlighted by green flags in the PIXEL plots and by green lettering in the LASSIE plot. Critical deletions in V1 associated with the development of neutralization breadth in the human subject CH848 and in RMs 6163 and 6167 are indicated by a yellow flag.

**Fig. S5. Glycan shield evolution in CH848 human and SHIV.CH848 infected RMs.** (**A**) Common patterns of glycan shield evolution between CH848 human and RM6167 and RM6163. Longitudinal evolution of the viral quasispecies in CH848 human resulted in two glycan changes by week 75. The loss of N413 glycan does not impact the glycan shield but could alter sensitivity to autologous V3 NAbs. The loss of N289 creates a glycan hole. Glycan shield evolution in RM6167 at week 28 and in RM6163 at week 80 shows similar changes. (**B**) Longitudinal glycan frequencies over time in human and RMs for selected glycans that show similar patterns across human (red) and RMs (different shades of blue and green). N413 is lost in all hosts with similar kinetics. N289 is transiently or permanently lost in human and most RMs. (**C**) Longitudinal glycan shield evolution over time in each host. Glycan shield calculation and coloring is the same as in **Fig. S3B**.

**Fig. S6. Selected sites identified by LASSIE in human subjects CH505 and CH848 are rarely mutated in RMs infected by SHIVs bearing heterologous Envs.** The uppermost LOGO plots at the top of this illustration depict HIV-1 Env residues in the human subjects CH505 (panel A) and CH848 (panel B) that were identified by LASSIE analysis to be under positive selection (**Figs. 2** and **S4**). The position of each of these residues (HXB2 numbering) is shown at the bottom of each panel. Only four residues were identified to be under positive selection in both human subjects, and these positions were 130, 300, 413 and 620 (indicated by the vertical red lines). The lower LOGO plots residues in the heterologous T/F Env (CH848 - panel A; CH505 - panel B), and these were tracked over time in the SHIV infected monkeys for comparison to the matched analyses presented in **Figs. 2** (CH505) and **S4** (CH848). Over each animal's LOGO display, there are two numbers in green at the upper right. The first digit shows the total number of sites under positive selection in the Env mismatched human and monkey, including sites highlighted in red. The second digit shows the number of sites under positive selection in the Env mismatched human and monkey *after* excluding the shared sites. Excluding the shared sites, we compared the patterns of selection in the different groups of animals. We found four of six animals infected with CH848 had no mutations in selected sites from the human subject CH505, RM6700 had 1 shared site, and RM6167 had 2 (panel A). In a comparison with **Fig. 2** from the main text, 3 of the six animals shared 9 sites, two shared 7 sites, and one shared 6 sites. A Wilcoxon test showed that the distribution of CH505 selected sites in SHIV.CH505 macaques (9, 9, 9, 7, 7, 6), versus six SHIV.CH848 macaques (2, 1, 0, 0, 0, 0), was highly significant ($p = 0.002$). We also compared the human CH848 selected sites shared among the six SHIV.CH505 macaques shown in part B (3, 2, 1, 0, 3, 7 shared sites) with the six SHIV.CH848 infected macaques (**Fig. S4**) (11, 11, 10, 6, 9, 4), and again found the distributions to be significantly different ($p = 0.006$). Note that because the human CAP256 was infected with two divergent HIV-1 strains, this control analysis was not applicable for this SHIV.

**Fig. S7. Conserved Env evolutionary patterns between the HIV-1 infected human subject CAP256 and SHIV.CAP256SU infected rhesus macaques.** (**A**) PIXEL plot highlighting amino acid mutations between the SHIV.CAP256SU infecting strain and subsequent samples in each RM. Red pixels indicate a change in amino acid relative to the infecting strain, and black indicates an INDEL, as in **Figs. 2** and **S3**. Green tags indicate residue positions that are selected in multiple RMs. Black numbers indicate HXB2 numbering and white numbers approximate HXB2 numbering in hypervariable regions. Yellow tags indicate identical INDELS. (**B**) LASSIE analysis of the same longitudinal Env RM sequences, summarizing mutations under positive selection in any of the four RMs with a threshold of 75% replacement of the infecting CAP256SU strain shown at the top. Green numbers represent mutations observed in more than one animal. (**C**) Recombination map of CAP256SU and CAP256PI. The primary infection (PI) variants are shown as solid blue lines at week 6 and the superinfection (SU) variants as red lines at week 15. In subsequent time points, amino acids where the two forms match are grey. Places where they differ are indicated by blue if they match the PI, red if they match SU, and black if the differ from both. Extensive recombination is seen as early as week 23. (**D**) The mutations under selection in the RMs shown in panel **B** are tracked in the human relative to the CAP256SU form that dominates at w015. Positions under selection in the RMs that differ in the SU and PI forms are evident as the w006 forms are different than SU reference amino acids along the top. The mutations that arose *de novo* in the macaques were also often favored in the human and were most often explained by recombination of the PI forms into the SU backbone (e.g., E87K and T525A), although sometimes the mutation selected in RMs arose independently in the human and was not evident in either the PI or SU forms (e.g., R46K and K340E). Mutations at residue 169 in the human and in RMs 40591 and 42056 conferred escape from V2 apex targeted bNAbs (**Fig. 3B,C**).

**Fig. S8. Glycan shield evolution in CAP256 human and SHIV.CAP256SU infected RMs.** (**A**) Common patterns of glycan shield evolution between CAP256 human and RM40591. CAP256SU has a glycan hole due to absence of N339. During longitudinal evolution in the human and RM40591, N396 is gained over time. (**B**) Longitudinal glycan frequencies over time in human and RMs for N396 that shows similar patterns across human (red) and RMs (different shades of blue and green). (**C**)

Longitudinal glycan shield evolution over time in each host. Glycan shield calculation and coloring same as **Fig. S3B**.

**Fig. S9. Summary of indels in the CH505 human and SHIV.CH505 infected RMs.** The HXB2 reference and CH505T/F sequences are indicated at the top for orientation. Hypervariable regions within the loops are shown in red. Each indel is aligned independently to the T/F as multiple alignments obscure the precise nature of the indel. Precise indel boundaries are indicated and unique indels are assigned a color. Only indels that were found in the human or repeated in multiple individuals are shown. With the exception of a single codon deletion (in red), all V1 indels were insertions. The T/F is shown at the top and the indel beneath it. The translated sequences are depicted on the right. Insertions are generally direct repeats, and the section that is repeated is highlighted (e.g. the 9 bases in green in V1) and then shown repeated in the inserted sequence and translated on the right. The V1 insertions almost always included the addition of at least one N-linked glycosylation site. The V5 deletions always changed the relative location of a glycan within the V5 loop. The distinct hosts that carried a precise indel are indicated by the colored boxes on the right. The vast majority of V1 indels were insertions in the seven CH505 Env infections, and the histogram on the right shows that insertions in V1 restore it to a more typical length. The distribution represents V1 loop lengths based on the global Env M group alignment from the Los Alamos HIV database (www.hiv.lanl.gov). CH505 T/F was unusually short in V1 (20 amino acids long, magenta column), while the median V1 length was 30 amino acids (red column). Similarly, the deletions found in the long CH505 T/F in V4 bring the viral quaisispecies closer to the median length of 31, and the repeated insertions in V5 bring the short CH505 TF V5 closer to a typical length. Only four time points collected during the first year of infection are shown here for simplicity; after the first year of infection, indels on top of indels begin to accrue and the evolutionary history becomes unresolvable.

**Fig. S10. Summary of indels in the CH848 human and SHIV.CH848 infected RMs.** The figure organization is the same as described in the legend to **Fig. S9**. Unlike CH505, the V1 length in CH848 T/F was atypically long and all Indels were comprised of deletions bringing the V1 loop length closer to the global median. Ten Indels in V1, V4 or V5 were precisely replicated between the human subject CH848 and one or more macaques. Additional Indels were replicated among macaques only.

**Fig. S11. Epitope mapping of autologous neutralizing antibody responses in the SHIV.CH505 infected RM6072.** (**A**) Mutations at residues N130, T234, N279, V281, K302, Y330, N334, H417 and K460 were strongly selected in human CH505 and in RM6072 (see **Fig. 2A**). Here, the left panel depicts fold changes in NAb titers of week 10 and 20 plasma against these mutant versus wildtype CH505 T/F Env pseudotyped virus. The right panel shows neutralization curves of RM6072 wk20 plasma against CH505 T/F and single and combination mutants that evolve in human and rhesus. MLV is a murine leukemia virus control for non-specific inhibition of virus entry. (**B**) Left, neutralization of wildtype CH505 T/F and observed *in vivo* mutants by autologous monoclonal antibodies (mAbs) DH647 and DH648 isolated from RM6072. Neutralization potency is expressed as $IC_{50}$ values (μg/ml). Right, neutralization curves of mAbs against CH505 T/F and single and combination mutants. (**C**) Neutralization of wildtype CH505 T/F and observed *in vivo* mutants by mAb DH650 isolated from RM6072 and its inferred germline unmutated common ancestor. mAbs isolated from human subject CH505 (*6*) are included as controls. Neutralization titers are shown as $IC_{50}$ values (μg/ml).

**Fig. S12. Epitope mapping of heterologous neutralizing activity in plasma from RMs infected by SHIVs CH505 (RM5695 and RM6070) or CAP256SU (RM40591 and RM42056).** (**A**) RM5695, **B**) RM6070, **C**) RM40591, **D**) RM42056. Neutralization was tested in the TZM-bl assay using plasma or isolated plasma IgG against viruses expressing wildtype Envs or Envs with mutations at residues 156, 166, 169 or 171. In a diverse set of heterologous Env backbones (Q23.17, T250, BG505, 246F3, MT145K, CE0217 and ZM233), bNAb activity was found to be dependent on basic residues (R or K) at positions 166 and 169, and to a lesser extent, on a PNG at position 156 and a basic residue at position

171. These results, together with *in vivo* escape mutations at residues 166, 169 and 171 (**Fig. 3B**), map the bNAb activity in these plasma specimens to the V2 apex.

**Fig. S13. Variable effects of N160K substitution on heterologous neutralization in RMs infected by SHIVs CH505 (RM5695 and RM6070) or CAP256SU (RM40591 and RM42056).** (**A**) RM5695, (**B**) RM6070, (**C**) RM40591, (**D**) RM42056. Neutralization was tested in the TZM-bl assay using plasma or isolated plasma IgG against viruses expressing wildtype Envs or Envs with N160K substitutions in a diverse set of heterologous Env backbones (Q23.17, T250, BG505, 246F3, MT145K, and ZM233). Plasma from every animal potently neutralized viruses expressing heterologous wildtype Envs. However, the effects of N160K substitution ranged from elimination of neutralization activity (e.g, RM5695 plasma against HIV-1 BG505) to enhancement of neutralization (RM5695 plasma against HIV-1 Q23.17) (panel **A**). In some animals, the enhancement of neutralization against heterologous N160K containing viruses was extraordinary [e.g., RM6070 (panel **B**) and RM42056 (panel **D**)]. N160K dependent neutralization was strictly dependent on antibody binding to basic C-strand residues as shown by N160K.K169E double mutants (panels **B** and **D**). The most likely explanation for these results is that SHIV-infected animals contain three specificities of V2 apex targeted C-strand dependent neutralizing antibodies. The first are the typical V2 apex bNAbs that depend on interactions with C-strand residues and N160 glycans. The second C-strand targeted bNAbs that do not require binding to N160 and instead are enhanced in their potency by the absence of N160 glycans. The third are "2909-like" antibodies (*83, 84, 135*) that are strain-specfic and entirely lacking in neutralization breadth, but whose potency against the homologous virus strain is enhanced by the absence of N160 glycans. We have evidence for the existence of all three types of V2 apex NAbs in SHIV-infected RMs (see Supplement Extended Discussion).

**Fig. S14. Epitope mapping of autologous neutralizing antibody responses in the SHIV.CH848 infected RM6163 and RM6167.** (**A**) At weeks 10 and 36 post SHIV infection, neutralizing antibody titers (reciprocal $ID_{50}$) against the T/F virus reached 1:181 and 1:435, respectively. A naturally-occurring 10 amino acid deletion in V1 resulted in a 30% reduction in neutralizing antibody titer in the week 10 plasma but enhanced neutralization titers in concurrent wk 36 plasma. This V1 deletion, together with week 36 mutations at positions 169, 240, 290, 336, 339, 413 and 65, led to complete escape from neutralizing antibodies in week 10 and 36 plasma, indicating again that selected mutations in Env generally arise as a consequence of neutralizing antibody pressure. (**B**) A six member autologous strain-specific neutralizing Ab lineage isolated from RM6163 (*136*) potently neutralizes CH848 T/F and is strictly dependent on N332 for activity. The naturally-occurring V1 deletion in CH848 T/F (**panel A**) reduces neutralization titers by five-fold, similar to its effects on the human mAb DH475 [see main text (*7*)].

**Fig. S15. Characterization of CD4bs antibodies in RM6072.** (**A**) Plasma antibody binding to wildtype and CD4-binding site mutant (Δ371I) variants of CH505 TF gp120 and resurfaced core 3, RSC3 (*95*). Bottom panels depict differential binding ratios. ELISA results are from a representative experiment. **(B)** Plasma antibody blocking of Env binding by CD4bs bnAbs CH235 and CH106 (from the CH103 lineage) and by soluble CD4 (sCD4) as determined by competitive ELISA in a representative experiment. **(C)** Phylogram of memory B cell-derived antibodies designated the "DH650 lineage" that were isolated from blood (black) and bone marrow aspirate (purple) cells by an antigen-specific B cell sort approach or from blood by memory B cell culture (blue). The unmutated common ancestor (UCA) was computational inferred using the Cloanalyst software of macaque immunoglobulin reference genes (*24*). **(D)** Immunogenetics of the DH650 heavy and light chain genes was inferred by the Cloanalyst software. **(E)** Binding of DH650 mAbs to CH505 T/F gp120 was measured by ELISA and titers reported as $EC_{50}$ (μg/ml). **(F)** DH650 mAbs were tested by ELISA for their ability to block Env binding by CH235, CH106 and sCD4. Inhibition is reported as percent blocking in a representative experiment.

**Fig. S16. Binding and neutralization profiles of DH650 lineage antibodies. (A)** Binding of DH650 mAbs to CH505 T/F gp120 and CH505 T/F SOSIPS was assessed by biolayer interferometry (BLI). mAb-captured sensors were placed into solutions of Envs at 50 μg/ml to measure binding depicted as response units over time. **(B)** Neutralization by DH650 lineage mAbs in TZM-bl cells against wildtype

CH505TF and mutant CH505TF depleted of four glycans surrounding the CD4bs (*137*). IA refers to computationally inferred intermediate antibodies. Murine leukemia virus (MuLV) was used as a negative virus control. **(C)** Neutralization by TZM-bl assay of DH650 lineage mAbs against heterologous HIV-1 strains previously used to track neutralization breadth in the plasma of the HIV-1 infected CH505 human subject (*2*). Neutralization results were confirmed and extended in a test of 118 heterologous virus strains (**Table S3**).

**Fig. S17. Genetic and biological characterization of DH650 lineage. (A)** The HCDR2 region of the VH gene of CH235 and 8ANC131 plays a critical role in engaging the CD4 binding loop of HIV-1 Env. Asterisks identify CH235VH Env contact residues. Dots indicate identical residues at each position and dashes indicate deletions. A residue 57 serine (S) to arginine (R) mutation in the CH235 UCA was found to be critical for binding of CH235 lineage bNAbs to HIV-1 Envs (Bonsignori Cell 65:449, 2016). We found a similar mutation at residue 57 asparagine (N) to R in the DH650 lineage NAbs. (**B**) Binding by ELISA of DH650 mAb to CH505 T/F Env monomer and SOSIP depends on R57 in CDRH2. Controls included CH235, 2G12 and influenza-reactive mAb CH65. (**C**) Complete VH sequences of DH650 lineage antibodies isolated from single cell sorts of blood derived B cells 20, 24 and 32 weeks post SHIV infection (DH650-DH650.13) and sequences generated by next generation sequencing from blood mononuclear cells 52 weeks post SHIV infection (DH650.15-34_NGS). **(D)** DH650.6, which contained VH residues S35 and P62, failed to bind CH505 T/F Env (**Figs. S15E, 16A**) or neutralize CH505 T/F or T/F-gly4 virus (**Figs. S16B,C**). Substitution of either S35 to N35 or P62 to Q62 restored functionality as measured by ELISA binding to CH505 T/F gp120 (**Fig. S17D**). However, for DH650.15_NGS and DH650.16_NGS heavy chains paired with the DH650 VL chain, N35 and Q62 were not sufficient to support Env binding, suggesting that authentic paired VL chains are essential for DH650.15-16 functionality. (**E**) VL chain sequences of DH650 lineage antibodies compared with CH235 UCA and mature bNAbs. DH650 lineage antibodies contained an exceptionally long LCDR1 of 17 amino acids compared with 11 amino acids in CH235 antibodies.

**Fig. S18. Heavy and light chain sequences of RHA1 bNAbs from RM5695.** (**A**) Nucleotide and amino acid alignments of rhesus mAb heavy chains RAH1.V2.01-04 compared with RM5695 germline gene IGHV4-ABB-5*01_S8200. (**B**) Nucleotide and amino acid alignments of rhesus mAb light chains RAH1.V2.01-04 compared with RM5695 germline gene IGLV1-ACN*02.

**Fig. S19. Structural, immunogenetic and neutralization properties of RHA1 bNAbs.** (**A**) The two residue insertion in the HCDR1 region of RHA1 bNAbs results in interactions with N160 and electrostatic contact between D29 and Env residue K171. (**B**) IGHV4-ABB-5*01_S8200 (blue), DH3-9 (red) and JH2-P (teal) plus six non-templated amino acids (black) contribute to the RHA1 HCDR3. Somatic hypermutation resulting in amino acid substitutions in D and J genes of the mature bNAb mAbs are shown. The amino-terminal Cys and carboxy-terminal Trp (underlined) are not part of the HCDR3 (IMGT). Dots indicate amino acid identities. (**C**) RHA1 mAbs are aligned to PCT64-35S and PGT145. RHA1 and PCT64-35S are similar in length (24 vs 25 amino acids, respectively). Seven of 24 RHA1 HCDR3 amino acids (yellow shading) are identical to those in PCT64-35S, including a critical DDY motif that is tyrosine-sulfated (red asterisks) in both antibody lineages and in PGT145. Blue shading highlights nonidentical residues that share chemical properties such as aromaticity or charge. Tyrosine sulfation plays a key role in binding HIV-1 Env by all three antibodies. (**D**) The neutralization fingerprint for RHA1.V2.01 illustrated in **Fig. 6E** showed it to cluster within the PGT145 class of bNAbs, which clustered significantly with all prototypic V2 apex bNAbs analyzed. Here, we show the additional 21 non-V2 apex targeted bNAb antibodies that were part of the same neutralization fingerprint analysis. (**E**) Venn diagram of the number of neutralization sensitive HIV-1 strains for RHA1.V2.01, CAP256-VRC26.25 and PGT145. Of the 101 viruses neutralized by RHA1.V2.01, 96 were neutralized by PGT145 and 84 were neutralized by CAP256-VRC26.25, supporting the neutralization fingerprint dendogram shown in **Fig. 6E**.

**Fig. S20. RHA1.V2.01 lacks autoreactivity.** (**A**) Indirect immunofluorescence showing lack of anti-nuclear antibody (ANA) reactivity by mAb RHA1.V2.01 in HEp-2 cells. The positive control mAb is DH1037 and negative control mAb is DH570.30 (50 µg/mL for each). The secondary antibody was a goat anti-rhesus Ig(H+L) FITC. (**B**) RHA1.V2.01, along with positive (4E10) and negative (CH65) control mAbs, were tested for reactivity to nine autoantigens in the ZEUS AtheNA Multi-Lyte ANA-II Plus Test System, according to manufacturer's recommendations. All antibodies were 2-fold serially diluted starting at 50 µg/mL.

**Fig. S21. Heterologous neutralization by RHA1 mAbs.** Rhesus mAbs were tested for neutralization against the autologous CH505 TF virus and the same global panel of heterologous viruses used to test all macaque plasma in this study (**Figs. 1**). Neutralization titers are reported as $IC_{50}$ (µg/ml) values. This same panel of viruses was also tested for neutralization by the RM5695 plasma from wk56 post infection, expressed as reciprocal dilution $ID_{50}$ (µg/ml) values. The breadth of the RHA1 mAbs recapitulated the breadth observed in the polyclonal plasma.

**Fig. S22. RHA1.V2.01 neutralization of a global panel of 208 heterologous HIV-1 viruses.** Neutralization titers are reported as $IC_{50}$ (µg/ml) values. This data is reflected in the dendrogram in **Fig. 6B**. 49% of viruses were neutralized at a concentration <50 µg/ml.

**Fig. S23. Heterologous neutralization of RHA1 lineage bNAbs maps to the V2 apex.** In multiple heterologous Env backbones, neutralization maps to N160, R166 and C-strand residues R/K169.

**Fig. S24. Neutralization profiles of RHA1.V2.01 and other V2 apex bNAbs.** (**A**) $IC_{50}$ titers for RHA1.V2.01 and prototypic human V2 apex bNAbs are shown as a heatmap. Rows indicate viruses from the 208 strain panel (**Fig. S22**) and columns the bNAbs tested. $IC_{50}$ titers are color-coded according to potency (red high, yellow low, blue >50µg/ml). Viruses and bNAbs are ordered using a hierarchical clustering on the Heatmap webtool (https://www.hiv.lanl.gov/content/sequence/HEATMAP/heatmap.html) (*60, 109-113*) using the Ward algorithm on Euclidean distances of $Log_{10}$ transformed $IC_{50}$ titers. A hierarchical clustering tree shows relatedness between $IC_{50}$ profiles of different V2 apex bNAbs. Bootstrap confidence (% of 1000 bootstraps) is indicated on the splits. (**B**) Contingency tables highlight the overlap between sensitive and resistant viruses between RHA1.V2.01 and other V2 apex bNAbs. Resistant viruses had $IC_{50}$ titers above the threshold. Below each table are listed Fisher's exact test p-values, odds ratios (OR) and accuracy estimates (Acc), which measure the fraction of concordant viruses (i.e. sensitive or resistant to both bNAbs). (**C**) Scatterplots of $IC_{50}$ titers with RHA1.V2.01 titers on the y-axis and other V2 apex bNAb titers on the x-axis. Viruses resistant to one but not the other bNAb are shown above or to the right of the dotted lines. The number in the top right indicates the number of viruses resistant to both. Pearson $R^2$ and p-values are listed in the bottom right of each panel and were calculated using viruses that were sensitive to both bNAbs. These analyses together show that RHA1.V2.01 is most similar to PCT64-35M.

**Fig. S25. Neutralization profiles of RHA1.V2.01 and other V2 apex bNAbs across HIV-1 group M subtypes.** Neutralization $IC_{50}$ titers are shown for the 208 virus global panel, broken down according to subtypes. $IC_{50}$ titers are color-coded from red to green indicating potent to less potent $IC_{50}$ titers, with black cells indicating $IC_{50}$ titers greater than maximum concentration tested (50µg/ml). V2 apex bNAbs are grouped according to similarities in their subtype-specific neutralization profiles into three sub-groups: a) PCT64-35M and CAP256 bNAbs; b) CH01; and c) PG9, PGT145 and PGDM1400. The rhesus bNAb RHA1.V2.01 is most similar to the PCT64-35M subgroup.

**Fig. S26. Signature analyses for RHA1.V2.01 and other V2 apex bNAbs.** (**A**) Signature analyses for RHA1.V2.01 were performed as described (*60*). Significant signatures had two or more of the following: i) phylogenetic correction at false-discovery rate (FDR, q) < 0.2; ii) contact site; and iii) at least one strong signature (q < 0.1). This analysis identified sites 130, 160, 166, 167 and 169 as signature sites for RHA1.V2.01. The variability at these sites in M-group is shown using sequence logo in the left panel. On the right, variability in RM5695 longitudinal sequences is shown. Height of the letters indicate the frequency, and O is a potential N-linked glycan site (NXT/S). Blue letters are associated with sensitivity, red resistance and grey neutral. (**B**) Comparison of RHA1.V2.01 signatures to those of other V2 apex bNAbs. The results at signature sites are shown for each bNAb on a single row, using amino acids colored according to sensitivity or resistance and with the height of letters indicating the frequency in M-group sequences (www.hiv.lanl.gov). Notably, only PCT64-35M shared all signature sites for RHA1.V2.01.

**Figure S27. Cryo-EM Details of RHA1.V2.01 in complex with HIV-1 Env BG505 DS-SOSIP.** (**A**) Representative micrograph and contrast transfer function of the micrograph are shown. (**B**) Representative 2D class averages are shown. (**C**) The orientations of all particles used in the final refinement are shown as a heatmap. (**D**) The gold-standard fourier shell correlation resulted in a resolution of 3.90 Å using non-uniform refinement with C1 symmetry. We note that local refinement (beta) of the same particles resulted in a resolution of 3.8 Å. (**E**) The local resolution of the full map is shown generated through cryoSPARC using a Fourier shell correlation (FSC) cutoff of 0.5. Two contour levels are shown. (**F**) Representative density is shown for the Fab (top left) with a magnified region of the inserted sulfated tyrosine, Tys (top right), and for N160 of each protomer labeled P1, P2 and P3 (bottom panels).

**A**

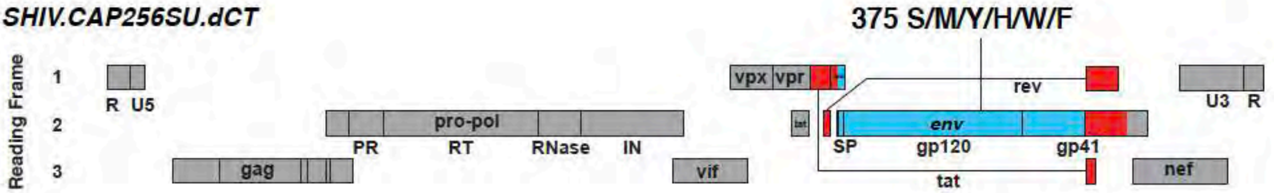

**B**

SHIV.CAP256SU Env375 variant

| | 375S | 375M | 375Y | 375H | 375W | 375F |
|---|---|---|---|---|---|---|
| PGT121 | 0.051 | 0.042 | 0.046 | 0.044 | 0.036 | 0.031 |
| 8ANC195 | 13.3 | 16.3 | 13.9 | 17.3 | 22.4 | 17.3 |
| 10E8 | 5.9 | 6.9 | 9.2 | 12.6 | 10.6 | 5.2 |
| VRC01 | 0.11 | 0.14 | 0.22 | 0.21 | 0.52 | 0.77 |
| PG9 | 0.038 | 0.050 | 0.053 | 0.048 | 0.063 | 0.038 |
| PGDM1400 | 0.005 | 0.007 | 0.005 | 0.006 | 0.010 | 0.006 |
| VRC26.25 | 0.0003 | 0.0004 | 0.0004 | 0.0006 | 0.0006 | 0.0003 |
| 447-52D | >25 | >25 | >25 | >25 | >25 | >25 |
| 17b | >25 | >25 | >25 | >25 | >25 | >25 |

| <0.1 | 0.1-1.0 | 1.0-10 | 10-50 |
|---|---|---|---|

IC50 ug/mL

**C**

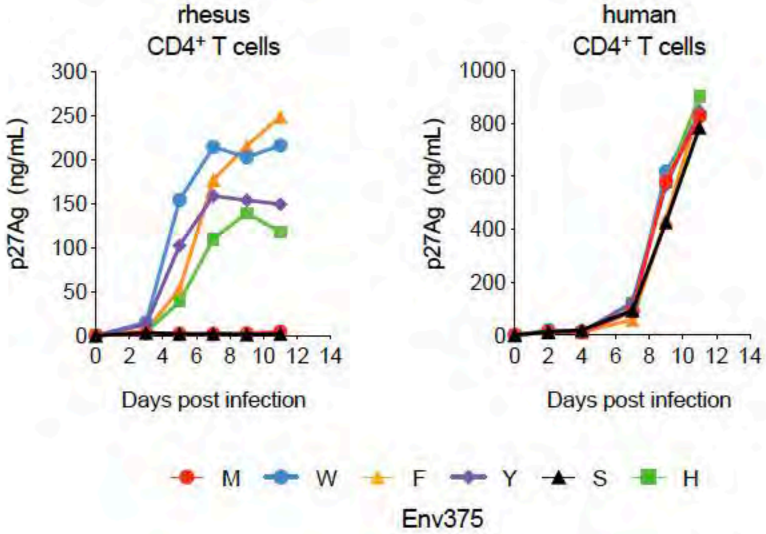

**Supplemental Figure S1**

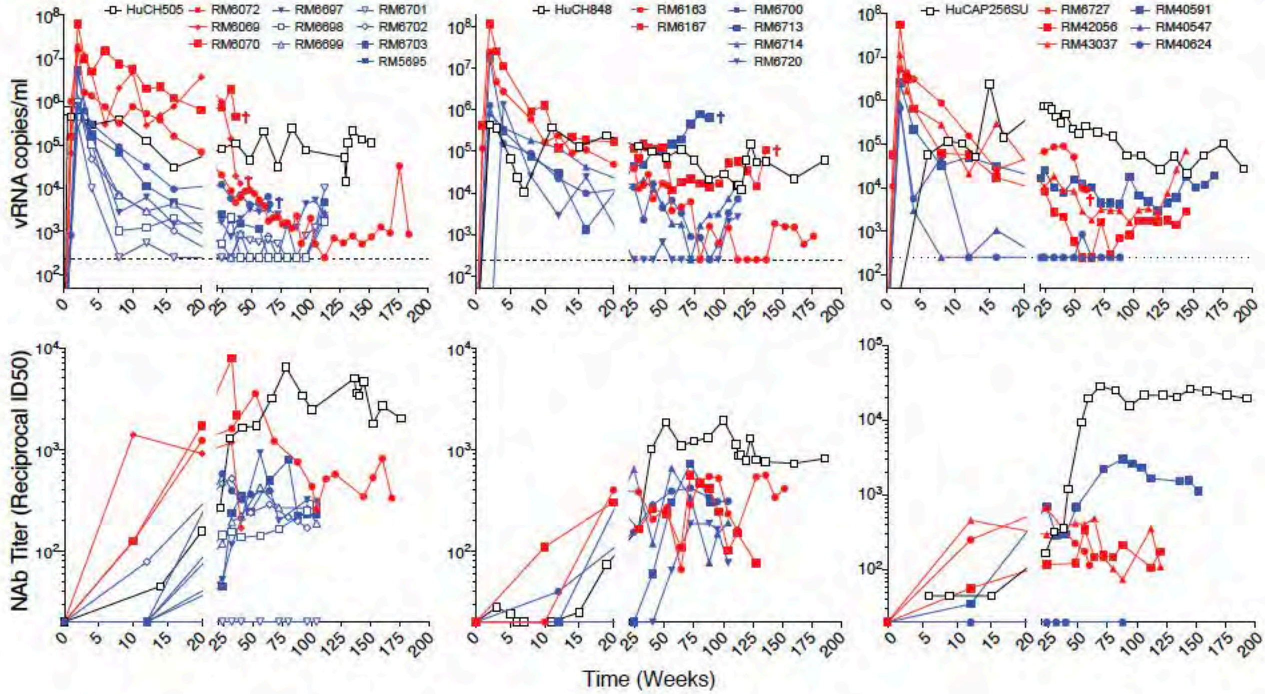

**Supplemental Figure S2**

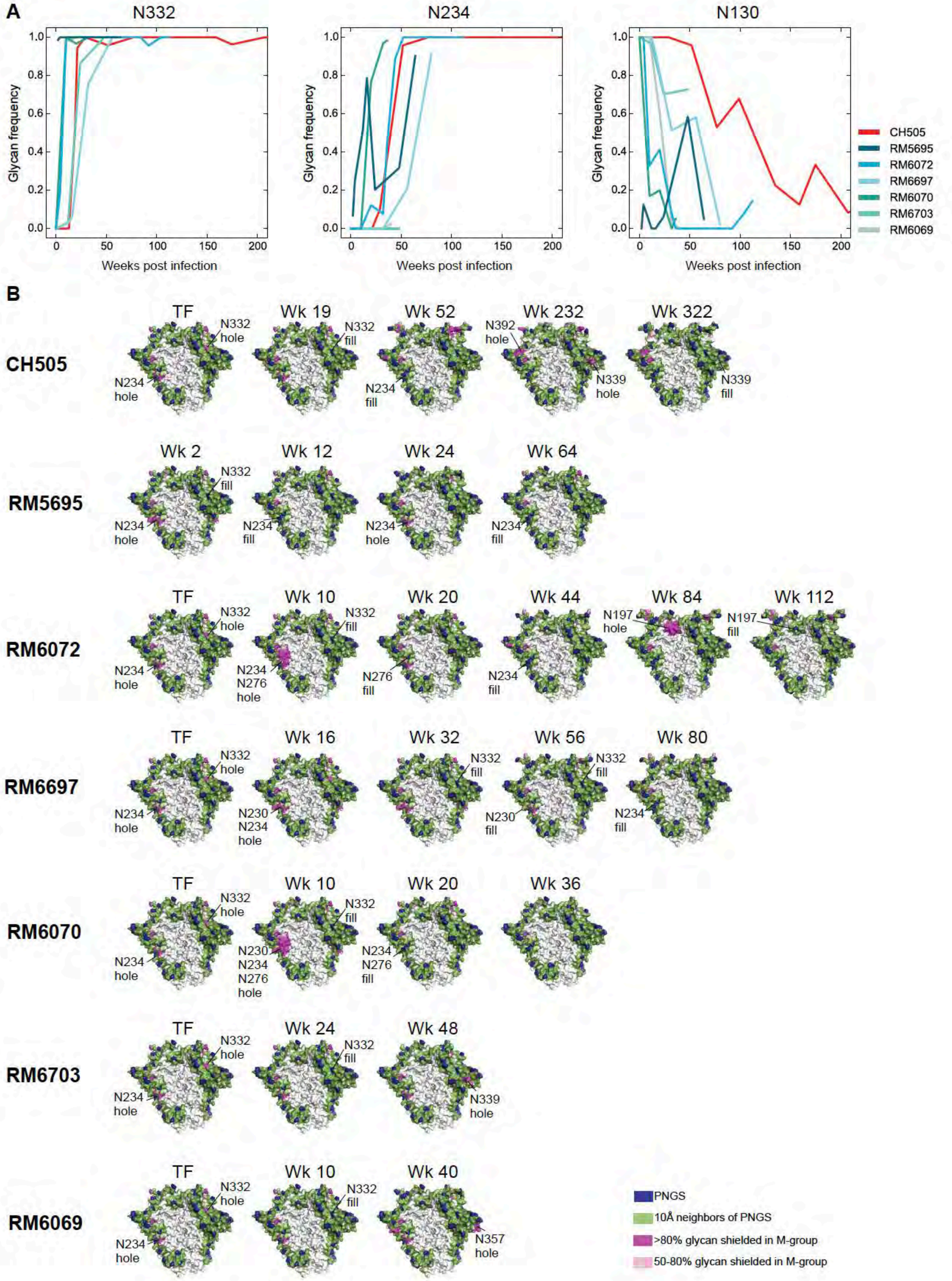

### Supplemental Figure S3

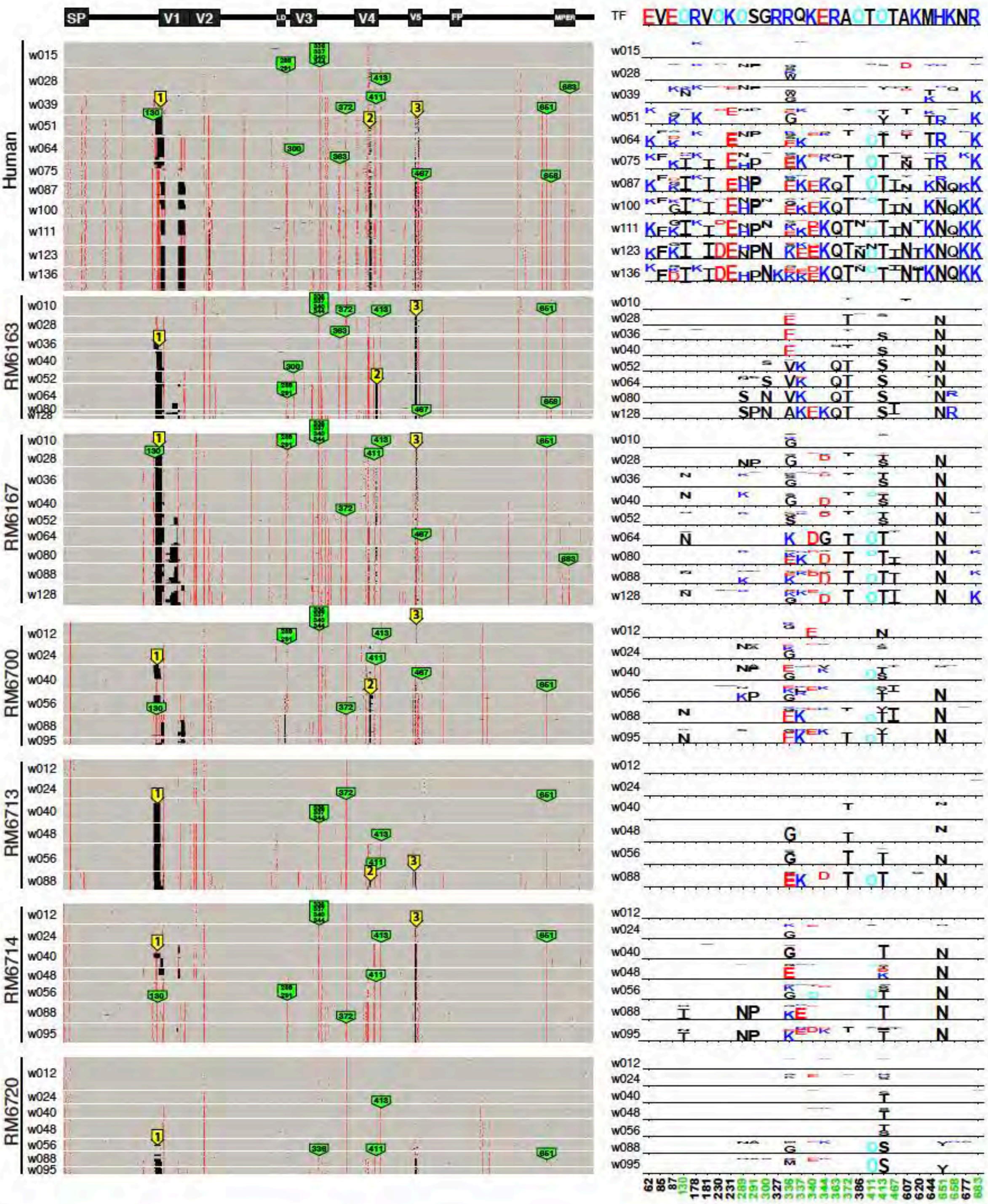

**Supplementary Figure S4**

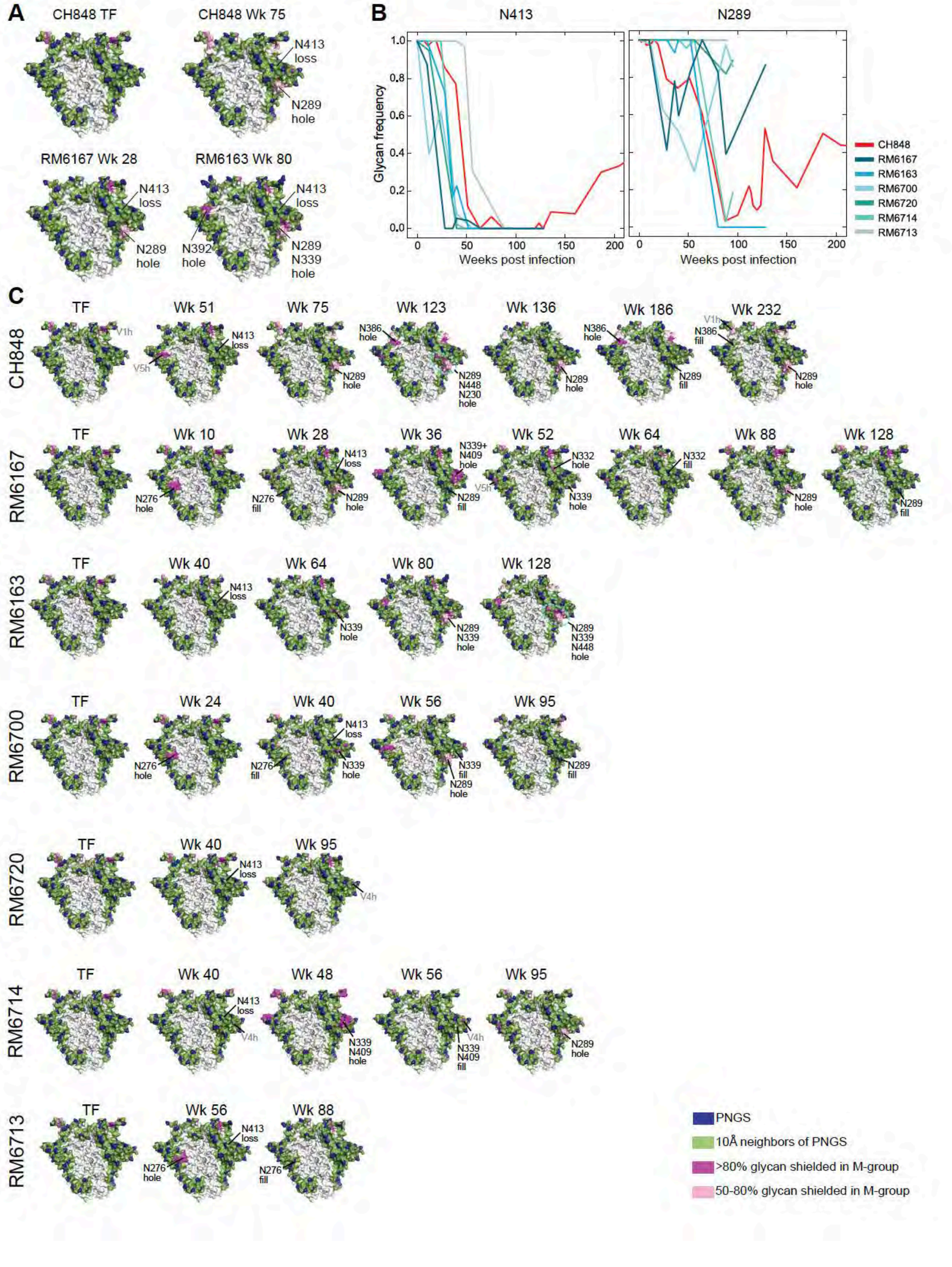

### Supplemental Figure S5

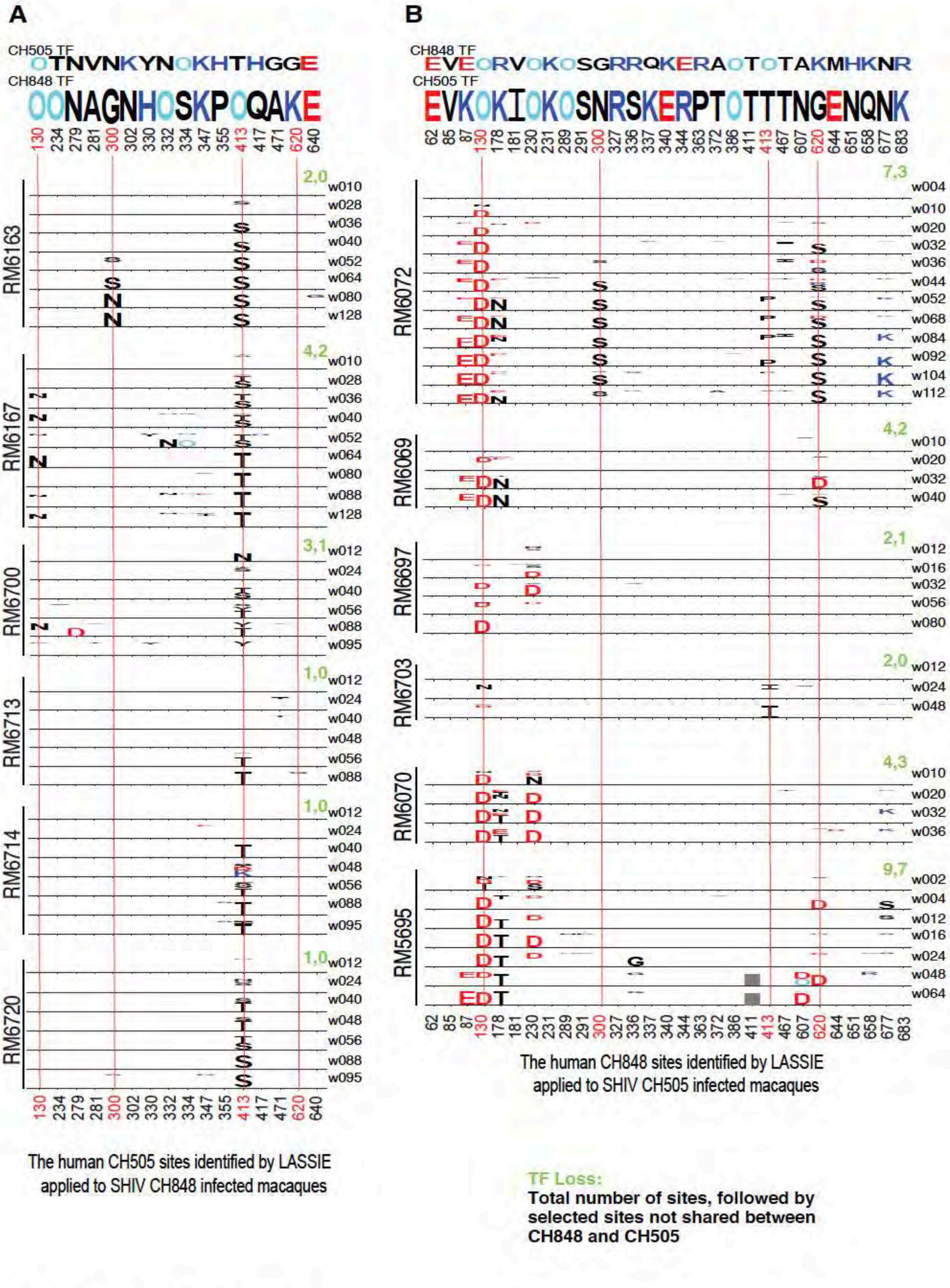

**Supplemental Figure S6**

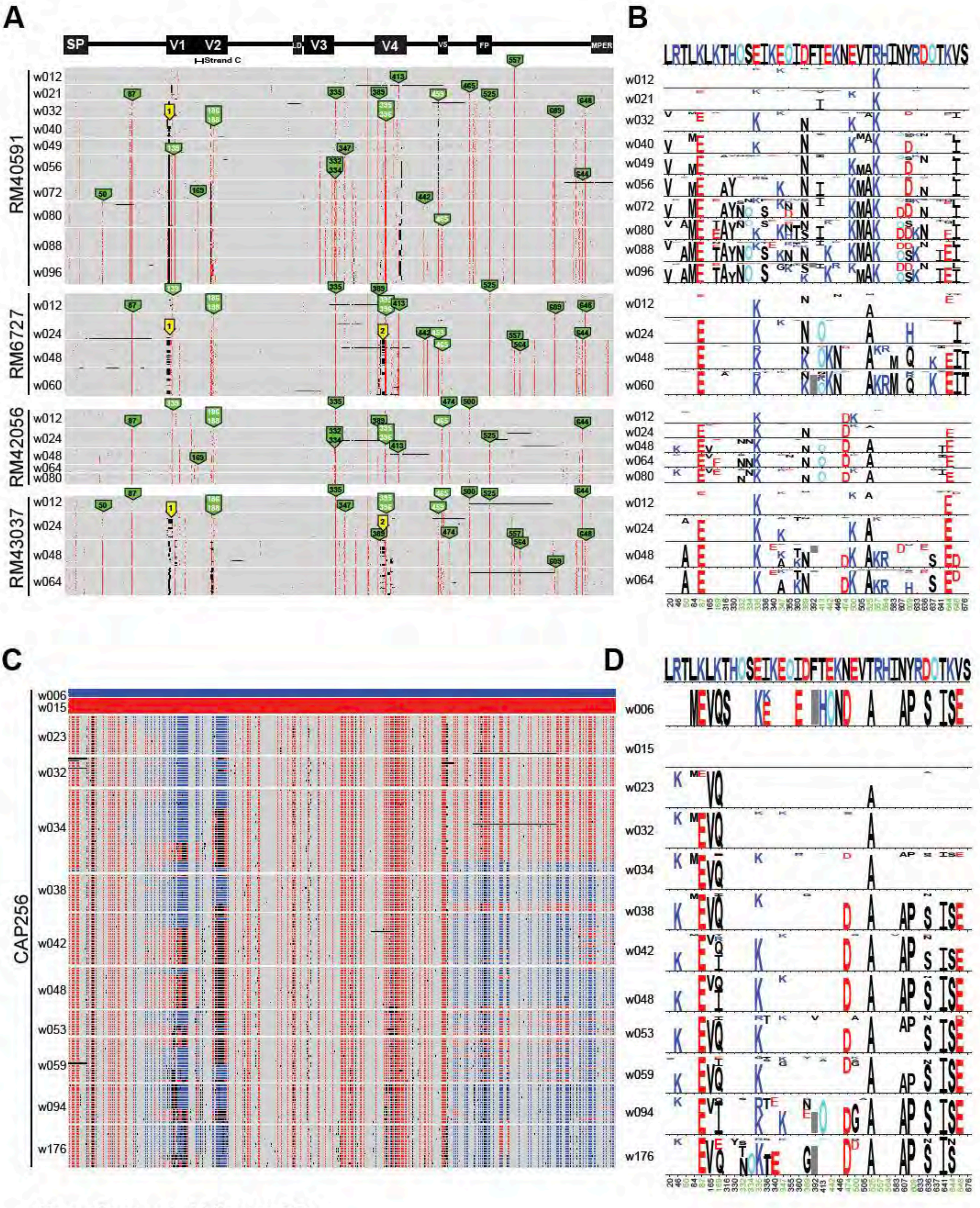

**Supplemental Figure S7**

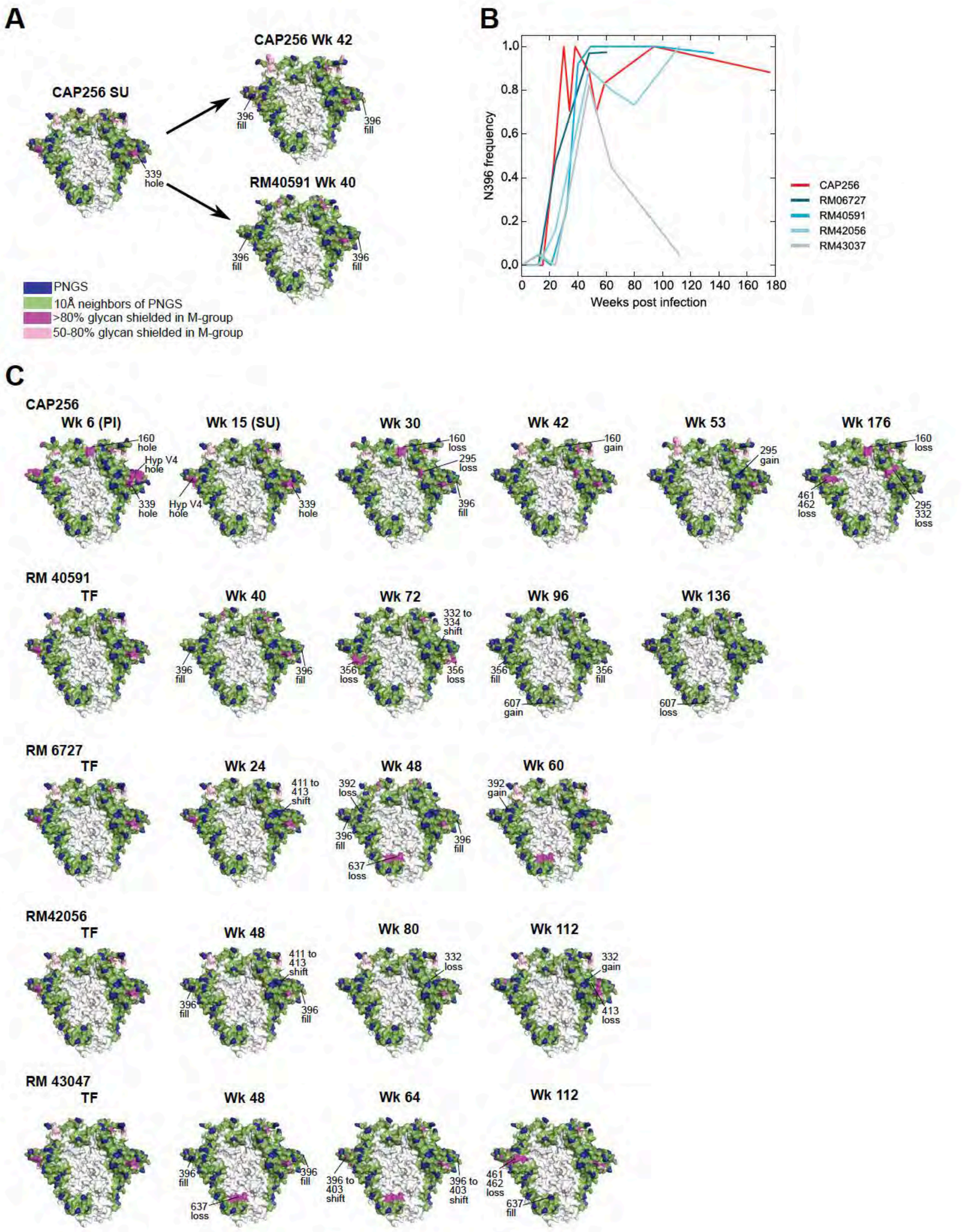

### Supplemental Figure S8

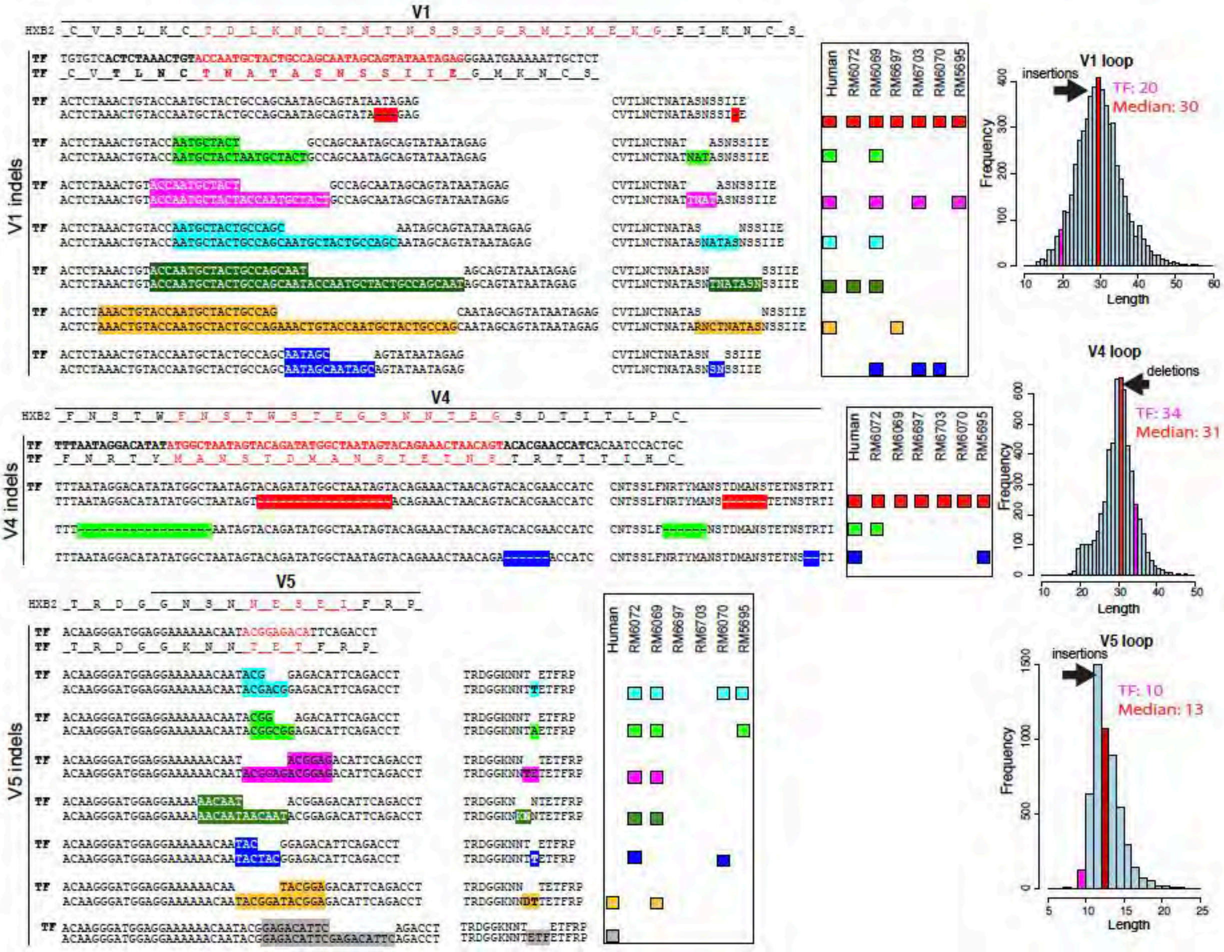

### Supplemental Figure S9

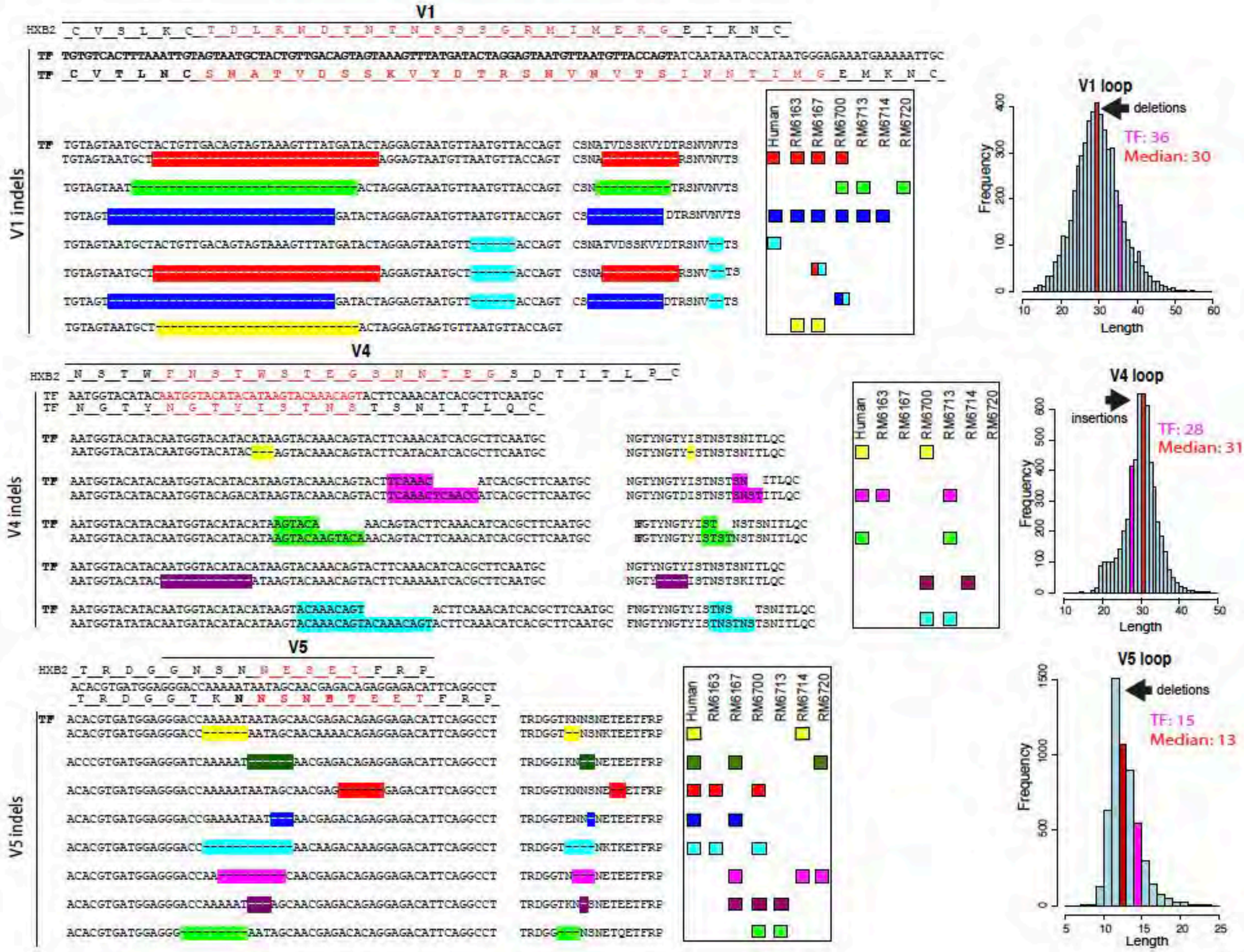

### Supplemental Figure S10

## A

| Virus | Mutation Location | RM6072 plasma w10 | RM6072 plasma w20 |
|---|---|---|---|
| TF_ID50 | | 286 | 2,397 |
| TF_RID50 ($ID50_{mut}/ID50_{TF}$) | | 1 | 1 |
| N130D | V1 | 0.3 | 0.6 |
| T234N | C2 | 0.3 | 1.2 |
| N279D | Loop-D | <0.1 | 0.7 |
| V281I | Loop-D | <0.1 | 0.7 |
| K302N | V3 | 224.3 | 51.9 |
| Y330H | V3 | 0.7 | 1.5 |
| N334S | V3C3 | 0.2 | 0.4 |
| H417R | V4 | 1.1 | 1.2 |
| K460E[a] | V5 | <0.1 | 1.8 |

>3 fold resistant >3 fold sensitive

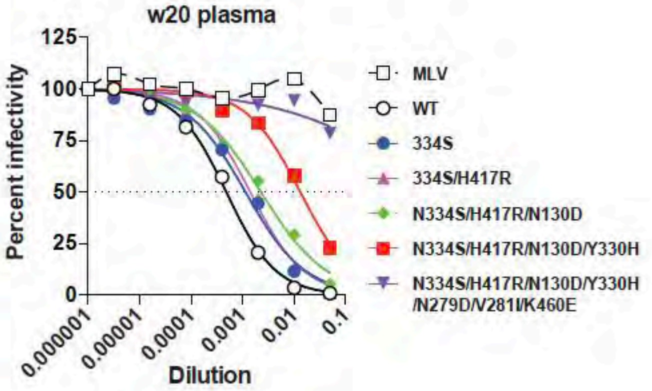

## B

| Virus | Mutation Location | RM6072 V3 mAbs DH647 | RM6072 V3 mAbs DH648 |
|---|---|---|---|
| TF_$IC_{50}$ (ug/ml) | | 2.2 | 1.0 |
| N130D | V1 | 3.3 | 0.9 |
| K302N | V3 | 15.5 | >20 |
| Y330H | V3 | 2.5 | 1.8 |
| N334S | V3C3 | >20 | >20 |
| H417R | V4 | 2.0 | 2.0 |

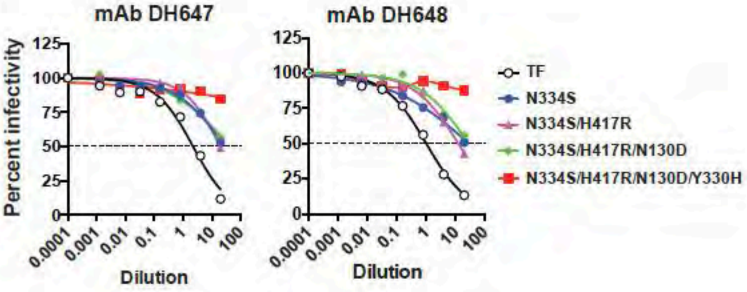

## C

| Virus | Mutation Location | RM6072 CD4bs mAbs DH650.UCA | RM6072 CD4bs mAbs DH650 | Human CH505 CD4bs mAbs CH235.UCA | Human CH505 CD4bs mAbs CH235.IA3 | Human CH505 CD4bs mAbs CH235.9 |
|---|---|---|---|---|---|---|
| TF_$IC_{50}$ (ug/ml) | | >20 | 2.2 | >20 | 3.6 | 0.4 |
| T234N | C2 | >20 | >20 | >20 | >20 | 0.6 |
| N279D | Loop-D | >20 | >20 | >20 | >20 | 0.4 |
| V281I | Loop-D | >20 | 3.5 | >20 | 5.6 | 0.2 |
| K460E | V5 | >20 | >20 | >20 | 13.2 | 1.3 |

### Supplemental Figure S11

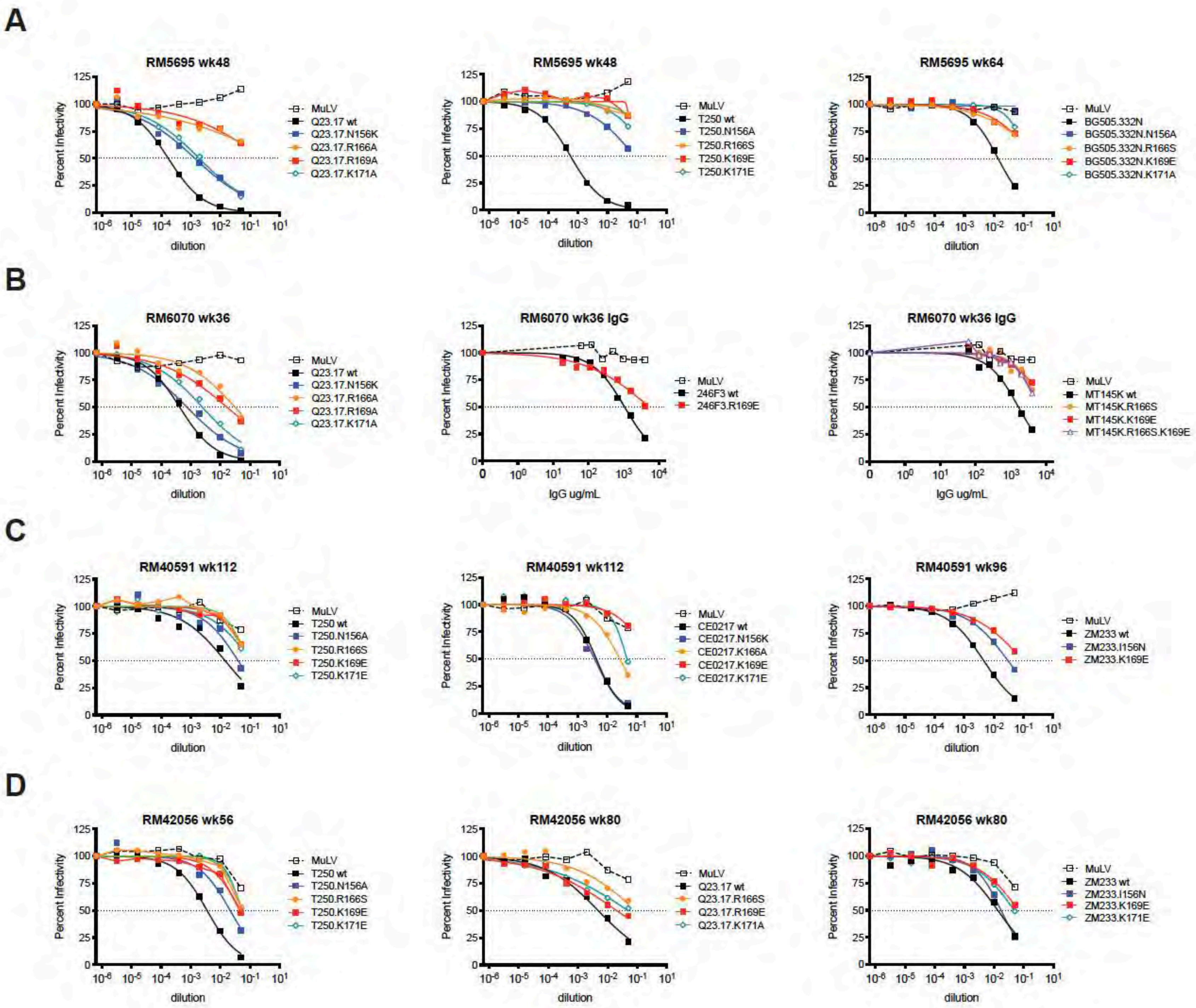

**Supplemental Figure S12**

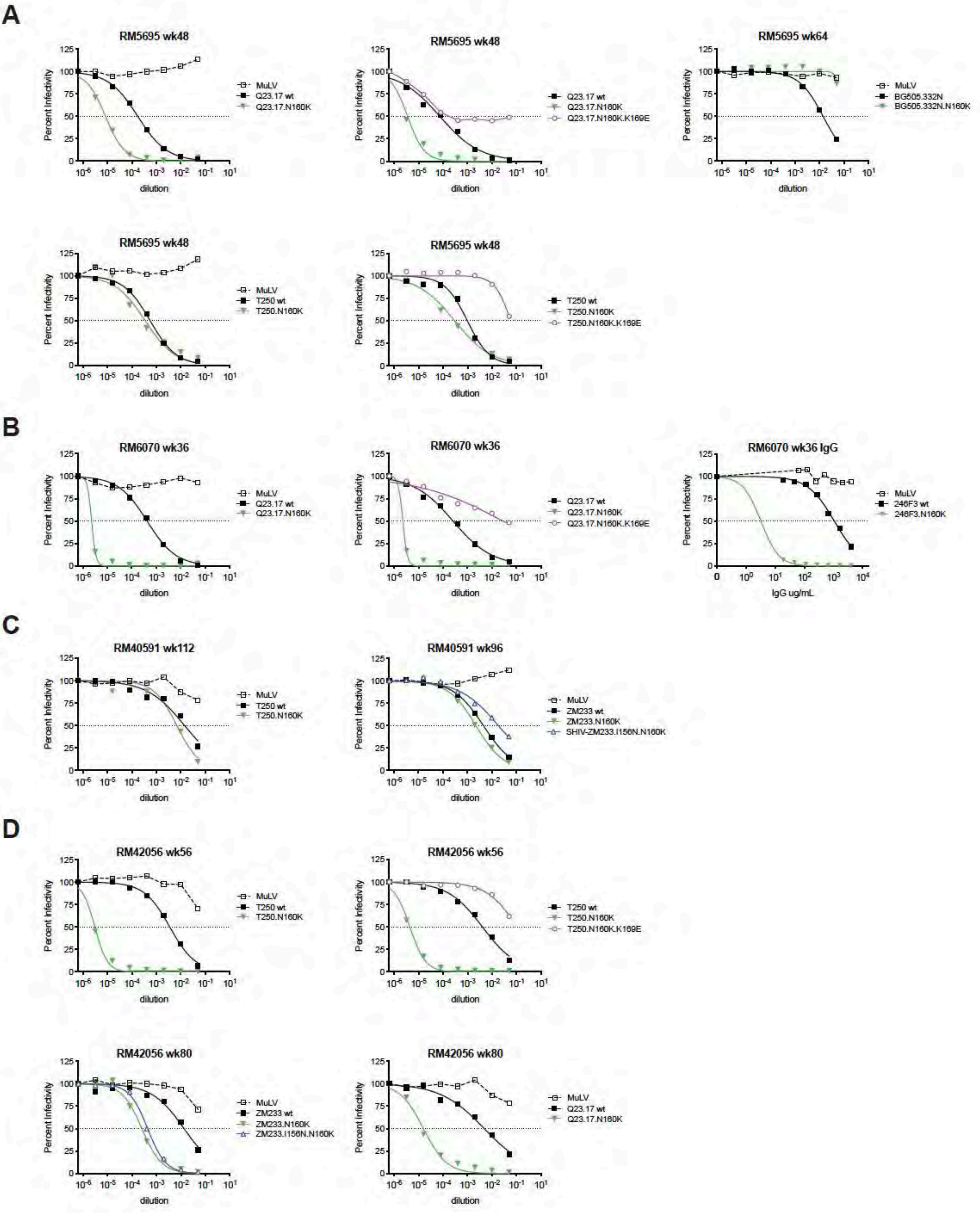

**Supplemental Figure S13**

**A**

**Autologous NAb responses to wildtype and V1 deleted variants of CH848 TF in SHIV CH848 infected RM plasma**

| Animal ID | Plasma timepoint | CH848 TF | CH848 TFdV1 (V1Δ10aa) | w36con (V1Δ10aa+ Mut) |
|---|---|---|---|---|
| RM6167 | w10 | 181 | 130 | <20 |
| | w36 | 435 | 948 | 28 |

**B**

**Epitope mapping of autologous, strain-specific neutralizing mAbs from RM6163**

| Animal ID | Antibody ID | CH848 TF | CH848 TFdV1 (Δ10aa) | CH848 N301D | CH848 N332D |
|---|---|---|---|---|---|
| RM6163 | DH898.1 | 2.84 | 11.49 | 3.19 | >50 |
| w52 LN | DH898.2 | 6.57 | 27.38 | 8.56 | >50 |
| | DH898.3 | 3.69 | 16.35 | 5.93 | >50 |
| | DH898.4 | 0.64 | 3.29 | 0.86 | >50 |
| | DH898.5 | 2.83 | 12.18 | 5.40 | >50 |
| | DH898.6 | 3.12 | 21.67 | 5.74 | >50 |

| IC50, *μ*g/mL |
|---|
| <1 |
| 1-10 |
| 10-50 |
| >50 |

**Supplemental Figure S14**

**A**

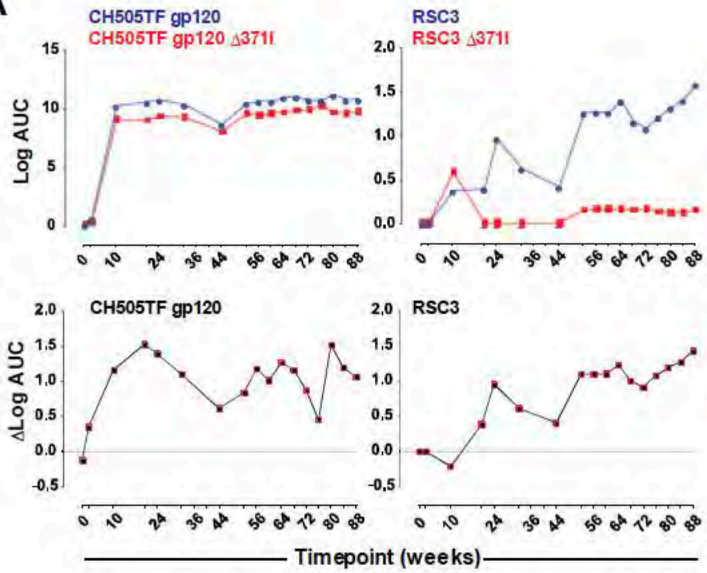

**B**

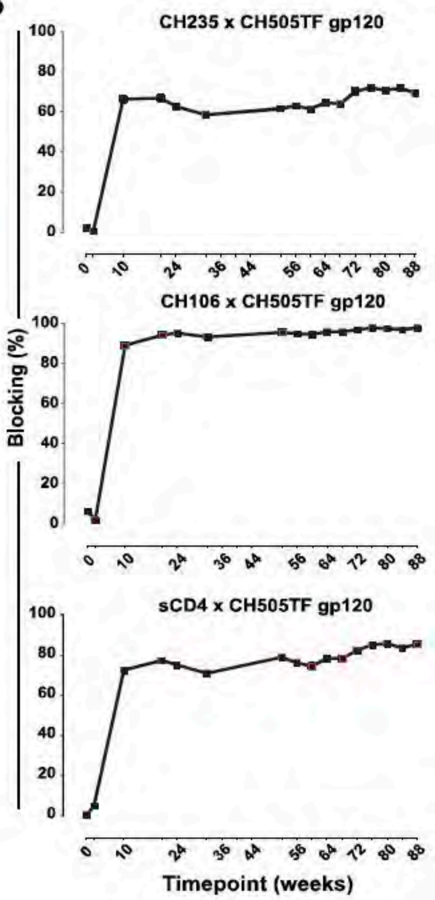

**C**

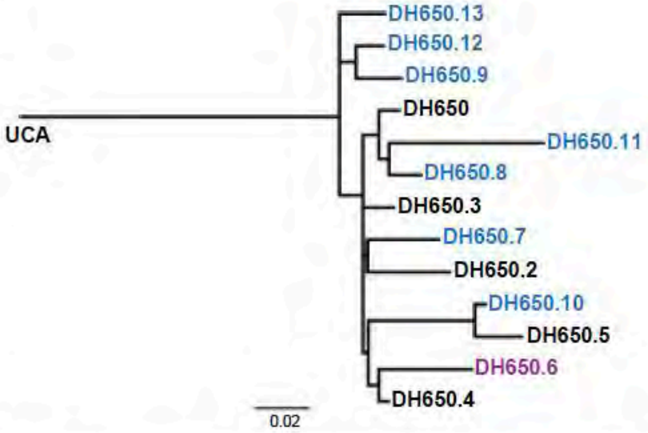

**D**

| mAb ID | Heavy chain (IGHV) | | | | Light chain (IGKV) | | |
|---|---|---|---|---|---|---|---|
| | IGHV | IGHJ | CDR3 length (aa) | Mut. Freq. (%, nn) | IGKV | CDR3 length (aa) | Mut. Freq. (%, nn) |
| DH650 | IGHV1-H | IGHJ4-1 | 14 | 4.9% | IGKV2-S17 | 9 | 3.2% |
| DH650.1 | IGHV1-H | IGHJ4-1 | 14 | 4.9% | *Unrecovered gene sequence* | | |
| DH650.2 | IGHV1-H | IGHJ4-1 | 14 | 6.2% | IGKV2-S17 | 9 | 4.3% |
| DH650.3 | IGHV1-H | IGHJ4-1 | 14 | 4.9% | IGKV2-S17 | 9 | 4.3% |
| DH650.4 | IGHV1-H | IGHJ4-1 | 14 | 4.5% | IGKV2-S17 | 9 | 2.8% |
| DH650.5 | IGHV1-H | IGHJ4-1 | 14 | 8.7% | IGKV2-S17 | 9 | 3.9% |
| DH650.6 | IGHV1-H | IGHJ4-1 | 14 | 4.9% | IGKV2-S17 | 9 | 5.3% |
| DH650.7 | IGHV1-H | IGHJ4-1 | 14 | 6.9% | IGKV2-S17 | 9 | 3.9% |
| DH650.8 | IGHV1-H | IGHJ4-1 | 14 | 4.9% | IGKV2-S17 | 9 | 4.6% |
| DH650.9 | IGHV1-H | IGHJ4-1 | 14 | 6.2% | IGKV2-S17 | 9 | 2.5% |
| DH650.10 | IGHV1-H | IGHJ4-1 | 14 | 8.3% | IGKV2-S17 | 9 | 3.2% |
| DH650.11 | IGHV1-H | IGHJ4-1 | 14 | 6.2% | IGKV2-S17 | 9 | 5.2% |
| DH650.12 | IGHV1-H | IGHJ4-1 | 14 | 5.6% | IGKV2-S17 | 9 | 3.5% |
| DH650.13 | IGHV1-H | IGHJ4-1 | 14 | 5.6% | IGKV2-S17 | 9 | 3.2% |
| DH650.14 | IGHV1-H | IGHJ4-1 | 14 | 8.0% | *Unrecovered gene sequence* | | |

**E**

| Antibody ID | CH505TF gp120 binding titer; EC50 (µg/ml) | |
|---|---|---|
| | WT | Δ371I |
| DH650 | 0.09 | 22.66 |
| DH650.3 | 1.94 | No binding |
| DH650.4 | 1.09 | No binding |
| DH650.5 | 2.02 | No binding |
| DH650.6 | No binding | No binding |
| DH650.7 | 0.22 | 8.72 |
| DH650.8 | 0.10 | 7.52 |
| DH650.9 | 0.16 | 24.05 |
| DH650.10 | 0.44 | No binding |
| CH235 | 0.05 | No binding |
| CH106 | 0.03 | No binding |

**F**

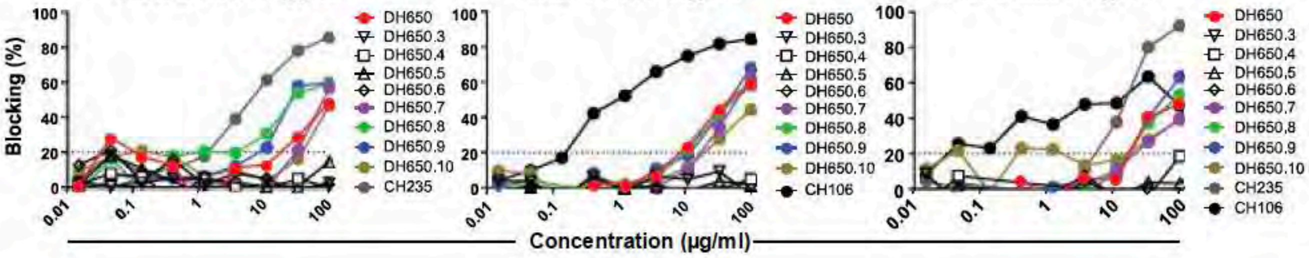

### Supplemental Figure S15

**A**

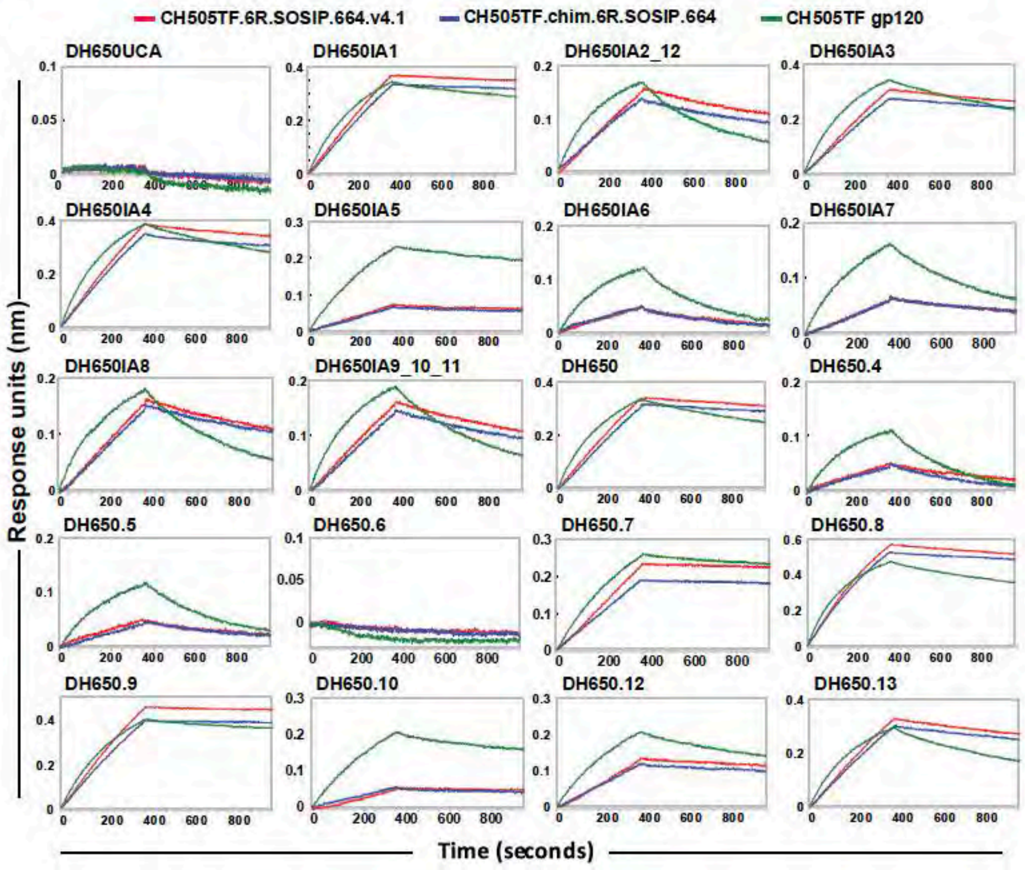

**B**

| IC50, µg/mL | >50 | 10-50 | 1-10 | 0.1-1 | <0.1 |
|---|---|---|---|---|---|

| Antibody ID | TF | TF gly4 | MuLV |
|---|---|---|---|
| DH650UCA | >50 | >50 | >50 |
| DH650IA1 | 3.8 | <0.02 | >50 |
| DH650IA2_12 | >50 | 0.06 | >50 |
| DH650IA3 | 4.2 | <0.02 | >50 |
| DH650IA4 | 4.6 | <0.02 | >50 |
| DH650IA5 | 11.4 | <0.02 | >50 |
| DH650IA6 | >50 | <0.02 | >50 |
| DH650IA7 | >50 | <0.02 | >50 |
| DH650IA8 | 20.0 | <0.02 | >50 |
| DH650IA9_10_11 | 20.0 | 0.03 | >50 |
| DH650 | 2.6 | <0.02 | >50 |
| DH650.2 | 10.0 | <0.02 | >50 |
| DH650.3 | >50 | 0.4 | >50 |
| DH650.4 | >50 | <0.02 | >50 |
| DH650.5 | >50 | 0.06 | >50 |
| DH650.6 | >50 | >50 | >50 |
| DH650.7 | 3.3 | <0.02 | >50 |
| DH650.8 | 2.0 | <0.02 | >50 |
| DH650.9 | 3.8 | <0.02 | >50 |
| DH650.10 | 10.3 | <0.02 | >50 |
| DH650.12 | 9.6 | <0.02 | >50 |
| DH650.13 | 8.3 | <0.02 | >50 |

**C**

| IC50, µg/mL | >50 | 10-50 | 1-10 | 0.1-1 | <0.1 |
|---|---|---|---|---|---|

| Antibody ID | HIV-1 isolate/ Tier phenotype/ IC50, µg/mL | | | | | | |
|---|---|---|---|---|---|---|---|
| | CH505 TF | JRFL | Q168 | Q842 | BG1168 | SF162 | MuLV |
| | 2 | 2 | 2 | 2 | 1 | 1 | n/a |
| DH650 | 9.6 | >50 | >50 | >50 | >50 | >50 | >50 |
| DH650.3 | >50 | >50 | >50 | >50 | >50 | >50 | >50 |
| DH650.4 | >50 | >50 | >50 | >50 | >50 | >50 | >50 |
| DH650.5 | >50 | >50 | >50 | >50 | >50 | >50 | >50 |
| DH650.6 | >50 | >50 | >50 | >50 | >50 | >50 | >50 |
| DH650.7 | 5.0 | >50 | >50 | >50 | >50 | >50 | >50 |
| DH650.8 | 4.5 | >50 | >50 | >50 | >50 | >50 | >50 |
| DH650.9 | 4.4 | >50 | >50 | >50 | >50 | >50 | >50 |
| DH650.10 | 23.7 | >50 | >50 | >50 | >50 | >50 | >50 |
| CH235 | 0.58 | 1.7 | >50 | 3.9 | >50 | >50 | >50 |

### Supplemental Figure S16

A

```
           51  HCDR2                    72
                   ********   *        *
CH235_UCA   INPSGGSTSYAQKFQGRVTMTR
CH235       .D..W.R.N...N....I....
8ANC131     .KR.-.RLM...N..D.LSLR.

DH650_UCA   INPYNGNTKYAQKFQGRVTMTR
DH650.13    ...RH.RIG.............
DH650.9     ...R..RIG.S.R.........
DH650.12    ...R..RIG.....KD......
DH650       ...R..RIG......D......
DH650.8     ...R..RIG......D......
DH650.11    ...RT.RIG...R..D......
DH650.7     ...R..RIG......D......
DH650.2     ...R..RIG.............
DH650.3     ...R..RIG.....KD......
DH650.10    .D.TY.R.G...R.KD.I....
DH650.5     .D.TH.R.G...R.KD.I....
DH650.4     ...R..RIG.....RD......
DH650.6     ...R..RIG..P..R.......
```

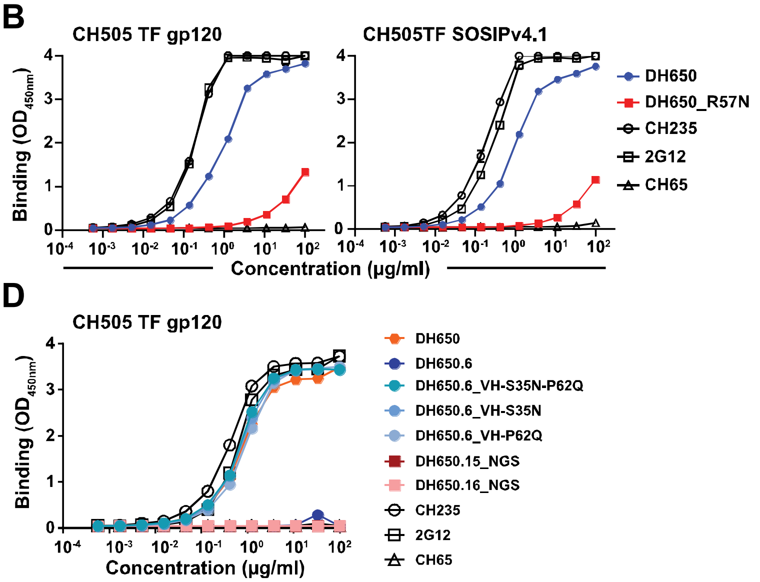

C

```
                                      HCDR1                     HCDR2                                                 HCDR3
DH650UCA      QVQLVQSGAEVKKPGSSVKVSCKASGYTFTDYYMHWVRQAPRQGLEWMGWINPYNGNTKYAQKFQGRVTMTRDTSTSTAYMELSSLRSEDTAVYYCARGRLELTASHFDYWGQGVLVTVSS
DH650.13      ...............A...L.......A..L.SIN......G...........RH.RIG..................I.....G..........F..............V.....P.....
DH650.9       ...............A...L.......A..S.SIN......G...........R..RIG.S.R..............I.....G..........F...........Q..K.....P.....
DH650.12      ...............A...L.......A..I.SIN......G...........R..RIG.....KD...........I.....G..........F...........Q..K.....P.....
DH650         ...............A...L.......G..I.SIN......G...........R..RIG......D...........I.....G...Y......F.T..Q......R..K.....P.....
DH650.8       ...............A...L.......A..I.SIN......G...........R..RIG......D..........NI.....G..........F....Q......R..K.....P.....
DH650.11      ...............D..RL..R....A..I.SIN......G...........RT.RIG...R..D..........NI.....G..........F....Q......R..K.....P.....
DH650.7       .........A.....A...L.......A..L.SIN......G...........R..RIG......D.........M.I....VG..........F..............T.....P.....
DH650.2       ......F........A...L.......A..L.SIN......G...........R..RIG..................IG....G..........F...........Q..K.....P.....
DH650.3       ...............A...L...G...A..I.SIN......G...........R..RIG.....KD...........I.....G..........F..............N.....P.....
DH650.10      ....L..........A...L.......A..S.SIN......G...Q...Y.D.TY.R.G...R.KD.I.........M.....GR.........F..............N.....P.....
DH650.5       ....L..........A...L.......A..S.SIN......G...Q...Y.D.TH.R.G...R.KD.I........NM.....GR.........F..............N.....P.....
DH650.4       ...............A...L.......A..I.SIN......G...........R..RIG.....RD...........I.....G..........F.T............N.....P.....
DH650.6       ...............A...L..R....A..I.SIS......G...........R..RIG..P..R............I.....G..........F.T..Q......R..T..R..P.....
DH650.15_NGS  ...............A...L.......N.IS.TIN......G...........K...IG........................G..........F.T..Q.D....R..T.....P.....
DH650.16_NGS  ...............A...L...G......S.SVN......G..P........N....D........L...............G..........F.T..Q.D.I..R..T.....P.....
DH650.17_NGS  ...............A...........P..P.KIN......G...........A..Q.G.........V.S.....T......G..........F.T..Q.D....R..T.....P.....
DH650.18_NGS  ...............A.......T..FI.GN.GIS......G.........D.R..RIG..P..R..................G..........F.T..Q.D....R..T.....P.....
DH650.19_NGS  ...............A..RL..........IHSIN......G.........D.R..RIG..P..R..........R......GG.........F.T..Q.D....R..T.....P.....
DH650.20_NGS  ...............A..RL..........IHSIN......G.........D.R..RIG..P..R..................G..........F.T..Q.D....R..T.....P.....
DH650.21_NGS  E.R.I.......M..A..RL..........IHSIN......G.........D.R..RIG..P..R..................G..........F.T..Q.D....R..T.....P.....
DH650.22_NGS  ...............A..RL..........IHSIN......G.........D.R..RIG..P..R..................G..........F.T..Q.D....R..T.....P.....
DH650.23_NGS  ...............A..RL..........IHSIN......G.........D.R..RIG..P..R..................G..........F.T..Q.D....R..T.....P.....
DH650.24_NGS  .........A.....A..RL..........IHSIN......G.........D.R..RIG..P..R..................G..........F.T..Q.D....R..T.....P.....
DH650.25_NGS  ...............A..RL..........IHSIN......G.........D.R..RIG..P..R..................G..........F.T..Q.D....R..T.....P.....
DH650.26_NGS  ...............A...L..........IHSIN......G.........D.R..RIG..P..R..................G..........F.T..Q.D....R..T.....P.....
DH650.27_NGS  ...............A..RL..........IHSIN......G.........D.R..RIG..P..R..................G..........F.T..Q.D....R..T.....P.....
DH650.28_NGS  ...............A..RL..........IHSIN......G.........D.R..RIGD.P..R..................G..........F.T..E.D....R..T.....P.....
DH650.29_NGS  ...............A..RL..........IHSIN......G.........V.S..RIG..P..R..................G..........F.T.CQ.D....R..T..H..P.....
DH650.30_NGS  ...............A..RL..........IHSIN......G.........D.R..RIG..P..R..................G..........F.T..Q.D....R..T.....P.....
DH650.31_NGS  ...............A..RL..........IHSIN......G.........D.R..RIG..P..R..................G..........F.T..Q.D....R..T.....P.....
DH650.32_NGS  ...............A..RL..........IHSIN......G.........D.R..RIG..P..R.............................F.T..Q.D....R..T.....P.....
DH650.33_NGS  ..R............A..RL..........IHSIN......G.........D.R..RIG..P..R..................G..........F.T..Q.D....R..T.....P.....
DH650.34_NGS  ...............A..RL..........IHSIN......G.........D.R..RIG.VP..R..................G..........F.T..Q.D....R..T.....P.....
```

E

```
                                        LCDR1                        LCDR2                                LCDR3
CH235UCA_VK   EIVLTQSPATLSVSPGERATLSCRASQSV------SSNLAWYQQKPGQAPRLLIYGASTRATGIPARFSGSGSGTEFTLTISSLQSEDFAVYYCQQYNNWWT-FGQGTKVEIK
CH235_VK      .............................------R........R...........T......V......R........A...M........L.L.......-..........

DH650UCA_VK   DIVMTQTPLSLPVTPGEPASISCRSSQSLLDSEDGNTYLDWYLQKPGQSPQLLIYEVSNRASGVPDRFSGSGSDTDFTLKISRVEAEDVGVYYCMQALEFPYSFGQGTKVEIK
DH650_VK      ......S............A...........RA...............................................................G................
DH650.2_VK    N.....S............A...........RA...............P........................H....................G................
DH650.3_VK    ......S............A...........RA...............................................................G................
DH650.4_VK    ......S............A...........RD...............................................................G.............Q..
DH650.5_VK    ......S............A...........RD..D...............................................IL..D.......L.G.....R..........
DH650.6_VK    ......S............A...........RS.........R...........F...................................I.....G.....R..........
DH650.7_VK    ......S............A...........RA.E.............................................................G..Y.............
DH650.8_VK    ......S............A...........RA.........................S....................R............F...G.A..............
DH650.9_VK    ...................A...........RA...............................................................G..Y.............
DH650.10_VK   .....HS.H..........A...........RD...............................................................G................
DH650.11_VK   ...................A...........RD..............PA.A.SVL...K.....................................G.G...T..........
DH650.12_VK   ......S............A...........RS...............................................................G..Y.............
DH650.13_VK   ...................A...........RA.........................S.....................................G................
```

### Supplemental Figure S17

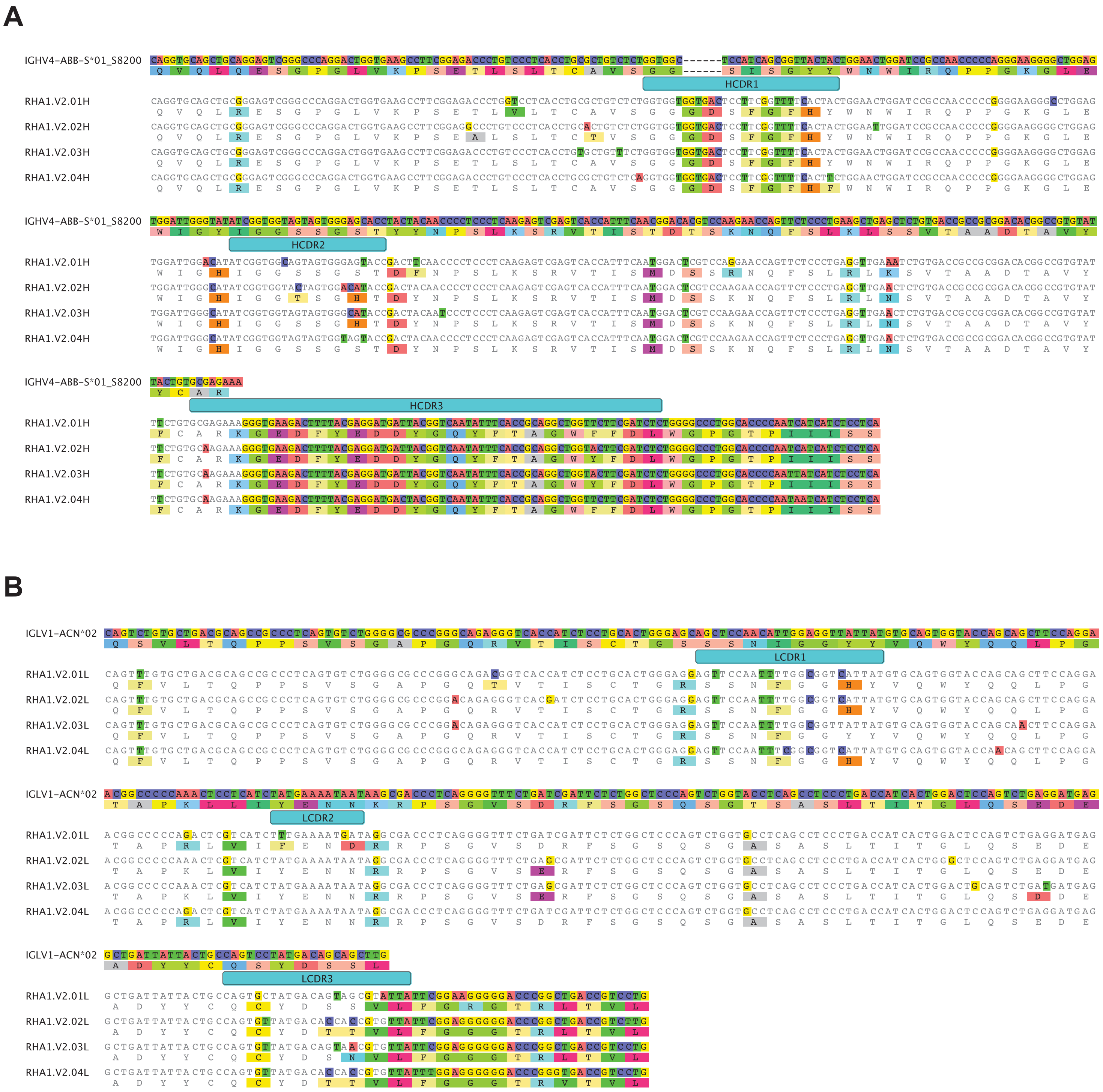

**Supplemental Figure S18**

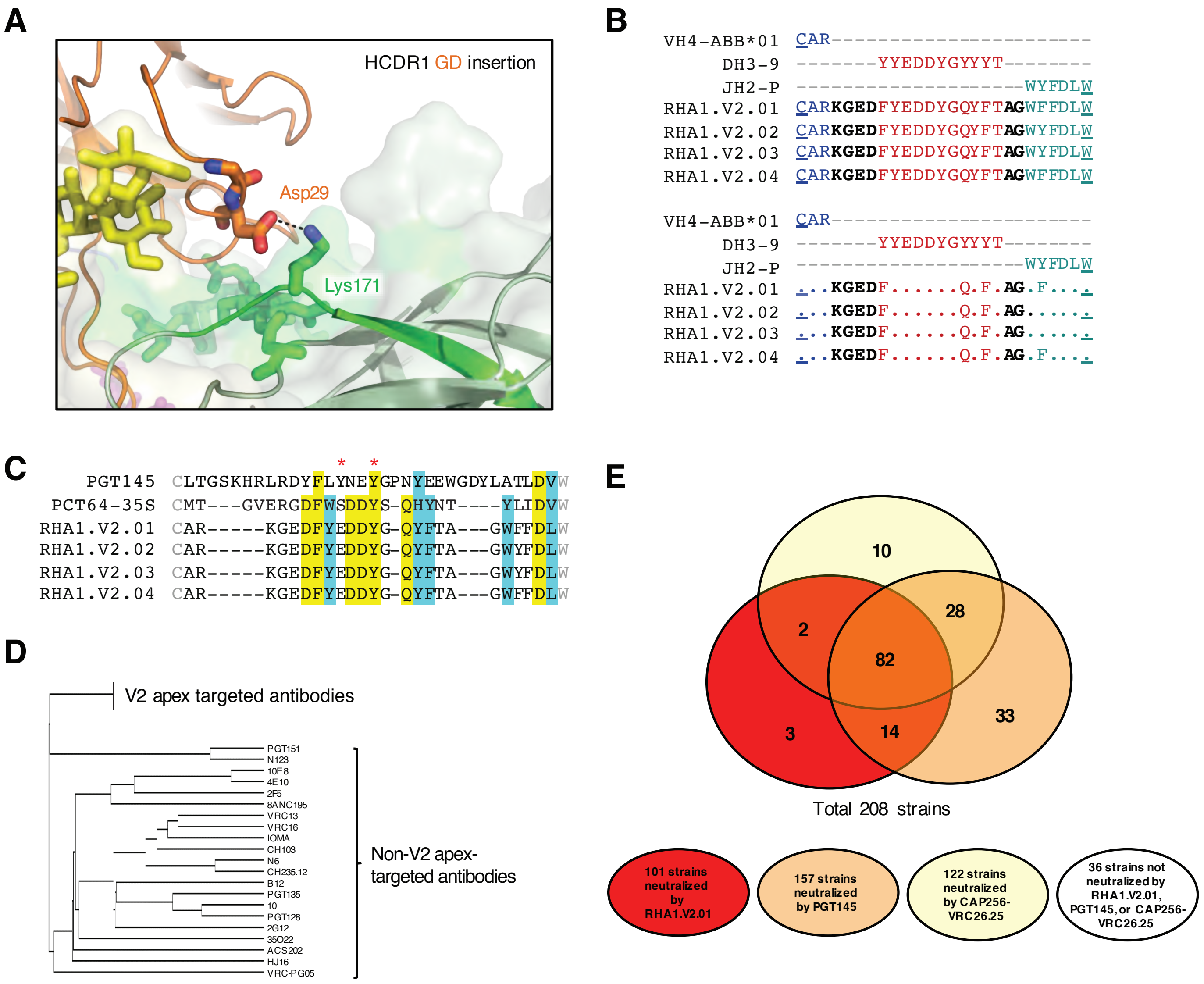

**Supplemental Figure S19**

A

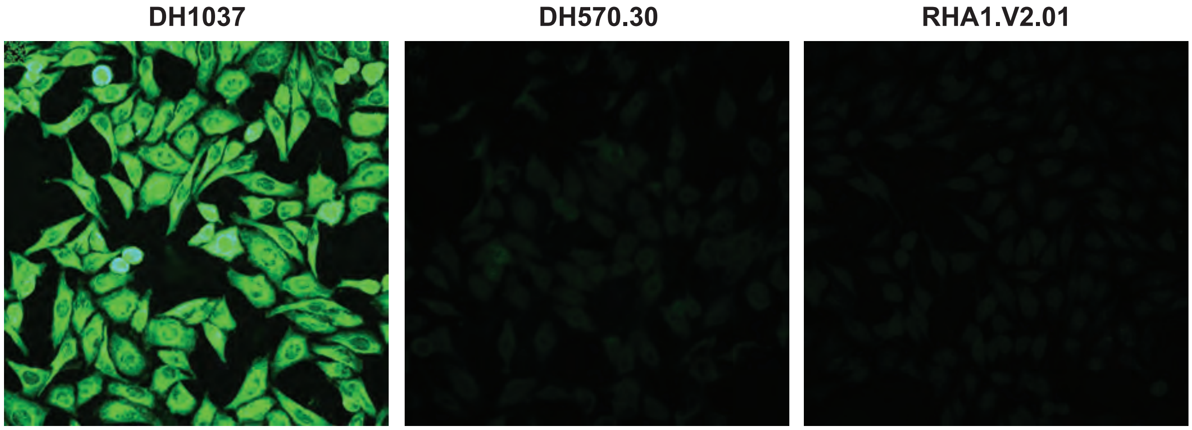

B

Autoantigens

| | SSA | SSB | Sm | RNP | Scl 70 | Jo 1 | dsDNA | Cent B | Histone |
|---|---|---|---|---|---|---|---|---|---|
| Positive control 1 | - | - | - | - | - | - | - | + | - |
| Positive control 2 | + | + | - | - | - | + | + | - | + |
| Positive control 3 | - | - | + | + | + | - | - | - | - |
| CH65 | - | - | - | - | - | - | - | - | - |
| 4E10 | - | + | + | - | - | + | - | + | + |
| RHA1.V2.01 | - | - | - | - | - | - | - | - | - |

Supplemental Figure S20

| | | Rhesus mAbs | | | | |
|---|---|---|---|---|---|---|
| | | RHA1.V2.01 | RHA1.V2.02 | RHA1.V2.03 | RHA1.V2.04 | RM5695 wk56 plasma |
| Global panel | 246F3 | 0.21 | 0.24 | 0.31 | 0.24 | 109 |
| | X1632 | 0.16 | 0.12 | 0.32 | 0.24 | 465 |
| | 25710 | 0.74 | 0.33 | 2.18 | 1.76 | 28 |
| | CNE55 | 0.35 | 0.63 | 2.75 | 1.45 | 20 |
| | X2278 | 0.40 | 0.41 | 1.87 | 1.18 | 28 |
| | BJOX0020000 | 1.94 | 0.64 | 1.99 | 2.09 | 22 |
| | CNE8 | 4.09 | 1.27 | 6.78 | 5.76 | 179 |
| | CE0217 | 3.085 | 1.77 | >50 | >50 | 26 |
| | TRO.11 | >50 | >50 | >50 | >50 | 72 |
| | CE1176.A3 | >50 | >50 | >50 | >50 | 20 |
| | CH119.10 | >50 | 31 | >50 | >50 | <20 |
| | BG505.T332N | 0.05 | 0.11 | 0.11 | 0.10 | 139 |
| | CAP256SU | 0.45 | 0.27 | 0.85 | 0.76 | 65 |
| | ZM233.6 | 0.01 | 0.43 | 0.02 | 0.01 | 413 |
| | WITO.33 | 0.03 | 0.04 | 0.04 | 0.03 | 138 |
| | T250-4 | 0.01 | 0.01 | 0.02 | 0.01 | 1977 |
| | Q23.17 | 0.005 | 0.005 | 0.006 | 0.003 | 5051 |
| | MT145K | 0.13 | 0.07 | 0.181 | 0.15 | 161 |
| autologous | CH505 TF | 0.463 | 0.180 | 0.648 | 0.507 | 421 |

IC50 ug/mL
- 10-50
- 1.0-10
- 0.10-1.0
- <0.10

Plasma ID50
- 20-100
- 100-500
- 500-5000

**Supplemental Figure S21**

#### RHA1.V2.01

IC50 (ug/mL)

| Legend |
|---|
| .001-.01 |
| .01-.100 |
| .100-1.00 |
| 1.00-10.0 |
| >10.0 |

| Virus ID | Clade | |
|---|---|---|
| 0260.v5.c36 | A | 13.9 |
| 0330.v4.c3 | A | 0.046 |
| 0439.v5.c1 | A | >50 |
| 3365.v2.c20 | A | 0.390 |
| 3718.v3.c11 | A | 0.809 |
| 398-F1.F6.20 | A | >50 |
| BB201.B42 | A | 0.010 |
| BB539.2B13 | A | >50 |
| BG505.W6M.C2 | A | 0.067 |
| BI369.9A | A | 0.083 |
| BS208.B1 | A | 0.045 |
| KER2008.12 | A | 0.043 |
| KER2018.11 | A | 0.013 |
| KNH1209.18 | A | 0.958 |
| MB201.A1 | A | 14.7 |
| MB539.2B7 | A | 1.36 |
| MI369.A5 | A | 0.682 |
| MS208.A1 | A | >50 |
| Q23.17 | A | 0.005 |
| Q259.17 | A | 0.397 |
| Q769.d22 | A | >50 |
| Q769.h5 | A | >50 |
| Q842.d12 | A | 0.020 |
| QH209.14M.A2 | A | >50 |
| RW020.2 | A | >50 |
| UG037.8 | A | >50 |
| 246-F3.C10.2 | AC | 0.179 |
| 3301.v1.c24 | AC | >50 |
| 3589.v1.c4 | AC | >50 |
| 6540.v4.c1 | AC | 0.199 |
| 6545.v4.c1 | AC | 0.120 |
| 0815.v3.c3 | ACD | >50 |
| 6095.v1.c10 | ACD | 12.5 |
| 3468.v1.c12 | AD | >50 |
| Q168.a2 | AD | 0.069 |
| Q461.e2 | AD | >50 |
| 620345.c1 | AE | 23.5 |
| BJOX009000.02.4 | AE | >50 |
| BJOX010000.06.2 | AE | >50 |
| BJOX025000.01.1 | AE | >50 |
| BJOX028000.10.3 | AE | >50 |
| C1080.c3 | AE | 0.001 |
| C2101.c1 | AE | >50 |
| C3347.c11 | AE | >50 |
| C4118.09 | AE | 0.049 |
| CM244.ec1 | AE | 0.004 |
| CNE3 | AE | 0.041 |
| CNE5 | AE | 0.017 |
| CNE55 | AE | 1.17 |
| CNE56 | AE | >50 |
| CNE59 | AE | 0.102 |
| CNE8 | AE | 8.55 |
| M02138 | AE | >50 |
| R1166.c1 | AE | >50 |
| R2184.c4 | AE | >50 |
| R3265.c6 | AE | >50 |
| TH023.6 | AE | >50 |
| TH966.8 | AE | 0.092 |
| TH976.17 | AE | >50 |
| 235-47 | AG | 0.251 |
| 242-14 | AG | 0.869 |
| 263-8 | AG | >50 |
| 269-12 | AG | >50 |
| 271-11 | AG | 0.018 |
| 928-28 | AG | >50 |
| DJ263.8 | AG | 1.06 |
| T250-4 | AG | 0.010 |
| T251-18 | AG | >50 |
| T253-11 | AG | >50 |
| T255-34 | AG | 0.077 |
| T257-31 | AG | 30.3 |
| T266-60 | AG | 13.5 |
| T278-50 | AG | 9.27 |
| T280-5 | AG | >50 |
| T33-7 | AG | 0.011 |

| Virus ID | Clade | |
|---|---|---|
| 3988.25 | B | 11.5 |
| 5768.04 | B | 0.985 |
| 6101.10 | B | >50 |
| 6535.3 | B | 4.16 |
| 45_01dG5 | B | 3.97 |
| 89.6.DG | B | >50 |
| AC10.29 | B | 1.17 |
| ADA.DG | B | 8.96 |
| Bal.01 | B | >50 |
| BaL.26 | B | >50 |
| BG1168.01 | B | >50 |
| BL01.DG | B | >50 |
| BR07.DG | B | >50 |
| BX08.16 | B | 0.073 |
| CAAN.A2 | B | >50 |
| CNE10 | B | >50 |
| CNE12 | B | >50 |
| CNE14 | B | >50 |
| CNE4 | B | >50 |
| CNE57 | B | >50 |
| HO86.8 | B | 43.6 |
| HT593.1 | B | 6.70 |
| HXB2.DG | B | >50 |
| JRCSF.JB | B | 0.015 |
| JRFL.JB | B | >50 |
| MN.3 | B | >50 |
| PVO.04 | B | >50 |
| QH0515.01 | B | >50 |
| QH0692.42 | B | >50 |
| REJO.67 | B | 1.06 |
| RHPA.7 | B | >50 |
| SC422.8 | B | >50 |
| SF162.LS | B | >50 |
| SS1196.01 | B | >50 |
| THRO.18 | B | >50 |
| TRJO.58 | B | >50 |
| TRO.11 | B | >50 |
| WITO.33 | B | 0.019 |
| X2278.C2.B6 | B | 0.133 |
| YU2.DG | B | >50 |
| BJOX002000.03.2 | BC | 0.821 |
| CH038.12 | BC | 1.97 |
| CH070.1 | BC | 0.052 |
| CH117.4 | BC | 0.005 |
| CH119.10 | BC | 21.1 |
| CH181.12 | BC | 0.009 |
| CNE15 | BC | >50 |
| CNE19 | BC | >50 |
| CNE20 | BC | 7.47 |
| CNE21 | BC | 0.069 |
| CNE40 | BC | >50 |
| CNE7 | BC | >50 |

| Virus ID | Clade | |
|---|---|---|
| 286.36 | C | 0.701 |
| 288.38 | C | >50 |
| 0013095-2.11 | C | >50 |
| 001428-2.42 | C | 0.057 |
| 00836-2.5 | C | >50 |
| 0921.v2.c14 | C | >50 |
| 16055-2.3 | C | 0.077 |
| 16845-2.22 | C | >50 |
| 16936-2.21 | C | >50 |
| 25710-2.43 | C | 0.856 |
| 25711-2.4 | C | >50 |
| 25925-2.22 | C | 0.156 |
| 26191-2.48 | C | 0.404 |
| 3168.v4.c10 | C | >50 |
| 3637.v5.c3 | C | >50 |
| 3873.v1.c24 | C | 6.10 |
| 426c | C | >50 |
| 6322.v4.c1 | C | >50 |
| 6471.v1.c16 | C | >50 |
| 6631.v3.c10 | C | >50 |
| 6644.v2.c33 | C | >50 |
| 6785.v5.c14 | C | 0.107 |
| 6838.v1.c35 | C | 0.271 |
| 96ZM651.02 | C | >50 |
| BR025.9 | C | 0.051 |
| CAP210.E8 | C | 11.2 |
| CAP244.D3 | C | >50 |
| CAP256.206.C9 | C | 0.225 |
| CAP45.G3 | C | 0.006 |
| Ce1176.A3 | C | >50 |
| CE703010217.B6 | C | 2.19 |
| CNE30 | C | >50 |
| CNE31 | C | >50 |
| CNE53 | C | 6.18 |
| CNE58 | C | 0.045 |
| DU123.06 | C | 0.281 |
| DU151.02 | C | 1.05 |
| DU156.12 | C | 0.350 |
| DU172.17 | C | >50 |
| DU422.01 | C | >50 |
| MW965.26 | C | 12.6 |
| SO18.18 | C | 0.205 |
| TV1.29 | C | 2.28 |
| TZA125.17 | C | 40.8 |
| TZBD.02 | C | 4.01 |
| ZA012.29 | C | >50 |
| ZM106.9 | C | 24.7 |
| ZM109.4 | C | >50 |
| ZM135.10a | C | >50 |
| ZM176.66 | C | 0.078 |
| ZM197.7 | C | 43.1 |
| ZM214.15 | C | >50 |
| ZM215.8 | C | >50 |
| ZM233.6 | C | 0.009 |
| ZM249.1 | C | 0.324 |
| ZM53.12 | C | 0.589 |
| ZM55.28a | C | >50 |
| 3326.v4.c3 | CD | 0.162 |
| 3337.v2.c6 | CD | >50 |
| 3817.v2.c59 | CD | >50 |
| 191821.E6.1 | D | 48.6 |
| 231965.c01 | D | >50 |
| 247-23 | D | 0.211 |
| 3016.v5.c45 | D | 0.572 |
| 57128.vrc15 | D | >50 |
| 6405.v4.c34 | D | >50 |
| A03349M1.vrc4a | D | >50 |
| A07412M1.vrc12 | D | 9.22 |
| NKU3006.ec1 | D | >50 |
| UG021.16 | D | >50 |
| UG024.2 | D | >50 |
| P0402.c2.11 | G | 2.14 |
| P1981.C5.3 | G | >50 |
| X1193.c1 | G | >50 |
| X1254.c3 | G | >50 |
| X1632.S2.B10 | G | 0.112 |
| X2088.c9 | G | >50 |
| X2131.C1.B5 | G | 5.90 |
| SIVmac251.30.SG3 | NA | >50 |
| SVA.MLV | NA | >50 |

### Supplemental Figure S22

| | RHA1.V2.01 | RHA1.V2.03 | RHA1.V2.04 |
|---|---|---|---|
| **T250-4 wt** | 0.012 | 0.019 | 0.017 |
| T250-4.K169E | >25 | >25 | >25 |
| T250-4.R166S | >25 | >25 | >25 |
| T250-4.N160K | >25 | >25 | >25 |
| **246-F3 wt** | 0.352 | 0.413 | 0.243 |
| 246-F3.R169E | >25 | >25 | >25 |
| 246-F3.N160K | >25 | >25 | >25 |
| **BG505.332N wt** | 0.164 | 0.268 | 0.278 |
| BG505.332N.K169E | >25 | >25 | >25 |
| BG505.332N.R166S | >25 | >25 | >25 |
| BG505.332N.N160K | >25 | >25 | >25 |
| **CAP256SU wt** | 0.412 | 0.791 | 0.428 |
| CAP256SU.K169E | >25 | >25 | >25 |
| CAP256SU.R166S | >25 | >25 | >25 |
| CAP256SU.T162I | >25 | >25 | >25 |
| **X1632 wt** | 0.433 | 0.284 | 0.118 |
| X1632.K169E | >25 | >25 | >25 |
| X1632.N160K | >25 | >25 | >25 |
| **MT145K wt** | 0.16 | 0.25 | 0.23 |
| MT145K.K169E | >25 | >25 | >25 |
| MT145K.R166S | >25 | >25 | >25 |
| MT145K.N160D | >25 | >25 | >25 |
| **Q23.17 wt** | 0.007 | 0.009 | 0.008 |
| Q23.17.R169E | >25 | >25 | >25 |
| Q23.17.R166S | >25 | >25 | >25 |
| Q23.17.N160K | 2.96 | 1.37 | 1.00 |

| <0.10 | 0.10-1.0 | 1.0-10 | 10-50 |
|---|---|---|---|

IC50 ug/mL

**Supplemental Figure S23**

**A**

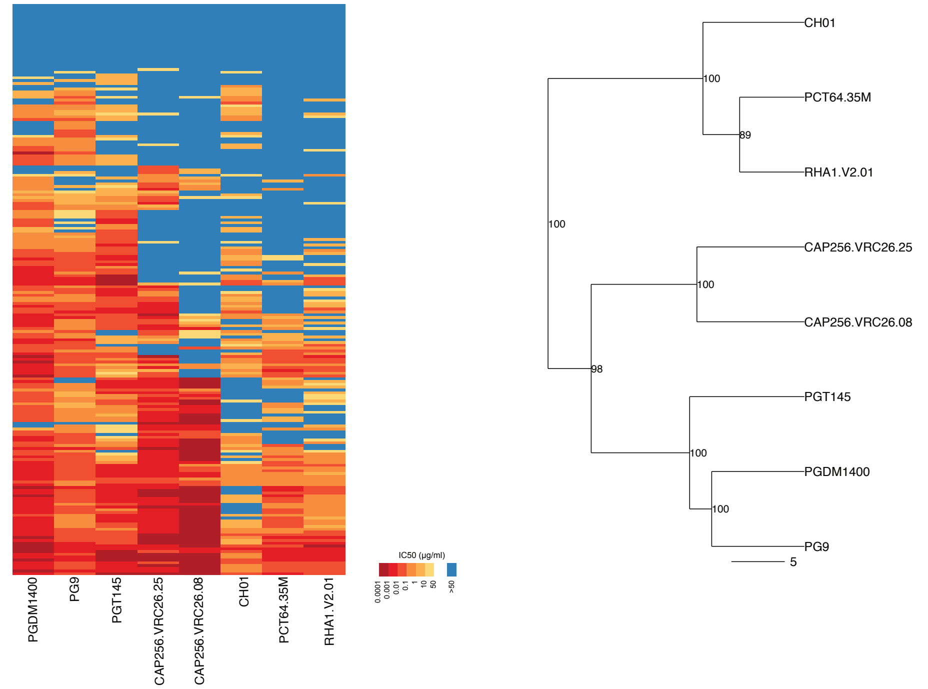

**B**

| | RHA1 sensitive | RHA1 resistant |
|---|---|---|
| **PCT64-35M sensitive** | 72 | 17 |
| **PCT6-35M resistant** | 29 | 90 |
| p=2.0x10^-16; Odd's ratio (OR) =13.14; Accuracy (Acc) = 0.78 | | |

| | RHA1 sensitive | RHA1 resistant |
|---|---|---|
| **CAP256.25 sensitive** | 84 | 38 |
| **CAP256.25 resistant** | 17 | 69 |
| p=1.7x10^-12; OR=57.35; Acc = 0.74 | | |

| | RHA1 sensitive | RHA1 resistant |
|---|---|---|
| **PG9 sensitive** | 100 | 68 |
| **PG9 resistant** | 1 | 39 |
| p=2.3x10^-12; OR=57.35; Acc = 0.69 | | |

| | RHA1 sensitive | RHA1 resistant |
|---|---|---|
| **PGDM1400 sensitive** | 99 | 68 |
| **PGDM1400 resistant** | 2 | 39 |
| p=2.8x10^-11; OR=28.39; Acc = 0.66 | | |

| | RHA1 sensitive | RHA1 resistant |
|---|---|---|
| **PGT145 sensitive** | 96 | 61 |
| **PGT145 resistant** | 5 | 46 |
| p=4.4x10^-11; OR=14.47; Acc=0.68 | | |

| | RHA1 sensitive | RHA1 resistant |
|---|---|---|
| **CAP256.08 sensitive** | 67 | 28 |
| **CAP256.08 resistant** | 34 | 79 |
| p=7.3x10^-9; OR=5.56; Acc=0.70 | | |

| | RHA1 sensitive | RHA1 resistant |
|---|---|---|
| **CH01 sensitive** | 72 | 38 |
| **CH01 resistant** | 29 | 69 |
| p=2.4x10^-7; OR=4.51; Acc=0.67 | | |

**C**

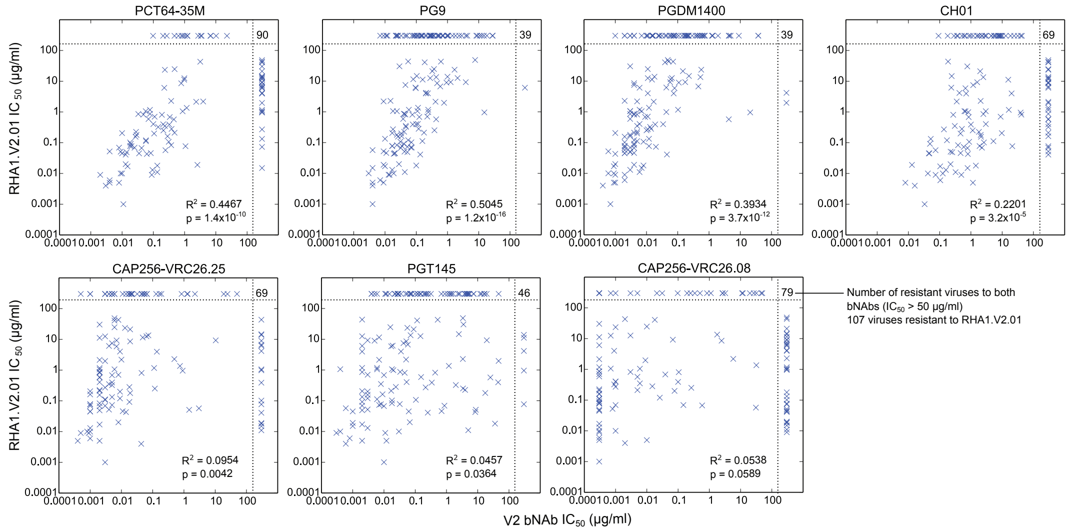

### Supplemental Figure S24

#### Subtype A

| Virus ID | Clade | RHA1. V2.01 | PCT64-35M | CAP256-VRC26.08 | CAP256-VRC26.25 | CH01 | PG9 | PGT145 | PGDM 1400 |
|---|---|---|---|---|---|---|---|---|---|
| Q23.17 | A | 0.005 | 0.004 | 0.010 | 0.002 | 0.008 | 0.009 | 0.002 | 0.0009 |
| BB201.B42 | A | 0.010 | 0.010 | 0.001 | 0.001 | 0.126 | 0.019 | 0.030 | 0.0009 |
| KER2018.11 | A | 0.013 | 0.094 | 0.0003 | 0.001 | 0.159 | 0.017 | 0.003 | 0.003 |
| Q842.d12 | A | 0.020 | 0.013 | | 0.008 | 0.223 | 0.028 | 0.067 | 0.001 |
| KER2008.12 | A | 0.043 | 0.062 | | 0.014 | 0.802 | 0.019 | 8.33 | 0.001 |
| BS208.B1 | A | 0.045 | 0.010 | 0.0003 | 0.001 | 0.045 | 0.021 | 0.001 | 0.002 |
| 0330.v4.c3 | A | 0.046 | 0.083 | | 0.011 | 0.468 | 0.011 | 0.697 | 0.004 |
| BG505.W6M.C2 | A | 0.067 | 0.042 | 0.589 | 0.002 | 0.777 | 0.032 | 0.010 | 0.004 |
| BI369.9A | A | 0.083 | 0.373 | 0.0003 | 0.002 | 0.387 | 0.024 | 0.018 | 0.004 |
| 3365.v2.c20 | A | 0.390 | 0.324 | 0.100 | | 1.00 | 0.077 | 0.339 | 0.019 |
| Q259.17 | A | 0.397 | 0.206 | 0.001 | 0.005 | | 0.099 | 45.0 | 0.034 |
| MI369.A5 | A | 0.682 | 0.335 | 0.039 | 0.005 | 0.849 | 0.125 | 0.010 | 0.009 |
| 3718.v3.c11 | A | 0.809 | 0.188 | 0.105 | 0.044 | 0.272 | 0.075 | 0.046 | 0.004 |
| KNH1209.18 | A | 0.958 | | | 0.910 | 38.9 | 15.1 | | 20.8 |
| MB539.2B7 | A | 1.36 | 1.17 | 31.9 | 0.747 | | 0.114 | 1.19 | 0.012 |
| 0260.v5.c36 | A | 13.9 | | | | | 0.999 | | 0.018 |
| MB201.A1 | A | 14.7 | | | | 0.243 | 0.104 | 2.49 | 0.228 |
| Q769.h5 | A | | | | | 39.4 | 0.012 | 0.023 | 0.002 |
| Q769.d22 | A | | | | | | 0.011 | 0.151 | 0.003 |
| BB539.2B13 | A | | 1.02 | 0.194 | 0.005 | 6.34 | 0.109 | 0.077 | 0.010 |
| MS208.A1 | A | | 0.098 | 0.0007 | 0.003 | 3.05 | 0.147 | 0.231 | 0.015 |
| RW020.2 | A | | | | | | 0.140 | 5.17 | 0.016 |
| 3415.v1.c1 | A | | | | | 5.46 | 0.121 | | 0.075 |
| UG037.8 | A | | | | | 0.287 | 0.026 | 2.33 | 0.078 |
| QH209.14M.A2 | A | | | | | | | 0.004 | 0.089 |
| 0439.v5.c1 | A | | | | | | | 3.78 | 37.8 |
| 398-F1.F6.20 | A | | | | | 0.517 | | | |

#### AC, AD, ACD

| Virus ID | Clade | RHA1. V2.01 | PCT64-35M | CAP256-VRC26.08 | CAP256-VRC26.25 | CH01 | PG9 | PGT145 | PGDM 1400 |
|---|---|---|---|---|---|---|---|---|---|
| Q168.a2 | AD | 0.069 | 0.025 | 0.148 | 0.003 | 2.26 | 0.072 | 0.860 | 0.003 |
| 6545.v4.c1 | AC | 0.120 | 0.058 | 0.026 | 0.009 | 0.171 | 0.055 | 0.003 | 0.004 |
| 246-F3.C10.2 | AC | 0.179 | 0.029 | | 0.019 | 2.06 | 0.024 | 17.4 | 0.003 |
| 6540.v4.c1 | AC | 0.199 | 0.106 | 0.077 | 0.009 | 0.465 | 0.033 | 0.003 | 0.005 |
| 6095.v1.c10 | ACD | 12.5 | 0.984 | 0.0007 | 0.003 | | 0.279 | 0.002 | 0.011 |
| 3301.v1.c24 | AC | | 3.31 | 0.0003 | 0.004 | | 0.372 | 0.193 | 0.040 |
| Q461.e2 | AD | | | 0.502 | 0.046 | | 3.86 | 16.0 | 0.161 |
| 3589.v1.c4 | AC | | | | | 7.29 | 0.023 | | |
| 3468.v1.c12 | AD | | | | | 31.5 | 18.2 | | |
| 0815.v3.c3 | ACD | | | | | | | | |

## B

| Virus ID | Clade | RHA1. V2.01 | PCT64-35M | CAP256-VRC26.08 | CAP256-VRC26.25 | CH01 | PG9 | PGT145 | PGDM 1400 |
|---|---|---|---|---|---|---|---|---|---|
| JRCSF.JB | B | 0.015 | | | | 0.782 | 0.004 | 0.001 | 0.007 |
| WITO.33 | B | 0.019 | 2.57 | | | 0.013 | 0.009 | 0.001 | 0.004 |
| BX08.16 | B | 0.073 | | | 0.005 | 0.590 | 0.052 | 0.007 | 0.020 |
| X2278.C2.B6 | B | 0.133 | | | 0.005 | 0.052 | 0.010 | 0.002 | 0.012 |
| 5768.04 | B | 0.985 | | | | 0.825 | 0.243 | 0.020 | 0.043 |
| REJO.67 | B | 1.06 | | | | 0.620 | 0.018 | 0.0004 | 0.003 |
| AC10.29 | B | 1.17 | | | 0.111 | 16.3 | 0.058 | 0.006 | 0.102 |
| 45_01dG5 | B | 3.97 | | | 0.136 | 0.705 | 0.202 | 0.002 | 0.074 |
| 6535.3 | B | 4.16 | | | | 0.428 | 0.206 | | |
| HT593.1 | B | 6.70 | | | | | 0.587 | 0.058 | 0.456 |
| ADA.DG | B | 8.96 | | | | | 0.925 | 0.005 | 0.108 |
| 3988.25 | B | 11.5 | | | 0.048 | | 0.080 | 0.007 | 0.059 |
| HO86.8 | B | 43.6 | | | | 0.186 | 0.055 | 0.002 | 0.051 |
| THRO.18 | B | | | | | 9.32 | 12.4 | 0.006 | 0.070 |
| BaL.26 | B | | | | | | 0.108 | 1.41 | 0.077 |
| Bal.01 | B | | | | | | 0.143 | | 0.137 |
| SS1196.01 | B | | | | | | 0.303 | 19.4 | 0.220 |
| YU2.DG | B | | | | | | 1.81 | 0.035 | 0.224 |
| RHPA.7 | B | | | | | 8.33 | *10.7* | 0.028 | 0.416 |
| TRO.11 | B | | | | | | *27.9* | 0.029 | 0.616 |
| SC422.8 | B | | | | | | 0.957 | 0.027 | 0.737 |
| 7165.18 | B | | | | | 2.30 | | 0.056 | 1.24 |
| PVO.04 | B | | | 11.8 | 0.065 | 24.2 | 8.34 | 0.143 | 1.29 |
| JRFL.JB | B | | | | | | | 4.90 | |
| CAAN.A2 | B | | | | | | 8.93 | 5.53 | |
| CNE10 | B | | | | | | 0.264 | | |
| TRJO.58 | B | | | | | | 0.349 | | |
| HXB2.DG | B | | | | | | 0.685 | | |
| 6101.10 | B | | | | | | | | |
| 89.6.DG | B | | | | 49.9 | | | | |
| BG1168.01 | B | | | | | | | | |
| BL01.DG | B | | | | | | | | |
| BR07.DG | B | | | | | | | | |
| CNE12 | B | | | | | | | | |
| CNE14 | B | | | | | | | | |
| CNE4 | B | | | | | | | | |
| CNE57 | B | | | | | | | | |
| MN.3 | B | | | | | | | | |
| QH0515.01 | B | | | | | | | | |
| QH0692.42 | B | | | | | | | | |
| SF162.LS | B | | | | | | | | |

## AE

| Virus ID | Clade | RHA1. V2.01 | PCT64-35M | CAP256-VRC26.08 | CAP256-VRC26.25 | CH01 | PG9 | PGT145 | PGDM 1400 |
|---|---|---|---|---|---|---|---|---|---|
| C1080.c3 | AE | 0.001 | 0.011 | 0.0003 | 0.003 | 0.033 | 0.004 | 0.010 | 0.0007 |
| CM244.ec1 | AE | 0.004 | 0.003 | 0.002 | 0.043 | 0.016 | 0.004 | 0.0006 | 0.0004 |
| CNE5 | AE | 0.017 | 0.015 | 0.0003 | 0.002 | 0.104 | 0.010 | 0.002 | 0.0009 |
| CNE3 | AE | 0.041 | 0.013 | | | | 0.134 | 0.252 | 0.045 |
| C4118.09 | AE | 0.049 | 0.014 | 0.0003 | 0.002 | 0.254 | 0.052 | 0.002 | 0.003 |
| TH966.8 | AE | 0.092 | 0.019 | 0.0003 | 0.018 | 0.049 | 0.040 | 0.002 | 0.004 |
| CNE59 | AE | 0.102 | 0.018 | | | 2.61 | 0.070 | 0.018 | 0.002 |
| CNE55 | AE | 1.17 | 0.078 | 0.0003 | 0.002 | | 0.241 | 0.002 | 0.006 |
| CNE8 | AE | 8.55 | 0.780 | 2.46 | 0.019 | | 0.464 | 0.052 | 0.004 |
| 620345.c1 | AE | 23.5 | 0.234 | 0.001 | 0.013 | | 1.68 | 0.056 | 0.565 |
| R2184.c4 | AE | | 0.475 | 19.8 | 0.022 | | 0.472 | 0.050 | 0.007 |
| C2101.c1 | AE | | 22.7 | | | 1.62 | 0.050 | 0.005 | 0.010 |
| TH023.6 | AE | | 0.636 | 26.4 | | | 0.309 | 0.022 | 0.010 |
| R3265.c6 | AE | | | | | 0.608 | 0.091 | 0.011 | 0.015 |
| C3347.c11 | AE | | 10.2 | | | 4.44 | 0.081 | 0.004 | 0.024 |
| BJOX025000.01.1 | AE | | | | | | 0.154 | 1.44 | 0.055 |
| BJOX028000.10.3 | AE | | | | | | 0.686 | 4.00 | 0.061 |
| BJOX009000.02.4 | AE | | | | 18.8 | 10.1 | 1.00 | 0.099 | 0.177 |
| M02138 | AE | | | | 24.7 | 8.29 | 0.306 | | 0.290 |
| R1166.c1 | AE | | | | | 13.1 | 0.526 | 4.44 | 0.587 |
| BJOX010000.06.2 | AE | | | | | 1.57 | 0.268 | | 4.26 |
| CNE56 | AE | | | | | | | 1.63 | 9.02 |
| TH976.17 | AE | | | | 0.056 | | | | |

## AG

| Virus ID | Clade | RHA1. V2.01 | PCT64-35M | CAP256-VRC26.08 | CAP256-VRC26.25 | CH01 | PG9 | PGT145 | PGDM 1400 |
|---|---|---|---|---|---|---|---|---|---|
| T250-4 | AG | 0.010 | 0.009 | 0.0003 | 0.0008 | 0.042 | 0.003 | 0.0004 | 0.0006 |
| T33-7 | AG | 0.011 | 0.134 | | | 0.323 | 0.043 | 4.97 | 0.006 |
| 271-11 | AG | 0.018 | 0.134 | | | 1.44 | 0.100 | 35.7 | 0.008 |
| T255-34 | AG | 0.077 | 1.07 | | | 21.1 | 0.035 | | 0.012 |
| 235-47 | AG | 0.251 | 0.210 | 0.265 | 0.117 | 1.14 | 0.382 | 3.38 | 0.010 |
| 242-14 | AG | 0.869 | 0.078 | 0.0003 | 0.002 | | 0.040 | 0.033 | 0.004 |
| DJ263.8 | AG | 1.06 | 0.057 | 0.0007 | 0.006 | 2.99 | 0.109 | 0.018 | 0.003 |
| T278-50 | AG | 9.27 | | | 10.4 | 1.59 | 1.26 | 19.5 | 0.173 |
| T266-60 | AG | 13.5 | | 1.78 | 0.073 | 2.49 | 0.433 | 0.071 | 0.531 |
| T257-31 | AG | 30.3 | | 0.0003 | 0.002 | 0.768 | 0.029 | 3.49 | 0.003 |
| 928-28 | AG | | | 0.097 | 0.006 | 4.51 | 0.088 | | 0.062 |
| T253-11 | AG | | 6.69 | 49.5 | 0.022 | 3.06 | 0.213 | | 0.011 |
| 263-8 | AG | | | 0.031 | 0.014 | | 0.345 | 13.2 | 0.016 |
| T280-5 | AG | | | | 0.052 | 1.85 | 0.549 | 0.029 | 0.443 |
| 269-12 | AG | | | | 0.190 | 7.68 | 2.36 | 4.36 | 0.673 |
| T251-18 | AG | | | | | 2.23 | 27.5 | 0.318 | 4.50 |

## CD

| Virus ID | Clade | RHA1. V2.01 | PCT64-35M | CAP256-VRC26.08 | CAP256-VRC26.25 | CH01 | PG9 | PGT145 | PGDM 1400 |
|---|---|---|---|---|---|---|---|---|---|
| 3326.v4.c3 | CD | 0.162 | 0.285 | 0.0003 | 0.001 | | 0.034 | 0.009 | 0.003 |
| 3337.v2.c6 | CD | | | 0.0003 | 0.0005 | | | | |
| 3817.v2.c59 | CD | | | | | 0.092 | 0.011 | | |

## D

| Virus ID | Clade | RHA1. V2.01 | PCT64-35M | CAP256-VRC26.08 | CAP256-VRC26.25 | CH01 | PG9 | PGT145 | PGDM 1400 |
|---|---|---|---|---|---|---|---|---|---|
| 247-23 | D | 0.211 | 0.478 | 0.005 | 0.002 | | 0.177 | 0.017 | 0.004 |
| 3016.v5.c45 | D | 0.572 | 0.454 | 0.005 | 0.002 | | 0.433 | 25.4 | 4.30 |
| A07412M1.vrc12 | D | 9.22 | | 0.068 | 0.007 | | 3.84 | 0.003 | 0.057 |
| 191821.E6.1 | D | 48.6 | | | 0.006 | | 7.64 | 3.35 | 0.048 |
| 231965.c01 | D | | | | 1.32 | | 0.495 | 0.122 | 0.016 |
| 57128.vrc15 | D | | | | 0.129 | | 0.195 | 0.118 | 0.025 |
| A03349M1.vrc4a | D | | | 2.68 | 0.021 | | | 4.60 | 0.106 |
| UG024.2 | D | | 6.85 | 0.843 | 0.166 | | | | 0.592 |
| 6405.v4.c34 | D | | | | | | | | |
| NKU3006.ec1 | D | | | | | | | | |
| UG021.16 | D | | | | | | | | |

## G

| Virus ID | Clade | RHA1. V2.01 | PCT64-35M | CAP256-VRC26.08 | CAP256-VRC26.25 | CH01 | PG9 | PGT145 | PGDM 1400 |
|---|---|---|---|---|---|---|---|---|---|
| X1632.S2.B10 | G | 0.112 | 0.058 | 0.0003 | 0.003 | 5.25 | 0.157 | 0.006 | 0.005 |
| P0402.c2.11 | G | 2.14 | 2.35 | 0.0003 | 0.002 | | 0.413 | 0.007 | 0.017 |
| X2131.C1.B5 | G | 5.90 | | | 0.007 | 0.668 | 0.097 | 0.017 | 0.079 |
| P1981.C5.3 | G | | | 0.0003 | 0.003 | | 0.276 | 1.08 | 0.076 |
| X1193.c1 | G | | | 0.343 | 0.032 | | 0.158 | 0.015 | 0.187 |
| X1254.c3 | G | | | | | 18.5 | 0.050 | 1.27 | 37.5 |
| X2088.c9 | G | | | | | | | | |

## BC

| Virus ID | Clade | RHA1. V2.01 | PCT64-35M | CAP256-VRC26.08 | CAP256-VRC26.25 | CH01 | PG9 | PGT145 | PGDM 1400 |
|---|---|---|---|---|---|---|---|---|---|
| CH117.4 | BC | 0.005 | 0.008 | 0.0003 | 0.0004 | 1.16 | 0.004 | 0.012 | 0.001 |
| CH181.12 | BC | 0.009 | 0.002 | 0.0003 | 0.0005 | 0.494 | 0.014 | 0.0003 | 0.002 |
| CH070.1 | BC | 0.052 | 0.004 | 0.0008 | 0.004 | | 0.009 | 0.003 | 0.0006 |
| CNE21 | BC | 0.069 | 0.008 | 0.001 | 0.001 | | 0.067 | 0.002 | 0.002 |
| BJOX002000.03.2 | BC | 0.821 | 0.048 | 0.003 | 0.002 | 41.2 | 0.045 | 0.434 | 0.005 |
| CH038.12 | BC | 1.97 | | 0.009 | 0.003 | | 0.775 | 46.5 | |
| CNE20 | BC | 7.47 | | | 0.005 | 1.18 | 0.045 | 0.042 | 0.004 |
| CH119.10 | BC | 21.1 | | | 0.019 | 1.98 | 0.326 | 0.089 | 0.146 |
| CNE19 | BC | | 0.269 | 11.1 | 0.001 | 0.864 | 0.024 | 0.068 | 0.002 |
| CNE15 | BC | | 1.24 | | 1.27 | 0.356 | 0.009 | 2.60 | 0.004 |
| CNE7 | BC | | 0.887 | 0.003 | 0.005 | | 1.07 | 0.259 | 0.026 |
| CNE40 | BC | | | | 0.020 | | 1.44 | 0.214 | 0.505 |

## C

| Virus ID | Clade | RHA1. V2.01 | PCT64-35M | CAP256-VRC26.08 | CAP256-VRC26.25 | CH01 | PG9 | PGT145 | PGDM 1400 |
|---|---|---|---|---|---|---|---|---|---|
| CAP45.G3 | C | 0.006 | 0.004 | 0.0003 | 0.001 | 0.105 | 0.007 | 0.0008 | 0.002 |
| ZM233.6 | C | 0.009 | 0.089 | | 0.0009 | 0.074 | 0.013 | 0.011 | 0.001 |
| CNE58 | C | 0.045 | 0.035 | 0.0003 | 0.001 | 0.047 | 0.020 | 0.062 | 0.002 |
| BR025.9 | C | 0.051 | 0.009 | | 1.47 | 0.224 | 0.042 | 8.67 | 0.003 |
| 001428-2.42 | C | 0.057 | 0.244 | 30.9 | 3.01 | 0.074 | 0.004 | 0.0006 | 0.002 |
| 16055-2.3 | C | 0.077 | 0.020 | 0.0003 | 0.001 | | 0.017 | 0.008 | 0.002 |
| ZM176.66 | C | 0.078 | 0.018 | 0.0003 | 0.001 | 1.89 | 0.017 | 0.032 | 0.002 |
| 6785.v5.c14 | C | 0.107 | 0.311 | 0.0003 | 0.003 | 0.879 | 0.031 | 0.105 | 0.008 |
| 25925-2.22 | C | 0.156 | 0.027 | 0.0003 | 0.002 | 2.10 | 0.026 | 0.003 | 0.002 |
| SO18.18 | C | 0.205 | 0.010 | 0.0003 | 0.001 | 9.73 | 0.017 | 0.002 | 0.001 |
| CAP256.206.C9 | C | 0.225 | 0.028 | 0.0003 | 0.002 | 1.81 | 0.041 | 3.95 | 0.005 |
| 6838.v1.c35 | C | 0.271 | | 0.001 | 0.003 | 0.137 | 0.018 | 0.215 | 0.011 |
| DU123.06 | C | 0.281 | 0.383 | 0.015 | 0.018 | 0.515 | 0.119 | 0.024 | 0.014 |
| ZM249.1 | C | 0.324 | 0.143 | 0.0009 | 0.005 | | 0.047 | 1.99 | 0.013 |
| DU156.12 | C | 0.350 | 0.234 | 0.006 | 0.004 | 0.305 | 0.046 | 0.008 | 0.005 |
| 26191-2.48 | C | 0.404 | 0.100 | 0.001 | 0.002 | 3.00 | 0.113 | 0.053 | 0.006 |
| ZM53.12 | C | 0.589 | 0.070 | 0.0003 | 0.002 | | 0.038 | 0.154 | 0.003 |
| 286.36 | C | 0.701 | 0.110 | 0.0003 | 0.008 | 8.92 | 0.225 | 0.003 | 0.011 |
| 25710-2.43 | C | 0.856 | 0.033 | 0.0003 | 0.002 | 2.50 | 0.042 | 0.002 | 0.003 |
| DU151.02 | C | 1.05 | 0.866 | 0.006 | 0.002 | 0.671 | 0.024 | 0.006 | 0.007 |
| CE703010217.B6 | C | 2.19 | 3.94 | 5.82 | 0.010 | 0.351 | 0.009 | 0.051 | 0.005 |
| TV1.29 | C | 2.28 | 0.372 | | 0.493 | 2.05 | 0.015 | 3.34 | 0.002 |
| TZBD.02 | C | 4.01 | | 0.0009 | 0.009 | 20.0 | 0.236 | 0.035 | 0.011 |
| 3873.v1.c24 | C | 6.10 | 0.224 | 0.0003 | 0.004 | | | 0.002 | 0.006 |
| CNE53 | C | 6.18 | | | | 0.236 | 0.094 | 0.019 | 0.043 |
| CAP210.E8 | C | 11.2 | 0.970 | 0.0003 | 0.002 | 15.7 | 0.160 | | 0.015 |
| MW965.26 | C | 12.6 | | 0.003 | 0.063 | | 2.10 | 0.147 | 0.065 |
| ZM106.9 | C | 24.7 | 0.479 | 0.010 | 0.006 | | 0.370 | 0.016 | 0.011 |
| TZA125.17 | C | 40.8 | | 0.018 | 0.006 | | 0.186 | 0.115 | 0.039 |
| ZM197.7 | C | 43.1 | 3.17 | 0.002 | 0.008 | | 0.616 | 0.523 | 0.071 |
| ZM215.8 | C | | | | | | 0.033 | | 0.001 |
| 0013095-2.11 | C | | | 0.021 | 0.044 | 0.981 | 0.026 | 16.6 | 0.003 |
| 0921.v2.c14 | C | | 0.799 | 0.0003 | 0.001 | 0.382 | 0.007 | 0.011 | 0.004 |
| 6322.v4.c1 | C | | 1.20 | 0.0003 | 0.004 | | | 0.073 | 0.011 |
| 0077.v1.c16 | C | | 3.51 | 0.001 | 0.005 | 42.9 | 0.104 | 0.194 | 0.016 |
| ZM109.4 | C | | | | 0.006 | | 0.270 | 0.012 | 0.016 |
| ZM55.28a | C | | 2.72 | 0.005 | 0.017 | | 0.706 | 0.687 | 0.031 |
| 00836-2.5 | C | | | | | | | 0.887 | 0.044 |
| 3168.v4.c10 | C | | | | | 1.18 | 0.282 | 0.028 | 0.172 |
| CNE31 | C | | | | | | 27.5 | 0.068 | 0.199 |
| 96ZM651.02 | C | | | 2.63 | 0.839 | | | 7.33 | 0.209 |
| 16936-2.21 | C | | 0.205 | 0.250 | 0.003 | | | 5.14 | 0.222 |
| Ce1176.A3 | C | | | 31.9 | 0.065 | 0.181 | 0.022 | | 0.369 |
| 25711-2.4 | C | | | | | 2.78 | 0.734 | 7.46 | 0.514 |
| 288.38 | C | | | | | 8.83 | 4.31 | 2.61 | 0.727 |
| ZA012.29 | C | | | | 2.25 | | 14.6 | 1.26 | 1.31 |
| DU172.17 | C | | | | | 7.70 | 0.291 | | 1.94 |
| CAP244.D3 | C | | | 46.0 | | 0.655 | 0.073 | 12.0 | 4.92 |
| 16845-2.22 | C | | | | | 5.74 | 2.86 | 46.7 | |
| 6644.v2.c33 | C | | | | | | 0.044 | | |
| DU422.01 | C | | | 1.12 | 0.018 | | 1.02 | | |
| ZM214.15 | C | | | 3.23 | 0.011 | | | | |
| 3637.v5.c3 | C | | | | | | | | |
| 426c | C | | | | | | | | |
| 6471.v1.c16 | C | | | | | | | | |
| 6631.v3.c10 | C | | | | | | | | |
| CNE30 | C | | | | | | | | |
| ZM135.10a | C | | | | | | | | |

### Supplemental Figure S25

## A

RHA1.V2.01 Signatures

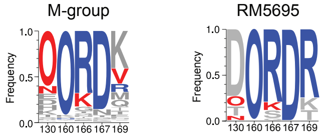

Sensitivity signature
Resistance signature
Neutral
O: PNGS

## B

RHA1.V2.01 & other V2 apex bNAb signatures

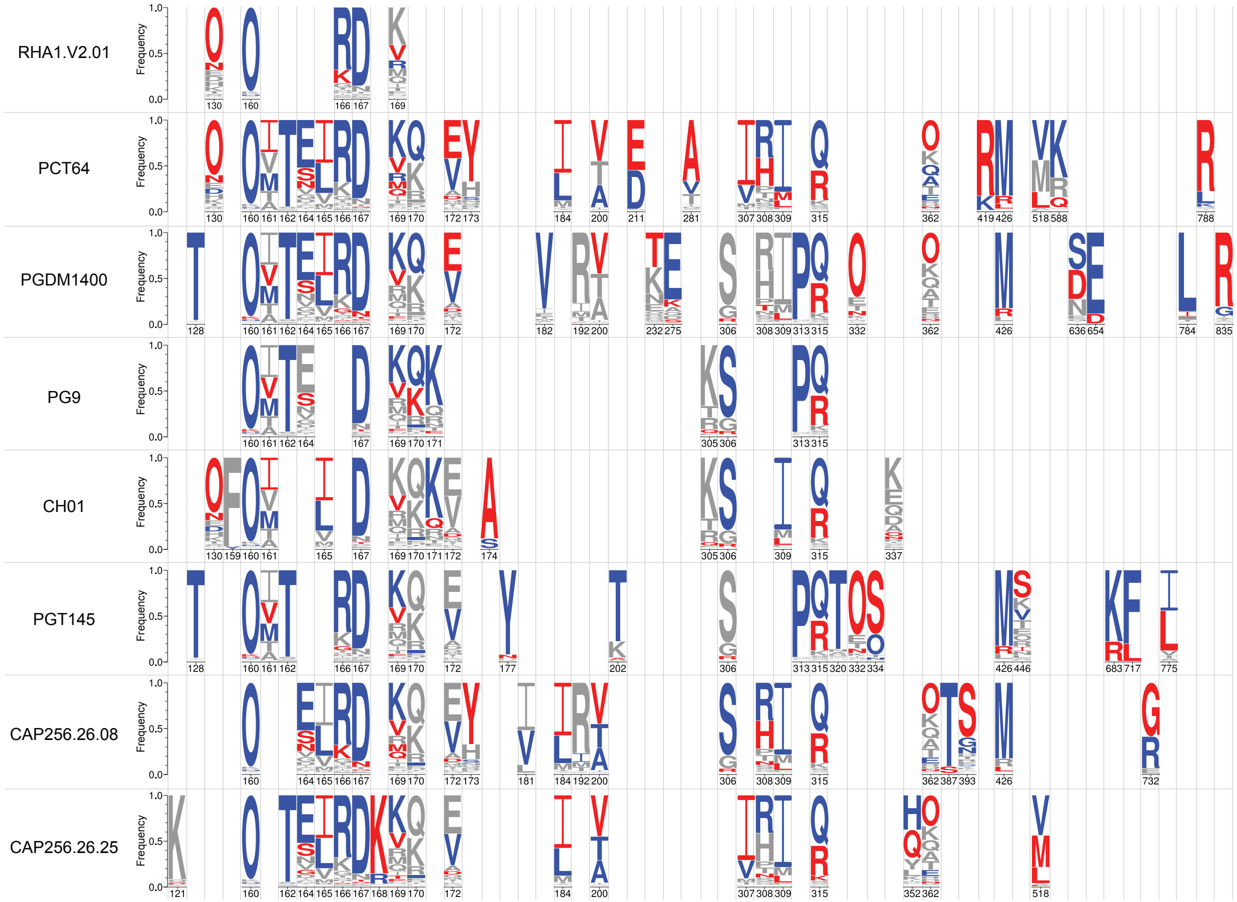

### Supplemental Figure S26

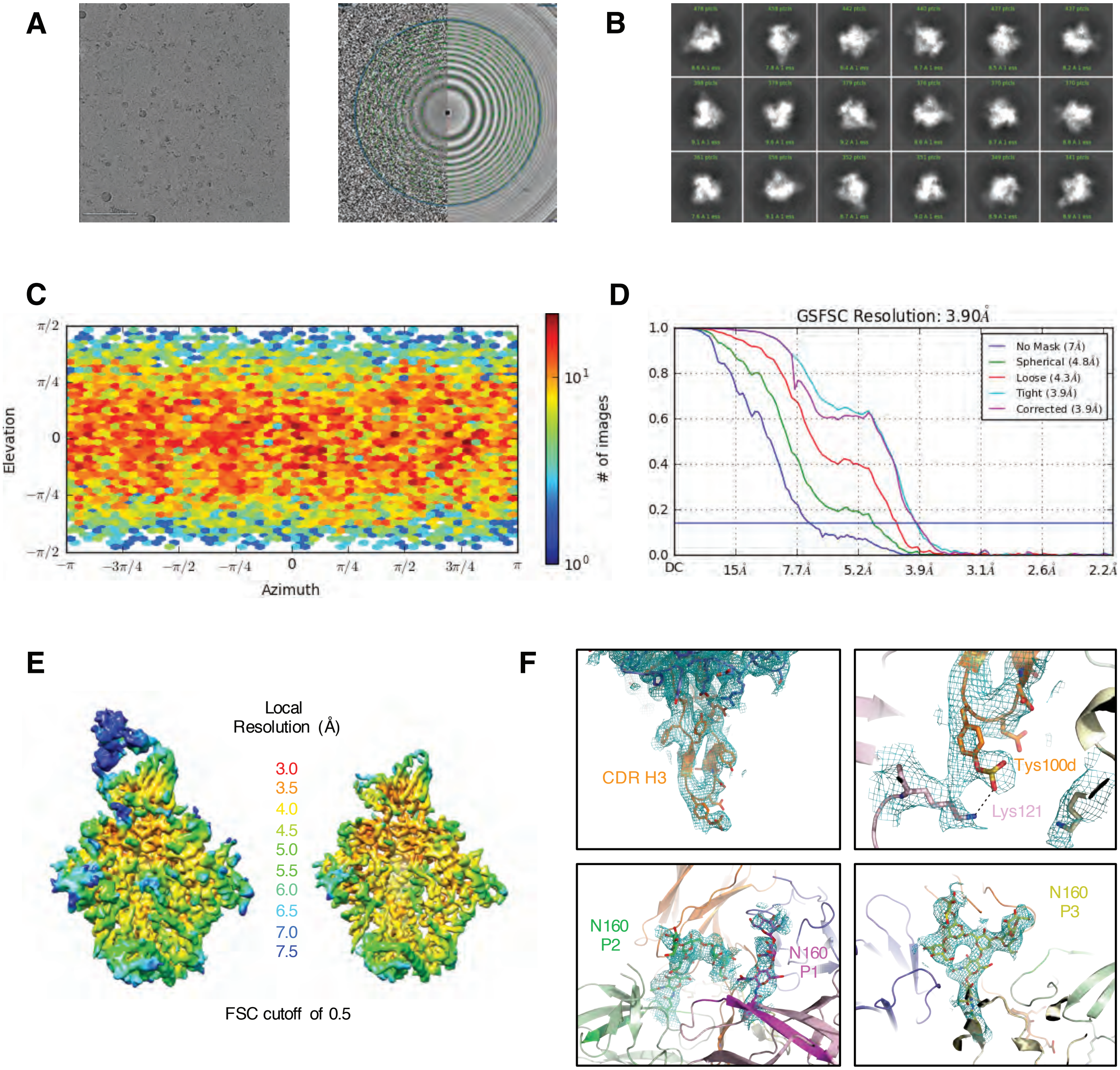

**Supplemental Figure S27**

**Table S1. Infection details, clinical parameters and neutralizing antibody responses in rhesus macaques**

| Animal ID | Inoculum | Env Subtype | Route | Depletion of CD8$^{+}$ cells | Weeks of Followup | VL Peak (copies/ml) | VL Setpoint (copies/ml) | Clinical AIDS | Peak autologous NAb Titer | bNAb Induction | bNAb Specificities |
|---|---|---|---|---|---|---|---|---|---|---|---|
| RM6069 | SHIV CH505 | C | IV | Yes* | 40 | 19,400,000 | 988,110 | Yes | 1396 | No | |
| RM6070 | SHIV CH505 | C | IV | Yes* | 36 | 62,600,000 | 736,483 | Yes | 7837 | Yes | V2 Apex |
| RM6072 | SHIV CH505 | C | IV | Yes* | 184 | 15,700,000 | 20,405 | No | 3544 | No | |
| RM6697 | SHIV CH505 | C | IV | No | 112 | 812,436 | 1,638 | No | 918 | No | |
| RM6698 | SHIV CH505 | C | IV | No | 112 | 943,718 | 526 | No | 277 | No | |
| RM6699 | SHIV CH505 | C | IV | No | 112 | 1,155,342 | 506 | No | 422 | No | |
| RM6701 | SHIV CH505 | C | IV | No | 112 | 302,350 | <250 | No | <20 | No | |
| RM6702 | SHIV CH505 | C | IV | No | 96 | 1,007,432 | <250 | No | 514 | No | |
| RM6703 | SHIV CH505 | C | IV | No | 112 | 5,257,396 | 2,546 | No | 787 | No | |
| RM5695 | SHIV CH505# | C | IV | No | 65 | 4,791,344 | 12,110 | Yes | 573 | Yes | V2 Apex |
| RM6163 | SHIV CH848 | C | IV | Yes* | 176 | 24,870,060 | 38,409 | No | 544 | Yes | V3 glycan |
| RM6167 | SHIV CH848 | C | IV | Yes* | 136 | 116,025,900 | 117,295 | Yes | 558 | Yes | V3 glycan |
| RM6700 | SHIV CH848 | C | IV | No | 112 | 1,271,384 | 13,760 | No | 418 | No | |
| RM6713 | SHIV CH848 | C | IV | No | 89 | 17,682,010 | 45,008 | Yes | 732 | No | |
| RM6714 | SHIV CH848 | C | IV | No | 112 | 832,110 | 15,114 | No | 655 | No | |
| RM6720 | SHIV CH848 | C | IV | No | 112 | 1,323,796 | <250 | No | 187 | No | |
| RM40591 | SHIV CAP256SU | C | IV | No | 129 | 2,540,090 | 24,676 | No | 3067 | Yes | V2 Apex |
| RM42056 | SHIV CAP256SU | C | IV | Yes^ | 88 | 55,885,850 | 8,130 | No | 347 | Yes | V2 Apex |
| RM6727 | SHIV CAP256SU | C | IV | Yes^ | 60 | 5,282,264 | 67,392 | Yes | 665 | Yes | Unknown |
| RM43037 | SHIV CAP256SU | C | IV | Yes^ | 88 | 11,662,030 | 10,280 | No | 492 | No | |
| RM40547 | SHIV CAP256SU | C | IV | No | 88 | 825,090 | <250 | No | <20 | No | |
| RM40624 | SHIV CAP256SU | C | IV | No | 88 | 670,330 | <250 | No | <20 | No | |

### Plasma from RMs 6069 (wk10 & wk20), 6070 (wk10 & wk20), 6072 (wk4, wk10 & wk20)

* Anti-CD8α antibody (MT807R1, NHP Reagent Resource), 50 mg/kg by slow intravenous push administered on day 0 with respect to SHIV inoculation.

^ Anti-CD8β antibody (CD8beta255R1, NHP Reagent Resource), 25mg/kg by slow intravenous push administered on day -7 with respect to SHIV inoculation.

#### Table S2. Autologous and heterologous neutralization by plasma from SHIV infected rhesus macaques

| | | | | Tier 1A | | | Tier 2 | | | | | | | | | | | | | | | | | | |
|---|---|---|---|---|---|---|---|---|---|---|---|---|---|---|---|---|---|---|---|---|---|---|---|---|---|
| | | | | MN.3 | MW965.26 | TH023.6 | TRO11 | 25710 | CNE8 | X2278 | BJOX 010000 | X1632 | Ce 1176 | 246F3 | CH119 | Ce0217 | CNE55 | Q23 | T250 | WITO | ZM233 | CAP256 SU | CH505 | CH848 | BG505 332N |
| | Animal ID | Wpi* | MLV# | B | C | AE | B | C | AE | B | AE | G | C | AC | AE | C | AE | A | A | B | C | C | C | C | A |
| CH505 | RM5695 | 0 | <20 | <20^ | 240 | 68 | <20 | <20 | <20 | <20 | <20 | <20 | <20 | <20 | <20 | <20 | <20 | <20 | <20 | <20 | <20 | <20 | <20 | <20 | <20 |
| | | 16 | <20 | 231 | 49,677 | 20,691 | <20 | 58 | <20 | <20 | <20 | 27 | <20 | <20 | <20 | <20 | <20 | 144 | 46 | <20 | 45 | <20 | 46 | <20 | <20 |
| | | 24 | <20 | 231 | 36,483 | 4,462 | <20 | <20 | <20 | <20 | <20 | 24 | <20 | <20 | <20 | <20 | <20 | 168 | 36 | 55 | 234 | 32 | 573 | 38 | 28 |
| | | 32 | <20 | 270 | 39,573 | 5,659 | <20 | <20 | <20 | <20 | <20 | 26 | <20 | 47 | <20 | <20 | <20 | 4,507 | 93 | 49 | 110 | 36 | 393 | 43 | 29 |
| | | 48 | <20 | 371 | 54,675 | 6,028 | <20 | 28 | <20 | <20 | <20 | 195 | <20 | 140 | <20 | <20 | <20 | 6,784 | 1,941 | 172 | 123 | 81 | 361 | 30 | 92 |
| | | 56 | <20 | 532 | 49,505 | 5,342 | 72 | 78 | 179 | 28 | 22 | 465 | 20 | 109 | <20 | 26 | 20 | 5,051 | 1,977 | 138 | 413 | 65 | 421 | 127 | 139 |
| | RM6070 | 0 | <20 | <20 | <20 | <20 | <20 | <20 | <20 | <20 | <20 | <20 | <20 | <20 | <20 | <20 | <20 | <20 | <20 | <20 | <20 | <20 | <20 | <20 | <20 |
| | | 8 | <20 | 128 | 1,522 | 735 | <20 | <20 | <20 | <20 | <20 | <20 | <20 | <20 | <20 | <20 | <20 | <20 | 25 | 35 | <20 | <20 | 108 | <20 | <20 |
| | | 16 | <20 | 1,458 | 25,484 | 6,472 | <20 | <20 | <20 | <20 | <20 | <20 | <20 | <20 | <20 | <20 | <20 | 42 | 28 | 46 | 48 | <20 | 2,556 | <20 | <20 |
| | | 24 | <20 | 1,882 | 69,252 | 10,212 | <20 | <20 | <20 | <20 | <20 | <20 | <20 | <20 | <20 | <20 | <20 | 182 | 48 | 61 | 73 | <20 | 4,172 | <20 | <20 |
| | | 36 | <20 | 1,830 | 43,403 | 1,724 | <20 | <20 | <20 | <20 | <20 | 40 | <20 | 27 | <20 | <20 | <20 | 3,003 | 59 | <20 | 65 | <20 | 2,178 | 21 | <20 |
| | RM6072 | 0 | <20 | <20 | <20 | <20 | <20 | <20 | <20 | <20 | <20 | <20 | <20 | <20 | <20 | <20 | <20 | <20 | <20 | <20 | <20 | <20 | <20 | <20 | <20 |
| | | 88 | <20 | 189 | 30,220 | 3,118 | <20 | <20 | <20 | <20 | <20 | <20 | <20 | <20 | <20 | <20 | <20 | <20 | <20 | <20 | <20 | <20 | 747 | <20 | <20 |
| | | 104 | <20 | 145 | 19,881 | 3,987 | <20 | <20 | <20 | <20 | <20 | <20 | <20 | <20 | <20 | <20 | <20 | <20 | <20 | <20 | <20 | <20 | 256 | <20 | <20 |
| | | 144 | <20 | 182 | 15,718 | 1,397 | <20 | <20 | <20 | <20 | <20 | <20 | <20 | <20 | <20 | <20 | <20 | <20 | <20 | <20 | <20 | <20 | 342 | <20 | <20 |
| | RM6069 | 0 | <20 | <20 | <20 | <20 | <20 | <20 | <20 | <20 | <20 | <20 | <20 | <20 | <20 | <20 | <20 | <20 | <20 | <20 | <20 | <20 | <20 | <20 | <20 |
| | | 40 | <20 | 115 | 38,037 | 1,579 | <20 | <20 | <20 | <20 | <20 | 24 | <20 | <20 | <20 | <20 | <20 | <20 | <20 | <20 | <20 | <20 | 168 | <20 | <20 |
| | RM6697 | 0 | <20 | <20 | <20 | <20 | <20 | <20 | <20 | <20 | <20 | <20 | <20 | <20 | <20 | <20 | <20 | <20 | <20 | <20 | <20 | <20 | <20 | <20 | <20 |
| | | 80 | <20 | 593 | 14,736 | 2,226 | <20 | <20 | <20 | <20 | <20 | <20 | <20 | <20 | <20 | <20 | <20 | 121 | 150 | <20 | 36 | <20 | 741 | <20 | <20 |
| | | 96 | <20 | 622 | 16,271 | 4,492 | <20 | <20 | 66 | <20 | <20 | <20 | <20 | <20 | <20 | <20 | <20 | 128 | <20 | <20 | <20 | <20 | 318 | <20 | <20 |
| | RM6703 | 0 | <20 | <20 | <20 | <20 | <20 | <20 | <20 | <20 | <20 | <20 | <20 | <20 | <20 | <20 | <20 | <20 | <20 | <20 | <20 | <20 | <20 | <20 | <20 |
| | | 64 | <20 | 160 | 5,373 | 416 | <20 | <20 | <20 | <20 | <20 | <20 | <20 | <20 | <20 | <20 | <20 | <20 | <20 | <20 | <20 | <20 | 493 | <20 | <20 |
| | | 80 | <20 | 359 | 8,865 | 734 | <20 | <20 | <20 | <20 | <20 | <20 | <20 | <20 | <20 | <20 | <20 | <20 | <20 | <20 | <20 | <20 | 787 | <20 | <20 |
| | | 88 | <20 | 108 | 4,677 | 257 | <20 | <20 | <20 | <20 | <20 | <20 | <20 | <20 | <20 | <20 | <20 | <20 | <20 | <20 | <20 | <20 | 221 | <20 | <20 |
| | RM6698 | 0 | <20 | <20 | <20 | <20 | <20 | <20 | <20 | <20 | <20 | <20 | <20 | <20 | <20 | <20 | <20 | <20 | <20 | <20 | <20 | <20 | <20 | <20 | <20 |
| | | 56 | <20 | 294 | 9,872 | 930 | 23 | <20 | <20 | <20 | <20 | <20 | <20 | <20 | <20 | <20 | <20 | 61 | <20 | <20 | 25 | <20 | 141 | <20 | <20 |
| | | 96 | <20 | 109 | 10,469 | 632 | <20 | <20 | 75 | <20 | <20 | <20 | <20 | <20 | <20 | <20 | <20 | 126 | <20 | <20 | <20 | <20 | 246 | <20 | <20 |
| | RM6699 | 0 | <20 | <20 | <20 | <20 | <20 | <20 | <20 | <20 | <20 | <20 | <20 | <20 | <20 | <20 | <20 | <20 | <20 | <20 | <20 | <20 | <20 | <20 | <20 |
| | | 56 | <20 | 491 | 6,481 | 1,437 | <20 | <20 | <20 | <20 | <20 | <20 | <20 | <20 | <20 | <20 | <20 | 24 | <20 | <20 | <20 | <20 | 422 | <20 | <20 |
| | | 72 | <20 | 515 | 9,124 | 1,156 | <20 | <20 | <20 | <20 | <20 | <20 | <20 | <20 | <20 | <20 | <20 | <20 | <20 | <20 | <20 | <20 | 267 | <20 | <20 |
| | | 80 | <20 | 433 | 8,489 | 1,706 | <20 | <20 | <20 | <20 | <20 | <20 | <20 | <20 | <20 | <20 | <20 | 26 | <20 | <20 | 24 | 32 | 261 | <20 | <20 |
| | RM6701 | 0 | <20 | <20 | <20 | <20 | <20 | <20 | <20 | <20 | <20 | <20 | <20 | <20 | <20 | <20 | <20 | <20 | <20 | <20 | <20 | <20 | <20 | <20 | <20 |
| | | 56 | <20 | 277 | 13,077 | 1,364 | <20 | <20 | <20 | <20 | <20 | <20 | <20 | <20 | <20 | <20 | <20 | <20 | <20 | <20 | <20 | <20 | <20 | <20 | <20 |
| | | 72 | <20 | 268 | 13,068 | 1,504 | <20 | <20 | <20 | <20 | <20 | <20 | <20 | <20 | <20 | <20 | <20 | <20 | <20 | <20 | <20 | <20 | <20 | <20 | <20 |
| | | 80 | <20 | 391 | 23,855 | 3,172 | <20 | 38 | <20 | <20 | <20 | <20 | <20 | <20 | <20 | <20 | <20 | 80 | <20 | <20 | 44 | <20 | <20 | <20 | <20 |
| | RM6702 | 0 | <20 | <20 | <20 | <20 | <20 | <20 | <20 | <20 | <20 | <20 | <20 | <20 | <20 | <20 | <20 | <20 | <20 | <20 | <20 | <20 | <20 | <20 | <20 |
| | | 64 | <20 | 2,259 | 5,924 | 794 | <20 | <20 | 63 | <20 | <20 | <20 | <20 | <20 | <20 | <20 | <20 | <20 | <20 | <20 | 69 | <20 | 286 | <20 | <20 |
| | | 80 | <20 | 2,037 | 9,183 | 1,272 | <20 | <20 | <20 | <20 | <20 | <20 | <20 | <20 | <20 | <20 | <20 | 108 | <20 | <20 | <20 | <20 | 215 | <20 | <20 |
| CH848 | RM6163 | 0 | <20 | <20 | <20 | <20 | <20 | <20 | <20 | <20 | <20 | <20 | <20 | <20 | <20 | <20 | <20 | <20 | <20 | <20 | <20 | <20 | <20 | <20 | <20 |
| | | 64 | <20 | 1,481 | 37,313 | 406 | <20 | <20 | <20 | <20 | <20 | <20 | <20 | <20 | <20 | <20 | <20 | <20 | <20 | <20 | <20 | <20 | <20 | 83 | <20 |
| | | 80 | <20 | 1,348 | 14,057 | 515 | <20 | 139 | <20 | <20 | <20 | <20 | 101 | <20 | <20 | <20 | <20 | <20 | <20 | <20 | <20 | <20 | <20 | 483 | <20 |
| | | 88 | <20 | 1,478 | 17,819 | 401 | <20 | 61 | 152 | <20 | <20 | <20 | 37 | <20 | <20 | <20 | <20 | <20 | <20 | <20 | <20 | <20 | <20 | 834 | <20 |
| | | 104 | <20 | 577 | 5,555 | 24 | <20 | 53 | <20 | <20 | <20 | <20 | 25 | <20 | <20 | <20 | <20 | <20 | <20 | <20 | <20 | <20 | <20 | 233 | <20 |
| | | 112 | <20 | 372 | 4,219 | 111 | <20 | 142 | 70 | 25 | <20 | <20 | 45 | <20 | <20 | <20 | <20 | <20 | 40 | <20 | 34 | <20 | <20 | 161 | <20 |
| | | 128 | <20 | 485 | 7,776 | 120 | <20 | 1,286 | <20 | <20 | <20 | <20 | 551 | <20 | <20 | <20 | <20 | <20 | 28 | <20 | <20 | <20 | <20 | 538 | <20 |
| | | 136 | <20 | 376 | 6,631 | 117 | <20 | 334 | <20 | <20 | <20 | <20 | 256 | <20 | <20 | <20 | <20 | <20 | <20 | <20 | <20 | <20 | <20 | 561 | <20 |
| | | 144 | <20 | 442 | 10,529 | 392 | <20 | 231 | 373 | <20 | <20 | <20 | 162 | <20 | <20 | <20 | <20 | <20 | <20 | <20 | <20 | <20 | <20 | 339 | <20 |
| | | 152 | <20 | 411 | 8,787 | 337 | <20 | 121 | 79 | <20 | <20 | <20 | 88 | <20 | <20 | <20 | <20 | <20 | <20 | <20 | <20 | <20 | <20 | 413 | <20 |
| | RM6167 | 0 | <20 | <20 | <20 | <20 | <20 | <20 | <20 | <20 | <20 | <20 | <20 | <20 | <20 | <20 | <20 | <20 | <20 | <20 | <20 | <20 | <20 | <20 | <20 |
| | | 64 | <20 | 21,533 | 138,427 | 1,957 | <20 | 44 | <20 | <20 | <20 | <20 | 35 | <20 | <20 | <20 | <20 | <20 | 26 | <20 | 32 | 32 | <20 | 176 | <20 |
| | | 80 | <20 | 25,687 | 88,417 | 3,272 | 97 | 314 | 25 | 38 | <20 | 49 | 78 | <20 | 44 | <20 | 21 | 39 | 215 | 42 | 135 | 153 | <20 | 383 | 62 |
| | | 88 | <20 | 13,394 | 34,686 | 1,426 | 38 | 39 | 192 | <20 | <20 | 37 | 33 | <20 | 19 | <20 | <20 | 57 | 43 | 31 | 86 | 66 | 26 | 313 | 28 |
| | | 104 | <20 | 13,110 | 24,027 | 29 | 48 | 59 | <20 | 24 | <20 | 62 | 59 | 51 | 117 | 44 | 22 | 36 | 126 | 23 | 90 | 58 | 21 | 101 | 23 |
| | | 112 | <20 | 6,892 | 38,256 | 402 | 135 | 97 | 128 | 25 | <20 | 35 | 51 | <20 | 56 | 25 | <20 | 74 | 176 | <20 | 93 | 142 | <20 | 150 | 23 |
| | | 128 | <20 | 18,129 | 64,893 | 1,238 | 134 | 159 | <20 | 39 | 27 | 42 | 84 | 20 | 84 | 21 | <20 | 113 | 209 | 25 | 47 | 135 | <20 | 74 | 39 |
| | RM6700 | 0 | <20 | <20 | <20 | <20 | <20 | <20 | <20 | <20 | <20 | <20 | <20 | <20 | <20 | <20 | <20 | <20 | <20 | <20 | <20 | <20 | <20 | <20 | <20 |
| | | 40 | <20 | 1,071 | 39,032 | 1,299 | <20 | <20 | <20 | <20 | <20 | <20 | <20 | <20 | <20 | <20 | <20 | <20 | <20 | <20 | <20 | <20 | <20 | 389 | <20 |
| | | 72 | <20 | 1,423 | 48,567 | 1,654 | <20 | 52 | <20 | <20 | <20 | <20 | <20 | <20 | <20 | <20 | <20 | <20 | <20 | <20 | <20 | <20 | <20 | 418 | <20 |
| | | 80 | <20 | 1,246 | 34,941 | 1,792 | <20 | 47 | <20 | <20 | <20 | <20 | <20 | <20 | <20 | <20 | <20 | <20 | <20 | <20 | <20 | <20 | <20 | 339 | <20 |
| | | 88 | <20 | 1,152 | 53,591 | 2,775 | <20 | 73 | <20 | 23 | <20 | 30 | <20 | <20 | <20 | <20 | <20 | <20 | <20 | <20 | 25 | <20 | <20 | 338 | <20 |
| | | 95 | <20 | 1,016 | 51,520 | 1,429 | <20 | 33 | <20 | <20 | <20 | 84 | <20 | <20 | <20 | <20 | <20 | <20 | <20 | <20 | 22 | <20 | <20 | 308 | <20 |
| | RM6713 | 0 | <20 | <20 | <20 | <20 | <20 | <20 | <20 | <20 | <20 | <20 | <20 | <20 | <20 | <20 | <20 | <20 | <20 | <20 | <20 | <20 | <20 | <20 | <20 |
| | | 40 | <20 | 8,375 | 64,350 | 1,057 | <20 | 23 | <20 | <20 | <20 | <20 | <20 | <20 | <20 | <20 | <20 | 35 | <20 | <20 | 31 | <20 | <20 | 75 | <20 |
| | | 72 | <20 | 6,406 | 43,159 | 1,644 | <20 | <20 | <20 | <20 | <20 | <20 | <20 | <20 | <20 | <20 | <20 | <20 | <20 | <20 | 38 | <20 | <20 | 732 | <20 |
| | | 80 | <20 | 7,800 | 47,081 | 973 | <20 | <20 | <20 | <20 | <20 | 23 | <20 | <20 | <20 | <20 | <20 | <20 | <20 | <20 | 31 | <20 | <20 | 447 | <20 |
| | RM6714 | 0 | <20 | <20 | <20 | <20 | <20 | <20 | <20 | <20 | <20 | <20 | <20 | <20 | <20 | <20 | <20 | <20 | <20 | <20 | <20 | <20 | <20 | <20 | <20 |
| | | 40 | <20 | 254 | 22,447 | 279 | <20 | 24 | <20 | <20 | <20 | <20 | <20 | <20 | <20 | <20 | <20 | <20 | <20 | <20 | 26 | <20 | <20 | 95 | <20 |
| | | 72 | <20 | 111 | 15,858 | 245 | <20 | 20 | <20 | <20 | <20 | <20 | <20 | <20 | <20 | <20 | <20 | <20 | <20 | <20 | <20 | <20 | <20 | 337 | <20 |
| | | 80 | <20 | 109 | 25,907 | 509 | <20 | 21 | <20 | <20 | <20 | <20 | <20 | <20 | <20 | <20 | <20 | <20 | <20 | <20 | <20 | <20 | <20 | 174 | <20 |
| | RM6720 | 0 | <20 | <20 | <20 | <20 | <20 | <20 | <20 | <20 | <20 | <20 | <20 | <20 | <20 | <20 | <20 | <20 | <20 | <20 | <20 | <20 | <20 | <20 | <20 |
| | | 40 | <20 | 1,211 | 11,203 | 533 | <20 | <20 | <20 | <20 | <20 | <20 | <20 | <20 | <20 | <20 | <20 | <20 | <20 | <20 | <20 | <20 | <20 | <20 | <20 |
| | | 72 | <20 | 1,944 | 19,301 | 853 | <20 | <20 | 42 | <20 | <20 | <20 | <20 | <20 | <20 | <20 | <20 | <20 | <20 | <20 | <20 | <20 | <20 | 185 | <20 |
| | | 80 | <20 | 2,838 | 26,695 | 1,149 | <20 | <20 | <20 | <20 | <20 | <20 | <20 | <20 | <20 | <20 | <20 | <20 | <20 | <20 | <20 | <20 | <20 | 113 | <20 |
| CAP 256 SU | RM40591 | 0 | <20 | <20 | <20 | <20 | <20 | <20 | <20 | <20 | <20 | <20 | <20 | <20 | <20 | <20 | <20 | <20 | <20 | <20 | <20 | <20 | <20 | <20 | <20 |
| | | 72 | <20 | 2,167 | 14,176 | 977 | <20 | <20 | <20 | <20 | <20 | <20 | <20 | <20 | <20 | <20 | <20 | <20 | <20 | <20 | <20 | 2,230 | <20 | <20 | <20 |
| | | 88 | <20 | 2,970 | 31,857 | 976 | <20 | <20 | 30 | <20 | 25 | <20 | <20 | <20 | <20 | 66 | <20 | <20 | <20 | <20 | 32 | 3,067 | <20 | <20 | <20 |
| | | 96 | <20 | 3,374 | 29,806 | 6,627 | <20 | 22 | <20 | <20 | 197 | <20 | <20 | <20 | <20 | 179 | <20 | <20 | <20 | <20 | 207 | 2,648 | <20 | <20 | <20 |
| | | 104 | <20 | 5,467 | 54,915 | 8,237 | 20 | 28 | 457 | 45 | 101 | <20 | <20 | <20 | <20 | 339 | <20 | 119 | <20 | <20 | 506 | 2,344 | <20 | <20 | <20 |
| | | 112 | <20 | 5,382 | 59,952 | 6,798 | 24 | 58 | 31 | 49 | 59 | 36 | <20 | 25 | <20 | 214 | <20 | 318 | 90 | <20 | 206 | 1,692 | <20 | <20 | <20 |
| | | 120 | <20 | 4,045 | 48,733 | 3,433 | <20 | <20 | 324 | 31 | 48 | <20 | <20 | <20 | <20 | 275 | 28 | 66 | 32 | <20 | 154 | 1,069 | <20 | <20 | <20 |
| | | 128 | <20 | 7,435 | 61,501 | 5,411 | <20 | <20 | 608 | 154 | 44 | <20 | <20 | <20 | <20 | 138 | 33 | 87 | <20 | <20 | 209 | 1,663 | <20 | <20 | <20 |
| | RM42056 | 0 | <20 | <20 | <20 | <20 | <20 | <20 | <20 | <20 | <20 | <20 | <20 | <20 | <20 | <20 | <20 | <20 | <20 | <20 | <20 | <20 | <20 | <20 | <20 |
| | | 48 | <20 | 1,083 | 2,462 | 65 | <20 | <20 | <20 | <20 | <20 | <20 | <20 | <20 | <20 | <20 | <20 | <20 | 106 | <20 | <20 | 123 | <20 | <20 | <20 |
| | | 56 | <20 | 1,260 | 6,378 | 221 | <20 | <20 | <20 | <20 | <20 | <20 | <20 | <20 | <20 | <20 | <20 | 36 | 248 | <20 | 20 | 347 | <20 | <20 | <20 |
| | | 64 | <20 | 753 | 2,846 | 367 | <20 | <20 | <20 | <20 | <20 | <20 | <20 | <20 | <20 | <20 | <20 | 62 | 160 | <20 | 24 | 149 | <20 | <20 | <20 |
| | | 72 | <20 | 811 | 3,778 | 286 | <20 | <20 | <20 | <20 | <20 | <20 | <20 | <20 | <20 | <20 | <20 | 81 | 92 | <20 | 26 | 159 | <20 | <20 | <20 |
| | | 80 | <20 | 488 | 2,850 | 287 | <20 | <20 | <20 | <20 | <20 | 20 | <20 | <20 | <20 | <20 | <20 | 141 | 134 | <20 | 37 | 149 | <20 | <20 | <20 |
| | | 88 | <20 | 521 | 2,757 | 362 | <20 | <20 | <20 | <20 | <20 | 21 | <20 | <20 | <20 | <20 | <20 | 190 | 159 | <20 | 42 | 217 | <20 | <20 | <20 |
| | | 104 | <20 | 678 | 2,378 | 414 | 34 | 23 | 42 | <20 | 21 | 57 | <20 | <20 | <20 | <20 | <20 | 574 | 115 | <20 | 38 | 140 | <20 | <20 | <20 |
| | RM6727 | 0 | <20 | <20 | <20 | <20 | <20 | <20 | <20 | <20 | <20 | <20 | <20 | <20 | <20 | <20 | <20 | <20 | <20 | <20 | <20 | <20 | <20 | <20 | <20 |
| | | 48 | <20 | 2,471 | 27,322 | 642 | 21 | <20 | <20 | <20 | <20 | 24 | <20 | <20 | <20 | <20 | <20 | <20 | 27 | <20 | 165 | 227 | <20 | <20 | <20 |
| | | 56 | <20 | 1,511 | 43,630 | 1,082 | 26 | 30 | <20 | 41 | <20 | <20 | <20 | <20 | <20 | <20 | <20 | <20 | 111 | <20 | 146 | 178 | <20 | <20 | <20 |
| | | 60 | <20 | 736 | 33,300 | 2,341 | 35 | 57 | 52 | 25 | <20 | 53 | <20 | <20 | 26 | <20 | <20 | 42 | 74 | <20 | 100 | 116 | <20 | <20 | <20 |
| | RM43037 | 0 | <20 | <20 | <20 | <20 | <20 | <20 | <20 | <20 | <20 | <20 | <20 | <20 | <20 | <20 | <20 | <20 | <20 | <20 | <20 | <20 | <20 | <20 | <20 |
| | | 48 | <20 | 2,242 | 5,669 | 510 | <20 | <20 | <20 | <20 | <20 | <20 | <20 | <20 | <20 | <20 | <20 | <20 | <20 | <20 | <20 | 424 | <20 | <20 | <20 |
| | | 80 | <20 | 925 | 4,845 | 555 | <20 | <20 | <20 | <20 | <20 | <20 | <20 | <20 | <20 | <20 | <20 | <20 | <20 | <20 | <20 | 104 | <20 | <20 | <20 |
| | | 104 | <20 | 2,990 | 8,711 | 2,687 | <20 | <20 | <20 | <20 | <20 | <20 | <20 | <20 | <20 | <20 | <20 | <20 | <20 | <20 | <20 | 100 | <20 | <20 | <20 |
| | RM40547 | 0 | <20 | <20 | <20 | <20 | <20 | <20 | <20 | <20 | <20 | <20 | <20 | <20 | <20 | <20 | <20 | <20 | <20 | <20 | <20 | <20 | <20 | <20 | <20 |
| | | 88 | <20 | 138 | 2,282 | 153 | <20 | <20 | <20 | <20 | <20 | <20 | <20 | <20 | <20 | <20 | <20 | <20 | <20 | <20 | <20 | <20 | <20 | <20 | <20 |
| | RM40624 | 0 | <20 | <20 | <20 | <20 | <20 | <20 | <20 | <20 | <20 | <20 | <20 | <20 | <20 | <20 | <20 | <20 | <20 | <20 | <20 | <20 | <20 | <20 | <20 |
| | | 88 | <20 | 136 | 1,583 | 51 | <20 | <20 | <20 | <20 | <20 | <20 | <20 | <20 | <20 | <20 | <20 | <20 | <20 | <20 | <20 | <20 | <20 | <20 | <20 |

* Wpi - weeks post-SHIV infection.

### Murine leukemia virus Env-pseudotyped virus; other HIV-1 Envs in this row are listed.

^ Reciprocal plasma neutralization titer (ID50 20-100, green; 100-1000, yellow; 1000-10000, orange; >10000, red).

Bold black rectangular boxes highlight results from seven rhesus macaques with broadly neutralizing antibodies.

#### Table S3. Neutralization breadth of DH650.8 against 118-strain panel

| Virus ID | Clade* | Titer (μg/ml) DH650.8 IC50 |
|---|---|---|
| 6535.3 | B | >50 |
| QH0692.42 | B | >50 |
| SC422661.8 | B | >50 |
| PVO.4 | B | >50 |
| TRO.11 | B | >50 |
| AC10.0.29 | B | >50 |
| RHPA4259.7 | B | >50 |
| THRO4156.18 | B | >50 |
| REJO4541.67 | B | >50 |
| TRJO4551.58 | B | >50 |
| WITO4160.33 | B | >50 |
| CAAN5342.A2 | B | >50 |
| WEAU_d15_410_787 | B (T/F) | >50 |
| 1006_11_C3_1601 | B (T/F) | >50 |
| 1054_07_TC4_1499 | B (T/F) | >50 |
| 1056_10_TA11_1826 | B (T/F) | >50 |
| 1012_11_TC21_3257 | B (T/F) | >50 |
| 6240_08_TA5_4622 | B (T/F) | >50 |
| 6244_13_B5_4576 | B (T/F) | >50 |
| 62357_14_D3_4589 | B (T/F) | >50 |
| SC05_8C11_2344 | B (T/F) | >50 |
| Du156.12 | C | >50 |
| Du172.17 | C | >50 |
| Du422.1 | C | >50 |
| ZM197M.PB7 | C | >50 |
| ZM214M.PL15 | C | >50 |
| ZM233M.PB6 | C | >50 |
| ZM249M.PL1 | C | >50 |
| ZM53M.PB12 | C | >50 |
| ZM109F.PB4 | C | >50 |
| ZM135M.PL10a | C | >50 |
| CAP45.2.00.G3 | C | >50 |
| CAP210.2.00.E8 | C | >50 |
| HIV-001428-2.42 | C | >50 |
| HIV-0013095-2.11 | C | >50 |
| HIV-16055-2.3 | C | >50 |
| HIV-16845-2.22 | C | >50 |
| Ce1086_B2 | C (T/F) | >50 |
| Ce0393_C3 | C (T/F) | >50 |
| Ce1176_A3 | C (T/F) | >50 |
| Ce2010_F5 | C (T/F) | >50 |
| Ce0682_E4 | C (T/F) | >50 |
| Ce1172_H1 | C (T/F) | >50 |
| Ce2060_G9 | C (T/F) | >50 |
| Ce703010054_2A2 | C (T/F) | >50 |
| BF1266.431a | C (T/F) | >50 |
| 246F C1G | C (T/F) | >50 |
| 249M B10 | C (T/F) | >50 |
| ZM247v1(Rev-) | C (T/F) | >50 |
| 7030102001E5(Rev-) | C (T/F) | >50 |
| 1394C9G1(Rev-) | C (T/F) | >50 |
| Ce704809221_1B3 | C (T/F) | >50 |
| CNE19 | BC | >50 |
| CNE20 | BC | >50 |
| CNE21 | BC | >50 |
| CNE17 | BC | >50 |
| CNE30 | BC | >50 |
| CNE52 | BC | >50 |
| CNE53 | BC | >50 |
| CNE58 | BC | >50 |
| MS208.A1 | A | >50 |
| Q23.17 | A | >50 |
| Q461.e2 | A | >50 |
| Q769.d22 | A | >50 |
| Q259.d2.17 | A | >50 |
| Q842.d12 | A | >50 |
| 0260.v5.c36 | A | >50 |
| 3415.v1.c1 | A | >50 |
| 0330.v4.c3 | A | >50 |
| 191955_A11 | A (T/F) | >50 |
| 191084 B7-19 | A (T/F) | >50 |
| 9004SS_A3_4 | A (T/F) | >50 |
| T257-31 | CRF02_AG | >50 |
| 928-28 | CRF02_AG | >50 |
| 263-8 | CRF02_AG | >50 |
| T250-4 | CRF02_AG | >50 |
| T251-18 | CRF02_AG | >50 |
| T278-50 | CRF02_AG | >50 |
| T255-34 | CRF02_AG | >50 |
| 211-9 | CRF02_AG | >50 |
| 235-47 | CRF02_AG | >50 |
| 620345.c01 | CRF01_AE | >50 |
| CNE8 | CRF01_AE | >50 |
| C1080.c03 | CRF01_AE | >50 |
| R2184.c04 | CRF01_AE | >50 |
| R1166.c01 | CRF01_AE | >50 |
| R3265.c06 | CRF01_AE | Not tested |
| C2101.c01 | CRF01_AE | >50 |
| C3347.c11 | CRF01_AE | >50 |
| C4118.c09 | CRF01_AE | >50 |
| CNE5 | CRF01_AE | >50 |
| BJOX009000.02.4 | CRF01_AE | >50 |
| BJOX015000.11.5 | CRF01_AE (T/F) | >50 |
| BJOX010000.06.2 | CRF01_AE (T/F) | >50 |
| BJOX025000.01.1 | CRF01_AE (T/F) | >50 |
| BJOX028000.10.3 | CRF01_AE (T/F) | >50 |
| X1193_c1 | G | >50 |
| P0402_c2_11 | G | >50 |
| X1254_c3 | G | >50 |
| X2088_c9 | G | >50 |
| X2131_C1_B5 | G | >50 |
| P1981_C5_3 | G | >50 |
| X1632_S2_B10 | G | >50 |
| 3016.v5.c45 | D | >50 |
| A07412M1.vrc12 | D | >50 |
| 231965.c01 | D | >50 |
| 231966.c02 | D | >50 |
| 3817.v2.c59 | CD | >50 |
| 6480.v4.c25 | CD | >50 |
| 6952.v1.c20 | CD | >50 |
| 6811.v7.c18 | CD | >50 |
| 89-F1_2_25 | CD | >50 |
| 3301.v1.c24 | AC | >50 |
| 6041.v3.c23 | AC | >50 |
| 6540.v4.c1 | AC | >50 |
| 6545.v4.c1 | AC | >50 |
| 0815.v3.c3 | ACD | >50 |
| 3103.v3.c10 | ACD | >50 |

* (T/F): Transmitted / Founder Virus

**Table S4. Diffraction and Refinement Statistics**

| | DH650 + GP120 |
|---|---|
| **Diffraction data** | |
| Space group | $P2_12_12$ |
| Cell dimensions | |
| Lengths (Å) a, b, c | 138.5 122.1 53.4 |
| Angles (°) α, β, γ | 90.0 90.0 90.0 |
| $d_{min}$ (Å) | 2.80 (2.97-2.80)[a] |
| $R_{merge}$[b] | 0.13 (1.52) |
| $CC_{1/2}$ | 99.7 (64.4) |
| Avg $I/\sigma_I$ | 13.99 (1.28) |
| Completeness (%) | 99.8 (99.9) |
| Average redundancy | 5.5 (5.6) |
| **Refinement** | |
| Data range (Å) | 49.9-2.80 |
| Reflections | 20,060 |
| $R/R_{free}$[c] | 0.23/0.28 |
| RMS deviations | |
| Bond (Å) | 0.008 |
| Angles (°) | 1.02 |
| Avg B-factor (Å$^2$) | |
| Protein | 54.2 |
| Water | 33.32 |
| Number of Atoms | |
| Protein | 5,565 |
| Water | 21 |

[a] Values in parentheses are for the outermost shell of data.

[b] $R_{merge} = \sum |I_i >< I >| / \sum I_i$, where $I_i$ is the intensity of the $i^{th}$ observation and $\langle I \rangle$ is the mean intensity. Sums are taken over all reflections.

[c] $R = \sum ||F_o| > |F_c|| / \sum |F_o|$. $R_{free}$ is calculated for a 5% subset of the data

#### Table S5. RM5695 personalized Ig germline repertoire

##### Heavy

| IGHV | GenBank Accession | IGHJ | GenBank Accession |
|---|---|---|---|
| IGHV1-AAU-U*01 | | JH-1*01 | |
| IGHV1-AAU-U*02 | | JH-2P*01 | |
| IGHV1-ABW-S*01_S9171 | XXXXXXXX | JH-3*01 | |
| IGHV1-AEP*01 | | JH-4*01 | |
| IGHV1-AFS*01 | | JH-5-1*01_S8786 | XXXXXXXX |
| IGHV1-AGS*02_S4535 | XXXXXXXX | JH-5-2*01 | |
| IGHV1-AGS*02_S6849 | XXXXXXXX | JH-6*01 | |
| IGHV2-ABG-S*01 | | | |
| IGHV2-ABG-S*02_S2032 | XXXXXXXX | | |
| IGHV2-ABU-S*01_S6524 | XXXXXXXX | | |
| IGHV2-AEY-S*01 | | | |
| IGHV2-AEY-S*01_S8940 | XXXXXXXX | | |
| IGHV3-AAB*01 | | | |
| IGHV3-AAB*02 | | | |
| IGHV3-ABA*01 | | | |
| IGHV3-ABA*01_S4331 | XXXXXXXX | | |
| IGHV3-ABJ*01_S1922 | XXXXXXXX | | |
| IGHV3-ABK*01_S3779 | XXXXXXXX | | |
| IGHV3-ABY-S*01 | | | |
| IGHV3-ABY-S*01_S8824 | XXXXXXXX | | |
| IGHV3-ABY-S*03_S0204 | XXXXXXXX | | |
| IGHV3-ACA*01 | | | |
| IGHV3-ACN*01_S7808 | XXXXXXXX | | |
| IGHV3-ACZ*03 | | | |
| IGHV3-ADF*01_S4250 | XXXXXXXX | | |
| IGHV3-ADL-S*01_S4190 | XXXXXXXX | | |
| IGHV3-ADO*01 | | | |
| IGHV3-ADR*01 | | | |
| IGHV3-ADR*03_S0556 | XXXXXXXX | | |
| IGHV3-ADR*03_S5688 | XXXXXXXX | | |
| IGHV3-ADX*01_S5510 | XXXXXXXX | | |
| IGHV3-AEH*01_S5714 | XXXXXXXX | | |
| IGHV3-AEH*01_S6567 | XXXXXXXX | | |
| IGHV3-AET*01_S5460 | XXXXXXXX | | |
| IGHV3-AEW-S*03 | | | |
| IGHV3-AEW-S*03_S2686 | XXXXXXXX | | |
| IGHV3-AFC*01_S2394 | XXXXXXXX | | |
| IGHV3-AFE*01_S4619 | XXXXXXXX | | |
| IGHV3-AFR*01_S0643 | XXXXXXXX | | |
| IGHV3-AFW*01_S5271 | XXXXXXXX | | |
| IGHV3-AFY-S*02 | | | |
| IGHV3-AFY-S*06 | | | |
| IGHV3-AGQ-S*02_S3249 | XXXXXXXX | | |
| IGHV4-ABB-S*01_S0511 | XXXXXXXX | | |
| IGHV4-ABB-S*01_S8200 | XXXXXXXX | | |
| IGHV4-ADD*01_S6003 | XXXXXXXX | | |
| IGHV4-ADD*01_S9501 | XXXXXXXX | | |
| IGHV4-ADG-U*01_S1296 | XXXXXXXX | | |
| IGHV4-AEB*01_S7180 | XXXXXXXX | | |
| IGHV4-AEX-S*01 | | | |
| IGHV4-AEX-S*01_S0860 | XXXXXXXX | | |
| IGHV4-AEX-S*01_S4729 | XXXXXXXX | | |
| IGHV4-AFB-S*01 | | | |
| IGHV4-AFB-S*01_S8393 | XXXXXXXX | | |
| IGHV4-AFI*01_S1365 | XXXXXXXX | | |
| IGHV4-AFL-U*01 | | | |
| IGHV4-AFQ-U*01 | | | |
| IGHV4-AFQ-U*01_S2532 | XXXXXXXX | | |
| IGHV4-AFU-U*01 | | | |
| IGHV4-AFU-U*01_S9450 | XXXXXXXX | | |
| IGHV4-AGL*01_S3100 | XXXXXXXX | | |
| IGHV4-AGR*01 | | | |
| IGHV4-AGU*01 | | | |
| IGHV4-AGU*01_S1118 | XXXXXXXX | | |
| IGHV4-AGU*01_S2780 | XXXXXXXX | | |
| IGHV5-ABI*01_S2502 | XXXXXXXX | | |
| IGHV5-ABI*02_S3096 | XXXXXXXX | | |
| IGHV7-ABS*01_S4281 | XXXXXXXX | | |
| IGHV7-AGO-S*01 | | | |
| IGHV7-AGO-S*03_S8272 | XXXXXXXX | | |

##### Lambda

| IGLV | GenBank Accession | IGLJ | GenBank Accession |
|---|---|---|---|
| IGLV1-ABB*01_S4519 | XXXXXXXX | JL1*01 | |
| IGLV1-ABB*01_S6715 | XXXXXXXX | JL2*01 | |
| IGLV1-ABB*01_S8237 | XXXXXXXX | JL3*01 | |
| IGLV1-ABN*02_S2133 | XXXXXXXX | JL6*01_S7212 | XXXXXXXX |
| IGLV1-ABN*02_S8168 | XXXXXXXX | JL7*02 | |
| IGLV1-ABS*01_S5111 | XXXXXXXX | | |
| IGLV1-ACN*02 | | | |
| IGLV1-ACR*01 | | | |
| IGLV1-ACV*01_S2830 | XXXXXXXX | | |
| IGLV1-ACV*01_S4108 | XXXXXXXX | | |
| IGLV1-ACW*02 | | | |
| IGLV1-ADA*01 | | | |
| IGLV1-ADL*01 | | | |
| IGLV1-ADU*01_S4142 | XXXXXXXX | | |
| IGLV1-ADU*01_S8066 | XXXXXXXX | | |
| IGLV2-ABE*01_S2946 | XXXXXXXX | | |
| IGLV2-ABE*01_S9768 | XXXXXXXX | | |
| IGLV2-ABI*01_S7244 | XXXXXXXX | | |
| IGLV2-ABI*01_S8018 | XXXXXXXX | | |
| IGLV2-ABJ*01_S9052 | XXXXXXXX | | |
| IGLV2-ABU*01_S3471 | XXXXXXXX | | |
| IGLV2-ABX*01_S5497 | XXXXXXXX | | |
| IGLV2-ACE*01_S4286 | XXXXXXXX | | |
| IGLV2-ACE*01_S7743 | XXXXXXXX | | |
| IGLV2-ADC*01 | | | |
| IGLV2-ADC*01_S5785 | XXXXXXXX | | |
| IGLV2-ADK*01 | | | |
| IGLV3-AAB*01_S3980 | XXXXXXXX | | |
| IGLV3-AAB*01_S5697 | XXXXXXXX | | |
| IGLV3-AAI*01_S0428 | XXXXXXXX | | |
| IGLV3-AAV*01 | | | |
| IGLV3-AAV*04 | | | |
| IGLV3-ACB*01_S7967 | XXXXXXXX | | |
| IGLV3-ADT*01_S0731 | XXXXXXXX | | |
| IGLV3-AEB-X*02 | | | |
| IGLV3-AEB-X*03_S6646 | XXXXXXXX | | |
| IGLV3-AEC-X*01 | | | |
| IGLV3-AED-X*01_S4855 | XXXXXXXX | | |
| IGLV3-AED-X*01_S6916 | XXXXXXXX | | |
| IGLV3-AED-X*06_S3217 | XXXXXXXX | | |
| IGLV3-AED-X*06_S4240 | XXXXXXXX | | |
| IGLV4-ACF*01_S1875 | XXXXXXXX | | |
| IGLV4-ACF*02 | | | |
| IGLV5-AAM*01 | | | |
| IGLV5-AAP*01 | | | |
| IGLV5-AAP*01_S8201 | XXXXXXXX | | |
| IGLV5-AAX*01 | | | |
| IGLV5-ABG*01_S1697 | XXXXXXXX | | |
| IGLV5-ABL*01 | | | |
| IGLV5-ABT*01_S4414 | XXXXXXXX | | |
| IGLV5-ACY*01 | | | |
| IGLV5-ADQ*01_S3524 | XXXXXXXX | | |
| IGLV6-AAC*01_S4139 | XXXXXXXX | | |
| IGLV6-ADD*01_S2475 | XXXXXXXX | | |
| IGLV6-ADW*01 | | | |
| IGLV7-AAK*01 | | | |
| IGLV7-ABO*01 | | | |
| IGLV7-ADB*01 | | | |
| IGLV8-ABK*01 | | | |
| IGLV10-AAO*01 | | | |
| IGLV11-AAY*01 | | | |

**Table S6. Cryo-EM Data Collection and Refinement Statistics**

| | RHA1.V2.01 in complex with HIV-1 Env BG505 DS-SOSIP |
|---|---|
| **EMDB ID** | xxxx |
| **PDB ID** | xxxx |
| Data Collection | |
| Microscope | FEI Titan Krios |
| Voltage (kV) | 300 |
| Electron dose ($e^-$/Å$^2$) | 71.06 |
| Detector | Gatan K2 Summit |
| Pixel Size (Å) | 1.07 |
| Defocus Range ($\mu$m) | -0.1 to -4.1 |
| Magnification | 22500 |
| Reconstruction | |
| Software | cryoSparcV2.12 |
| Particles | 21,480 |
| Symmetry | C1 |
| Box size (pix) | 288 |
| Resolution (Å) ($FSC_{0.143}$) | 3.9 |
| Refinement | |
| Software | Phenix 1.18 |
| Protein residues | 1954 |
| Chimera CC | 0.84 |
| EMRinger Score | 1.44 |
| R.m.s. deviations | |
| Bond lengths (Å) | 0.003 |
| Bond angles (°) | 0.756 |
| Validation | |
| Molprobity score | 1.69 |
| Clash score | 4.41 |
| Favored rotamers (%) | 99.8 |
| Ramachandran | |
| Favored regions (%) | 92.4 |
| Disallowed regions (%) | 0.10 |
